# Multi-omic QTL mapping in early developmental tissues reveals phenotypic and temporal complexity of regulatory variants underlying GWAS loci

**DOI:** 10.1101/2024.04.10.588874

**Authors:** Timothy D. Arthur, Jennifer P. Nguyen, Agnieszka D’Antonio-Chronowska, Jeffrey Jaureguy, Nayara Silva, Benjamin Henson, iPSCORE Consortium, Athanasia D. Panopoulos, Juan Carlos Izpisua Belmonte, Matteo D’Antonio, Graham McVicker, Kelly A. Frazer

**Affiliations:** Biomedical Sciences Program, University of California, San Diego, La Jolla, CA, USA; Department of Biomedical Informatics, University of California, San Diego, La Jolla, CA, USA; Bioinformatics and Systems Biology Graduate Program, University of California, San Diego, La Jolla, CA, USA; Center for Epigenomics, University of California San Diego, School of Medicine, La Jolla, CA, USA; Integrative Biology Laboratory, Salk Institute for Biological Studies, La Jolla, CA, USA; Department of Pediatrics, University of California, San Diego, La Jolla, CA, USA; Institute of Genomic Medicine, University of California San Diego, La Jolla, CA, USA; Board of Governors Regenerative Medicine Institute, Cedars-Sinai Medical Center, Los Angeles, CA, USA; Department of Biomedical Sciences, Cedars-Sinai Medical Center, Los Angeles, CA, USA; Altos Labs, Inc., San Diego, CA, USA

## Abstract

Most GWAS loci are presumed to affect gene regulation, however, only ∼43% colocalize with expression quantitative trait loci (eQTLs). To address this colocalization gap, we identify eQTLs, chromatin accessibility QTLs (caQTLs), and histone acetylation QTLs (haQTLs) using molecular samples from three early developmental (EDev) tissues. Through colocalization, we annotate 586 GWAS loci for 17 traits by QTL complexity, QTL phenotype, and QTL temporal specificity. We show that GWAS loci are highly enriched for colocalization with complex QTL modules that affect multiple elements (genes and/or peaks). We also demonstrate that caQTLs and haQTLs capture regulatory variations not associated with eQTLs and explain ∼49% of the functionally annotated GWAS loci. Additionally, we show that EDev-unique QTLs are strongly depleted for colocalizing with GWAS loci. By conducting one of the largest multi-omic QTL studies to date, we demonstrate that many GWAS loci exhibit phenotypic complexity and therefore, are missed by traditional eQTL analyses.

## Introduction

Over 90% of genome-wide association study (GWAS) loci are in non-coding regions of the genome, and the causal variants at these loci are presumed to modulate the expression of genes. Expression quantitative trait loci (eQTL) analyses have been employed to interpret the regulatory function of GWAS signals^1^, however, only ∼43% of GWAS loci colocalize with eQTLs identified in post-mortem adult tissues^1^. Various hypotheses have been proposed to explain this colocalization gap including that regulatory variants may not be captured by eQTLs identified in adult bulk tissues because they are only active in context-specific conditions (e.g., early fetal development^2^). Additionally, it has been proposed that current eQTL study sample sizes are underpowered and biased toward discovering common variants with large effects on gene expression, which overlap promoters and affect genes under low evolutionary constraint^3^. In contrast, GWAS loci often overlap distal regulatory elements with small effects on the expression of genes under strong selective constraint^3^. These proposed hypotheses for the existing colocalization gap could be examined by focusing on cells representing early developmental time points, and by mapping QTLs for other types of molecular assays that specifically capture the function of distal regulatory elements such as enhancers.

The iPSC Omics Resource (iPSCORE)^2,4–17^ was developed to study the association between regulatory variation and molecular phenotypes in early developmental (EDev)-like tissues. iPSCORE is composed of early embryonic-like induced pluripotent stem cells (iPSCs) from hundreds of individuals with whole genome sequencing data^6,7,17^, as well as fetal-like iPSC-derived cardiovascular progenitor cells (CVPCs)^4,5,11,12^ and iPSC-derived pancreatic progenitor cells (PPCs)^2^. We have previously shown that the iPSCs, CVPCs, and PPCs are suitable surrogate models to identify eQTLs active during embryonic and fetal-like stages because they exhibit EDev-like molecular properties^2,4,6^. Moreover, we have shown the utility of combining the iPSCORE EDev-like and the GTEx adult expression datasets^1^ to functionally annotate regulatory variants in a temporal-specific manner^2,4^.

Molecular assays for transposase-accessible chromatin (ATAC-seq) and chromatin immunoprecipitation for H3K27 acetylation (H3K27ac ChIP-seq) capture both proximal and distal regulatory elements that modulate gene expression^18,19^. H3K27ac peaks primarily mark active enhancers^20,21^, whereas ATAC-seq identifies all open chromatin regions, which may contain both activating and repressive regulatory elements as well as insulators^18^. Integration of chromatin accessibility QTL (caQTL) and histone acetylation QTL (haQTL) mapping with eQTL analyses in the iPSCORE Collection could be particularly useful for discovering variants that affect different types of regulatory elements and enhance the annotation of GWAS variants, particularly those that are not associated with eQTLs.

To better understand the utility of using multi-omic QTLs from early developmental (EDev)-like tissues to functionally annotate GWAS loci, we analyzed RNA-seq, ATAC-seq, and H3K27ac ChIP-seq samples generated from iPSCs, CVPCs, and PPCs derived from genetically diverse individuals in the iPSCORE Collection. We performed QTL analyses and identified 79,081 QTLs (30,265 eQTLs, 36,559 caQTLs, 12,257 haQTLs), and defined complex QTL modules associated with multiple molecular elements (genes and/or peaks), and 7,614 EDev-unique QTLs (i.e., not present in adult QTL databases). We performed colocalization with GWAS loci from 17 developmental and adult traits and diseases and identified 586 loci that colocalized with a QTL. Of these 586 loci, 47.8% (n=280) only colocalized with caQTLs and/or haQTLs, marking a 1.9-fold increase compared to those that colocalized with the traditional eQTL approach. Additionally, we show that EDev-unique QTLs are strongly depleted for colocalizing with GWAS loci compared to adult-shared QTLs. Finally, to demonstrate the phenotypic and temporal complexity of EDev QTLs underlying adult disease-associated loci, we highlight two examples, including a CVPC QTL module that colocalized with a birth weight GWAS locus, and an EDev-unique PPC caQTL that colocalized with a body mass index GWAS locus.

In summary, our study is the first to show that integrative eQTL, caQTL, and haQTL analyses explain a large proportion of GWAS loci that are not functionally annotated using eQTLs alone. We are also the first to identify a large set of EDev-unique QTLs and show that, while a small fraction of GWAS loci colocalized only with EDev-unique regulatory variation, most of the QTLs that colocalized with GWAS loci were also active in adult tissues.

## Results

### Overview of Molecular Datasets

We analyzed 1,261 molecular samples including whole genome sequencing (WGS) and three molecular data types generated from three different iPSCORE early developmental (EDev)-like tissues (Figure 1) from 221 ethnically diverse iPSCORE subjects (170 Europeans, 4 Africans, 34 East Asians, 6 South Asians, and 7 Admixed Americans, Figure S1, Table S1). Specifically, we examined RNA-seq (Table S2), ATAC-seq (Table S3), and H3K27ac ChIP-seq (Table S4), from 220 induced pluripotent stem cell (iPSC) lines, 181 iPSC-derived cardiovascular progenitor cells (CVPCs), and 109 iPSC-derived pancreatic progenitor cells (PPCs). Of the 1,261 molecular samples, 400 are newly released in this study (Figure 1; indicated by the asterisks), and 861 have been previously published^2,6,7,12,17^.

**Figure 1.**
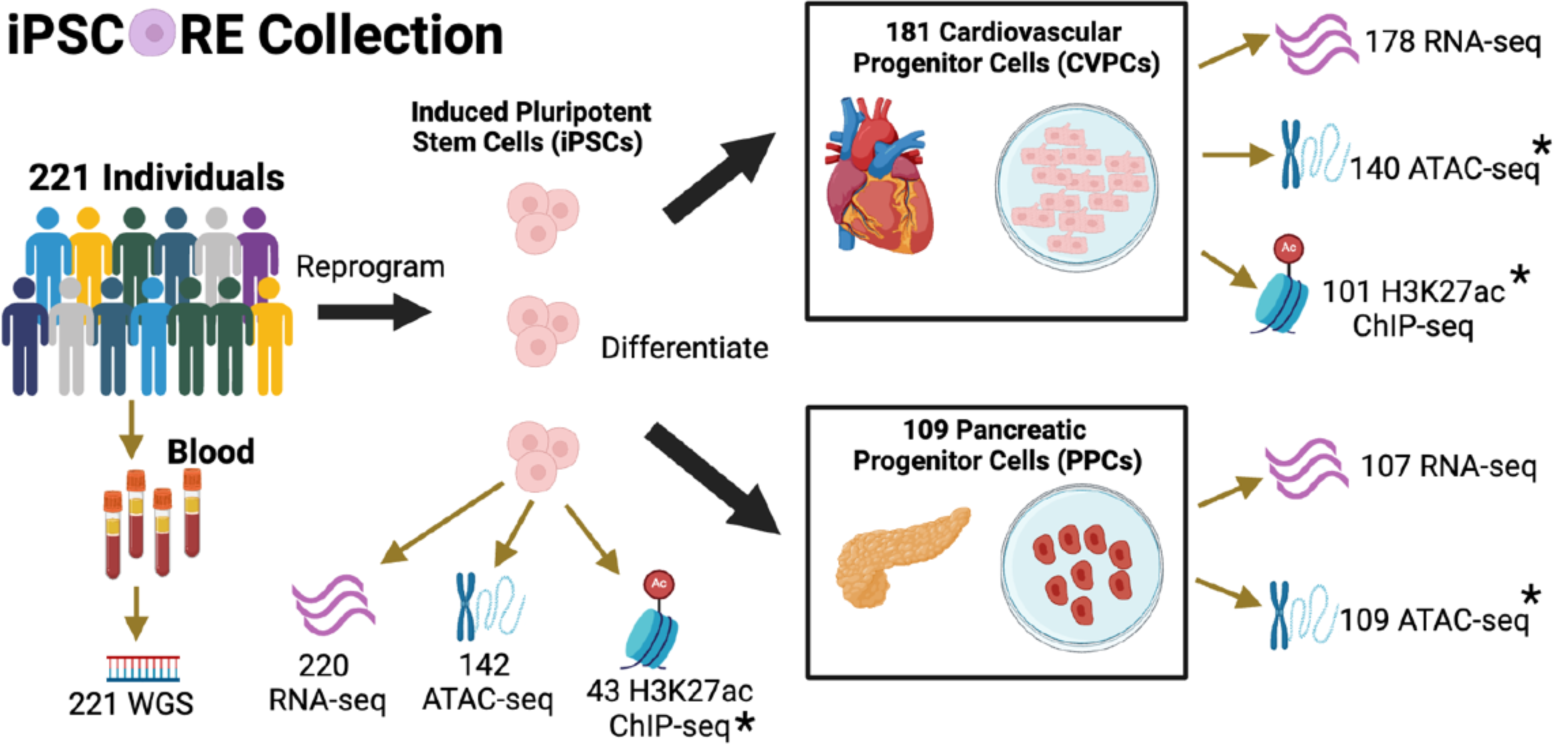
Characterization of iPSCORE Epigenomic Datasets. Overview of the iPSCORE molecular samples generated from blood, reprogrammed iPSCs, and derived tissues. Of the 1,261 molecular samples, 861 were previously published and 400 were newly released in this study. In addition to the 393 samples in the four new molecular data sets (indicated by asterisks), 7 of the 220 iPSC RNA-seq samples were not previously published. WGS analyses identified 16,360,123 single nucleotide polymorphisms (SNPs). The RNA-seq, ATAC-seq, and H3K27ac ChIP-seq libraries were sequenced to median depths of 71.7, 90.9, and 52.1 million reads, respectively (See Methods). The figure was created using Biorender.com.

### iPSCORE early developmental-like tissues display lineage-specific regulatory landscapes

To examine the epigenomic properties of the three EDev-like tissues, we performed ATAC-seq on 142 hiPSCs, 140 CVPCs, and 109 PPCs, as well as H3K27ac ChIP-seq on 43 iPSCs and 101 CVPCs (Figure 1). For each tissue and datatype, we called consensus peaks using a subset of the samples from unrelated individuals (see Methods). In total, we called 208,581 ATAC-seq peaks in iPSCs, 278,471 in CVPCs, and 289,980 in PPCs. Similarly, we called 67,873 consensus H3K27ac ChIP-seq peaks in iPSCs and 63,811 in CVPCs. A UMAP analysis of the ATAC-seq and ChIP-seq peaks showed that samples cluster by tissue, indicating that the iPSCs and the derived fetal-like CVPCs and PPCs each have distinct regulatory landscapes (Figure S2a-b).

To examine the TF binding patterns of the EDev-like tissue-specific and shared regulatory elements (Figure S2c), we performed footprinting analysis to predict TF binding sites (TFBSs) for 1,147 motifs in the consensus ATAC-seq peaks for each tissue^22–24^. TFs with known roles in pluripotency, cardiac, and pancreatic development were strongly enriched in iPSC-, CVPC-, and PPC-specific ATAC-seq peaks, respectively (Figure 2a). For example, NANOG and POU5F1 TFBSs were exclusively enriched in iPSC-specific ATAC-seq peaks, MEF2 and NKX2-5 TFBSs were strongly enriched in CVPC-specific ATAC-seq peaks, and HNF1B, ONECUT1, MEIS1, and PDX1 TFBSs were strongly enriched in PPC-specific ATAC-seq peaks. On the other hand, the shared ATAC-seq peaks were strongly enriched with TFs associated with essential cellular processes including chromatin organization (CTCF, CTCFL) and cell growth during the G1 phase (E2F Family; Figure 2a). As expected, TFs associated with tissue-specific ATAC-seq peaks had higher expression in the corresponding tissue, and TFs associated with binding sites enriched in shared ATAC-seq peaks had similar expression levels in all three tissues (Figure S3).

**Figure 2.**
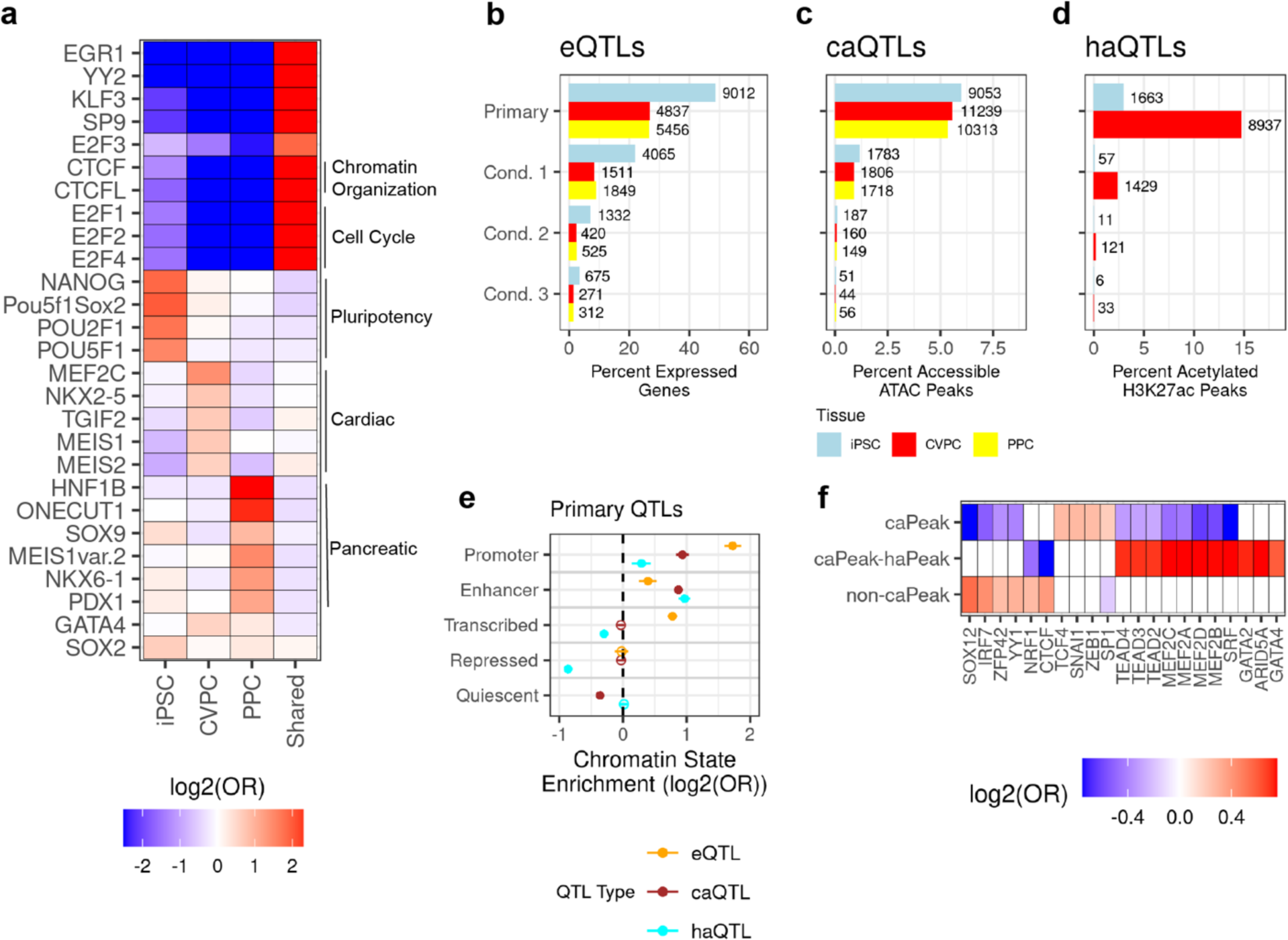
Characterization of multi-omic regulatory variation in early developmental tissues. **a)** Heatmap showing TFBS enrichments in iPSC-, CVPC-, PPC-specific, and shared ATAC-seq peaks. Two-sided Fisher’s Exact tests were performed to test the enrichment (odds ratio) of the predicted TFBSs in each of the four ATAC-seq peak sets. Each cell is filled with the log2(Odds Ratio) of the association between predicted TFBSs (y-axis) and tissue-specific and shared ATAC-seq peaks (x-axis). **b-d)** Bar plots showing the fraction of elements (genes and peaks) with at least one eQTL (**b**), caQTL (**c**), and haQTL (**d**). For eQTLs, variants (MAF > 0.05) within 1 Mb of each gene were tested for an association with gene expression. For chromatin QTLs, variants (MAF > 0.05) within 100 kb of each peak were tested for an association with chromatin accessibility (**c**) or histone acetylation (**d**). If a QTL was discovered for a gene or peak, up to three additional conditional QTLs were tested by using the lead variant as a covariate. The x-axis is the fraction of elements with a QTL for each tissue and the y-axis is QTL type (i.e. Primary or Conditional). Each bar is colored by tissue (iPSC = light blue, CVPC = red, PPC = yellow). **e)** Plot showing the enrichment of primary iPSC and CVPC eQTLs, caQTLs, and haQTLs in chromatin states. The x-axis is the enrichment log2(Odds Ratio) and the y-axis contains the five collapsed chromatin states. The points are colored by the QTL type (eQTL = “orange”, caQTL = “brown”, and haQTL = “light blue”). The whiskers represent the log2 upper and lower 95% confidence intervals. Significant enrichments are represented by filled circles and non-significant enrichments are represented by circles without a fill. Enrichment of conditional iPSC and CVPC eQTLs, caQTLs, and haQTLs in chromatin states is shown in Figure S6. **f)** Heatmap showing the enrichment of TFBSs in CVPC ATAC-seq peaks without caQTLs (non-caPeaks), with caQTLs overlapping haQTLs (caPeak-haPeak), and with caQTLs not overlapping haQTLs (caPeaks). A two-sided Fisher’s Exact test was performed to test the enrichment of TFBSs within these three categories. The x-axis represents the TFBSs, the y-axis corresponds to the ATAC-seq peak annotation, and each cell is filled with the corresponding log2(Odd Ratio) from the Fisher’s Exact test. Non-significant tests (Benjamini-Hochberg adjusted P-value > 0.05) are filled with white.

In summary, tissue-specific regulatory elements in the embryonic-like iPSCs and fetal-like cardiac and pancreatic tissues are bound by appropriate lineage-specific developmental TFs, and shared regulatory elements are bound by TFs governing essential cellular processes such as chromatin organization and cell growth.

### Identification of multi-omic regulatory variation in iPSCORE tissues

To identify and characterize regulatory variation associated with the three molecular phenotypes in the iPSCORE EDev-like tissues, we established a standardized QTL discovery model. Since the iPSCORE cohort contains close relatives, we utilized a linear mixed model to calculate the association between SNP genotypes (5,536,303 with MAF > 5%) and molecular phenotypes (gene expression, open chromatin, H3K27 acetylation), while controlling for relatedness (See Methods).

We identified 30,265 eQTLs (19,305 Primary and 10,960 Conditional) for 12,174 genes across the three tissues (Figure 2b; Table S6). iPSCs had approximately 1.4-fold more eGenes (n=9,012) than the CVPCs (n=4,837) and PPCs (n=5,456) most likely because there were more iPSC samples, and they have lower cellular heterogeneity. To examine the relative power of identifying eQTLs, we compared the three iPSCORE EDev-like tissues to the 49 tissues in the GTEx Consortium^1^ and showed that they had similar eGene discovery rates (Figure S4).

We identified 36,559 caQTLs (30,605 Primary and 5,954 Conditional) for 30,605 caPeaks (ATAC-seq peaks with at least one caQTL, Figure 2c; Table S7). Across all three tissues, between 5-6% of accessible ATAC-seq peaks had a caQTL. We observed that caPeaks are enriched for being near eGenes compared to genes without eQTLs (odds ratio [OR] = 1.49, P-value = 3.7×10^−30^ by two-tailed Fisher’s Exact test [FET]), which is consistent with caPeaks being regulatory elements of neighboring genes whose function is affected by variants.

In the iPSC and CVPC tissues, we identified 12,257 haQTLs (10,600 primary and 1,657 conditional) for 10,600 haPeaks (H3K27ac peaks with at least one haQTL) (Figure 2d; Table S8). Of the 12,257 haQTLs, 10,520 were detected in the CVPCs and 1,737 were detected in iPSCs, reflecting the greater number of CVPC H3K27ac samples (n = 101 CVPCs vs n = 43 iPSCs). Of note, ∼15% of the CVPC H3K27ac peaks had at least one haQTL, which was approximately 3-fold greater than the percent of CVPC ATAC-seq peaks with caQTLs.

We next performed Bayesian fine mapping on all 79,081 multi-omic QTLs and calculated the distance between variants with a causal posterior probability (PP) > 1% and the associated qElement (genes and peaks associated with a QTL). We observed a high proportion of primary eQTLs located in the TSS window (TSS ± 1 kb; 23.3%) and a high proportion of primary caQTLs (47.7%) and haQTLs (47.4%) in the associated caPeak and haPeak, respectively (Figure S5). In all molecular data types, conditional QTLs captured more distal regulatory variation compared to primary QTLs, which may reflect greater uncertainty in the posterior probability assignments of the fine-mapped variants for the conditional caQTLs.

In summary, we identified 79,081 QTLs from the iPSCORE EDev-like tissues, making this one of the largest reports of multi-omic QTLs. We show that caPeaks are enriched for being near eGenes, haPeaks are discovered at a ∼3-fold rate than caPeaks, and that primary and conditional QTLs have different distribution patterns from their qElement.

### Integrative QTL analyses capture variation impacting different types of regulatory elements

To examine whether regulatory variation affecting the three molecular phenotypes is located in different types of regulatory elements, we annotated the iPSC and CVPC QTL lead variants with five chromatin states (Promoters, Enhancers, Transcribed, Repressed, and Quiescent regions). For each QTL condition (primary or conditional), we tested the QTL chromatin state enrichment, using lead variants for non-significant elements (i.e. genes and peaks) as background (Figure 2e; Figure S6). Consistent with previous findings^1,3^, primary eQTLs are most significantly enriched in promoters (OR = 3.3) and transcribed regions (OR = 1.7), exhibit weaker enrichments in enhancers (OR = 1.3), and are strongly depleted in quiescent chromatin (OR = 0.47; Figure 2e). Primary caQTLs are strongly enriched in both promoters (OR = 1.9) and enhancers (OR = 1.8), and primary haQTLs are strongly enriched in enhancers (OR = 2.0) and exhibit weaker enrichments in promoters (OR = 1.2; Figure 2e). On the other hand, conditional QTLs were not as strongly enriched in promoters and enhancers (Figure S6), further substantiating uncertainty in their underlying lead variants (Figure S5). These findings support that eQTLs are biased towards identifying regulatory variation in promoters^1,3^, and suggest that caQTLs capture both promoter- and enhancer-acting regulatory variation while haQTLs primarily capture enhancer-acting regulatory loci.

Recent studies have shown that some TFs are more likely than other TFs to have their binding sites impacted by variants^25,26^. To investigate if different TFs are bound to CVPC caPeaks and non-caPeaks (i.e., peaks not associated with a caQTL), we examined the 55,269 CVPC ATAC-seq peaks with at least one predicted TFBS. Predicted TFBSs for six TFs were enriched in the 51,035 non-caPeaks (Figure 2f) including CTCF TFBSs (OR = 1.3, P-value = 5.6×10^−14^). The binding sites of several cardiac TF markers (i.e., MEF2 TFs) were enriched in the 1,641 caPeaks that overlapped haPeaks, and the 2,593 caPeaks not overlapping haPeaks (Figure 2f). Our observations show that caPeaks harbor different predicted TFBSs than non-caPeaks, which is consistent with previous findings that regulatory variation impacts the binding of some TFs more than others^25,26^.

In summary, we show that regulatory variation affecting the three molecular phenotypes (i.e. gene expression, chromatin accessibility, or histone acetylation) is located in different types of regulatory elements and that caPeaks harbor different predicted TFBSs than non-caPeaks.

### QTL modules are regulatory loci with strong effect sizes impacting multiple elements

The same QTL signal can be associated with multiple qElements^2,14,27,28^, therefore we sought to determine the fraction of qElements within each of the iPSCORE EDev-like tissues that shared QTL signals. Noting the lower confidence in causal variants for conditional QTLs, we performed pairwise Bayesian colocalization between primary eQTLs, caQTLs, and haQTLs separately for each of the tissues and identified a total of 9,667 colocalizations (posterior probability that both signals are shared, PP.H4 ≥ 80%, See Methods) between 10,894 unique QTL signals (18.0% of 60,510 primary QTLs) as well as 49,616 singleton QTLs that did not colocalize with any other QTL and hence affected a single qElement (Table S9).

To generate discrete annotations for QTLs associated with multiple qElements, we created three networks by loading the colocalized QTL pairs as edges for each of the tissues independently and identified 4,402 QTL modules. Across the three tissues, 3,078 (69.9%) of the QTL modules were associated with only two qElements (Figure 3a; Table S10) while the remaining 1,324 (31.1%) were associated with three or more qElements. The largest modules were composed of a colocalized QTL associated with 9 or 10 different qElements; and 3.5% modules (n=169) were composed of a colocalized QTL associated with 5 or more qElements. A total of 2,030 (42.5%) of the modules contained at least one eQTL and at least one chromatin QTL (caQTL and haQTL, Figure 3b), supporting that caQTLs and haQTLs capture regulatory variation affecting regulatory elements that modulate gene expression. The CVPCs had the greatest number of QTL modules, of which 49.3% (1,072) were composed of only caQTLs and haQTLs (Figure 3c). The observation that about half of the CVPC QTL modules only contain caQTLs and haQTLs suggests that these analyses capture enhancer-acting regulatory variation that is missed by eQTL analyses conducted using similarly sized sample sets.

**Figure 3.**
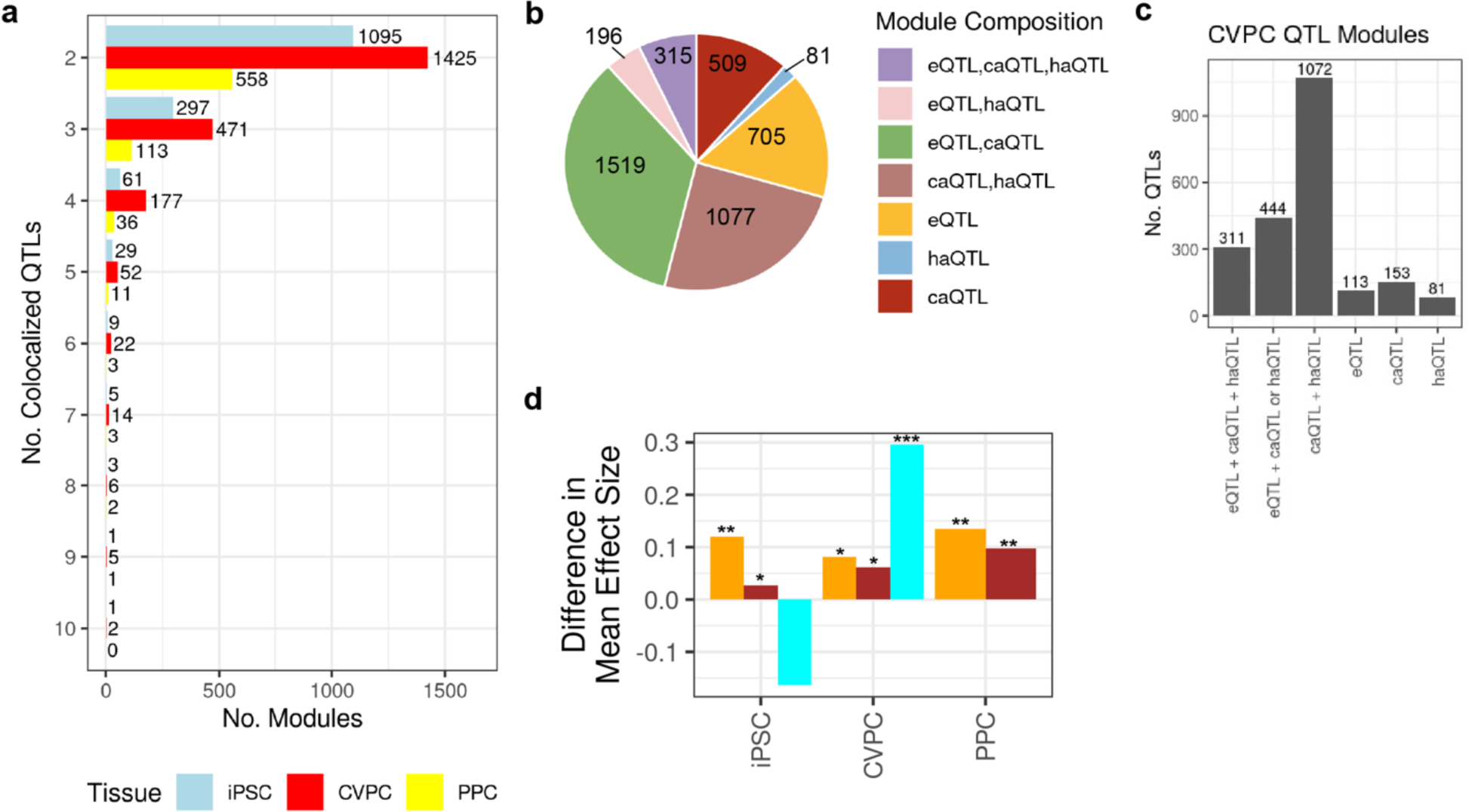
QTLs affect multiple elements across molecular phenotypes. **a)** Bar plot showing the number of QTLs in the 4,402 modules. The x-axis is the number of QTL modules, the y-axis represents the number of colocalized QTLs per module, and the bars are colored by tissue (iPSC = light blue, CVPC = red, and PPC = yellow). **b)** Pie chart showing the number of modules based on their composition. The modules were characterized based on their composition of molecular QTLs (i.e. eQTLs, caQTLs, and haQTLs). **c)** Bar plot displaying the number of CVPC QTLs by module classification. For example, there are 1,072 caQTL and haQTLs in QTL modules exclusively composed of caQTL(s) and haQTL(s). The x-axis is the QTL module classification, and the y-axis is the number of CVPC QTLs. **d)** Bar plot showing the effect size differences between module and singleton QTLs for each dataset. The x-axis corresponds to the eight QTL molecular datasets, the y-axis contains the difference between the mean absolute effect size of QTLs in a module and singleton QTLs, and the bars are colored by molecular QTL type. The asterisks represent the one-sided Wilcoxon P-value significance (P-value < 0.05 = “*”; P-value < 1×10^−20^ = “**”; P-value < 1×10^−30^ = “***”).

Colocalization requires both genetic signals to have sufficiently strong effect sizes, therefore we tested whether the module QTLs had larger absolute effect sizes than singleton QTLs. For seven of the molecular datasets, module QTLs exhibited significantly higher effect sizes than singleton QTLs (Figure 3d). iPSC haQTLs were the only dataset that was not significant likely because of power. The largest difference was observed in CVPC haQTLs (one-sided Wilcoxon test P-value = 7.9×10^−176^), where the module haQTLs mean absolute effect size was 29.6% greater than singleton haQTLs (Figure 3d).

In summary, we identified 4,402 QTL modules composed of 18.0% of all primary QTLs and affecting multiple qElements. We show that module QTLs have stronger effect sizes than singleton QTLs. Notably, we found that nearly half of the QTL modules consist exclusively of caQTLs and haQTLs, underscoring the importance of these QTLs in capturing regulatory variations missed by eQTL analyses.

### Colocalization of iPSCORE multi-omic QTLs with GWAS trait and disease loci

To examine the impact of including caQTLs and haQTLs on the annotation rate of adult GWAS loci, we performed Bayesian colocalization between 6,371 independent GWAS loci from 17 traits (Figure 4a; See Methods), and the 60,510 primary QTLs discovered in the three iPSCORE tissues.

**Figure 4.**
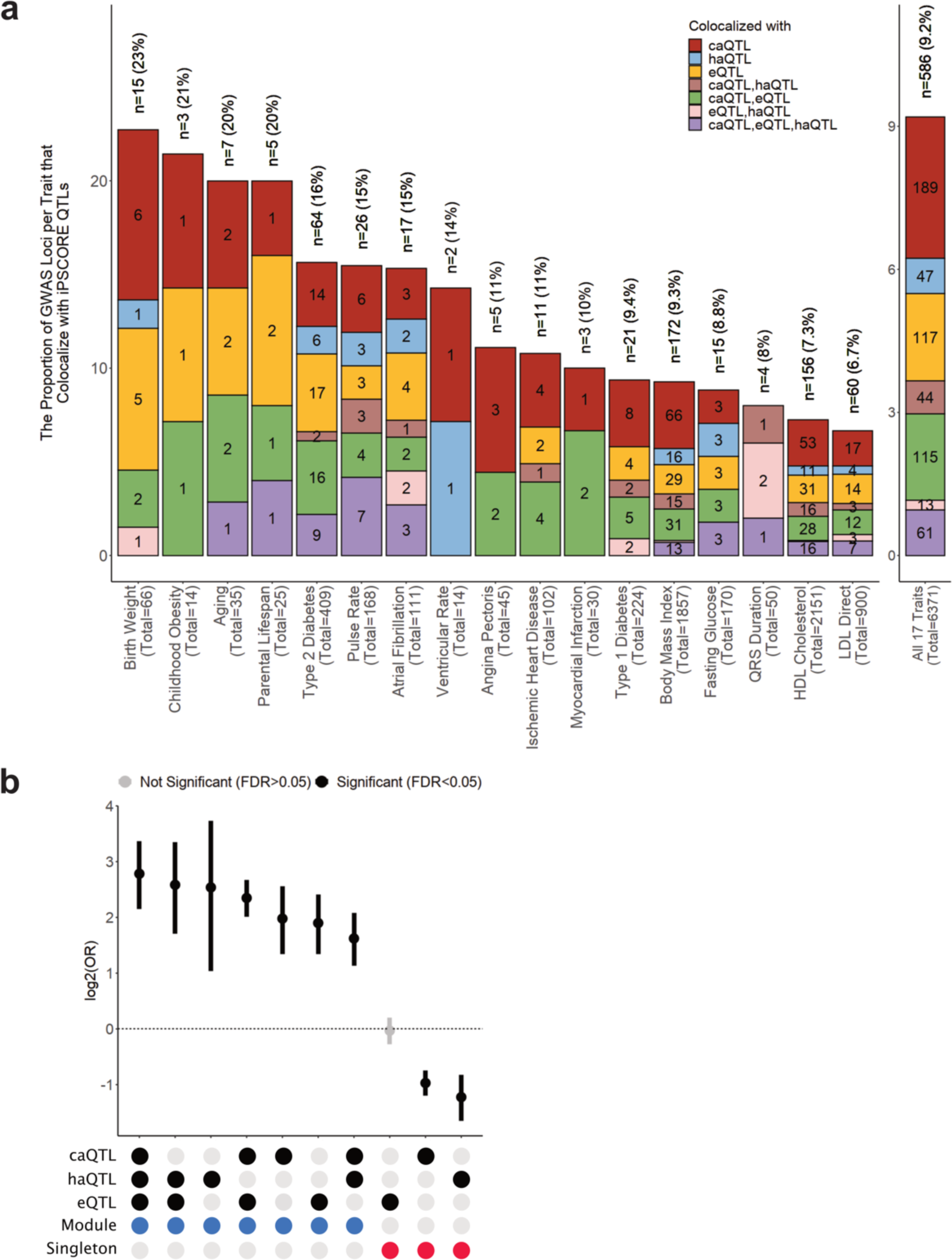
Multi-omic QTLs are enriched for GWAS colocalizations. **a)** Bar plot showing the fraction of GWAS loci that are explained by iPSCORE QTLs. The x-axis contains the GWAS trait name, along with the total number of GWAS loci for each trait, and the y-axis shows the proportion of GWAS loci that colocalize with iPSCORE QTLs. The bars were colored according to the colocalized QTL types (i.e. caQTL-haQTL-eQTL, eQTL-haQTL, eQTL-caQTL, caQTL-haQTL, eQTL, caQTL, haQTL), and the numbers correspond to the number of GWAS loci that colocalized with the QTL types. At the top of each bar, we indicate the total number and fraction of GWAS loci that colocalized with the QTLs. The rightmost bar corresponds to all 586 loci across the 17 GWAS traits that colocalized with an iPSCORE QTL. **b)** Plot showing the enrichment of different QTL combinations for GWAS colocalizations. We categorized each QTL module (indicated by the blue label in the x-axis legend) and singleton (indicated by the red label in the x-axis legend) according to the QTL types they were associated with. Two-sided Fisher’s Exact Tests were performed to test the enrichment (odds ratio) of each category for GWAS colocalization compared to all other categories. P-values were corrected using Benjamini-Hochberg’s Method. Tests that had corrected P-value < 0.05 were considered significant (colored in black).

In total, 9.2% of the GWAS loci (n = 586) across the 17 traits colocalized (PP.H4 ≥ 80%, GWAS P-value < 5×10^−8^, QTL P-value < 5×10^−5^, causal SNP PP ≥ 1%) with one or more of the EDev QTLs (Figure 4a; Table S11). Of the 586 colocalized GWAS loci, 353 (60.2%) colocalized with QTL(s) from only one molecular phenotype, and 233 (39.8%) colocalized with QTLs from two or more molecular phenotypes (Figure 4a; Table S11). Birth weight had the largest proportion of GWAS loci that colocalized with iPSCORE EDev QTLs (n = 15; 23%), including 7 loci that colocalized only with a chromatin QTL (caQTL or haQTL; Figure 4a). In total, 280 (47.8%) GWAS loci only colocalized with caQTLs and/or haQTLs, and thus, including these chromatin QTLs increased the number of GWAS loci annotated with a molecular phenotype by 1.9-fold compared to using eQTLs alone (n = 306, Figure 4a; Table S11).

Given that QTL modules represent complex regulatory loci with strong effect sizes, we sought to determine if they colocalized with GWAS loci at a significantly higher rate than singleton QTLs. We binned the 60,510 primary QTLs into ten categories based on whether they were in QTL modules or were singletons, and their associated molecular phenotypes (Figure 4b). We found that all 7 QTL module categories were enriched for GWAS colocalization while the singleton caQTL and haQTL categories were depleted (Figure 4b). The QTL modules composed of all three molecular types (caQTL-haQTL-eQTL) were the most enriched (Figure 4b). Of the three QTL module categories with only a single molecular type, those containing haQTLs (i.e. two or more haQTLs for different haPeaks colocalized), which likely represent complex active enhancers with large effect sizes, were the most enriched (Figures 2e, 3d, 4b).

Altogether, our results show that including caQTLs and haQTLs results in a marked increase in annotated GWAS loci and support previous observations that GWAS loci are more likely to colocalize with QTLs with strong effect sizes^29^. Our study further extends knowledge about high-effect regulatory variants that underlie adult disease GWAS loci showing that they are enriched for affecting multiple molecular phenotypes.

### CVPC QTL module containing cardiac TF TBX20 colocalizes with Birth Weight locus

To illustrate the functional information gained by colocalizing QTL modules with GWAS loci, we focused on the birth weight GWAS locus on chromosome 7 that colocalized with the CVPC QTL module 47, which is located 12 kb upstream of *TBX20*. CVPC QTL module 47 consists of eQTLs for *TBX20* and four lnc-RNAs (*ENSG00000289335, ENSG00000226063, ENSG00000288947, ENSG00000289089*) and one caPeak (cvpc_atac_peak_233871; Figure 5a-b). *TBX20* encodes a TF that is a key regulator of cardiac development and cardiomyocyte homeostasis^30^. Notably, CVPCs may be an important cell type for birth weight, because infants with a low birth weight have a higher risk of developing congenital heart defects and cardiovascular disease^31,32^. Our fine-mapping identified rs6959887 as the lead variant (chr7:35255754:A>G, causal PP = 17.2%), which has also previously been associated with congenital heart defects^32^. We show that the non-reference allele of rs6959887 is associated with decreased accessibility of a regulatory element (cvpc_atac_peak_233871), decreased expression of *TBX20, ENSG00000289335, ENSG00000288947,* and *ENSG00000289089*, and increased expression of *ENSG00000226063*. Changes in the expression of one or more of these genes may contribute to birth weight and the risk of developing a congenital heart defect or cardiovascular disease later in life. Together, these results illustrate that annotating GWAS loci with QTL modules can help elucidate the underlying molecular mechanisms relevant to development and disease.

**Figure 5.**
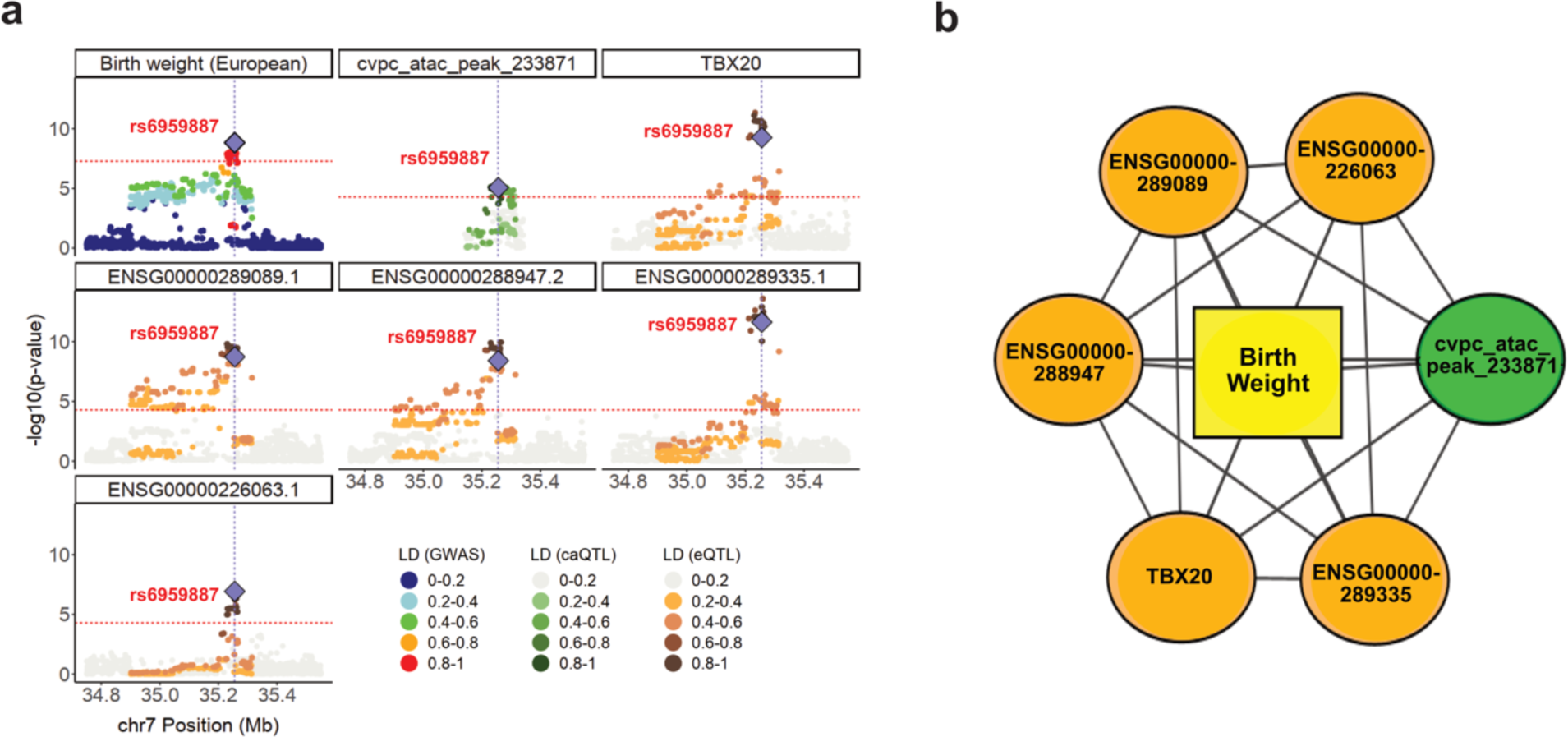
CVPC QTL module containing an eQTL for TBX20 associated with a birth weight GWAS locus. **a)** A birth weight locus colocalized with CVPC QTL Module 47 containing one caPeak and five eGenes, one of which is a known cardiac transcription factor *TBX20*. The fixed genomic coordinates are on the x-axes, and the −log10(P-values) for the associations between the genotype of the tested variants and gene expression or chromatin accessibility are plotted on the y-axes. Red horizontal lines indicate genome-wide significance thresholds for GWAS (P-value = 5×10^−8^) and QTL (P-value = 5×10^−5^) for plotting purposes. Each variant was colored according to their LD with the lead fine-mapped variant (purple diamond; rs6959887, chr7:35255754:A>G, causal PP = 17.2%) using the 1000 Genomes Phase 3 Panel (Europeans only) as reference. **b)** Network diagram showing the colocalized associations between the elements in the CVPC QTL Module 47, in which the edges represent strong evidence of colocalization (PP.H4 ≥ 80%) between the two QTLs (nodes). We observe that all elements had strong colocalization with each other and with the birth weight locus (yellow square), indicating that changes in chromatin accessibility and gene expression at this region are associated with birth weight. Orange nodes represent the CVPC eGenes. The green node represents the CVPC caPeak.

### Identification and functional characterization of early developmental-unique QTLs

Regulatory variation active uniquely during fetal development has been shown to contribute to adult diseases^2,4^, therefore we sought to identify and characterize EDev-unique QTLs (i.e., regulatory variants that are not active in adult tissues); and then examine their overall impact on adult disease and traits. We calculated linkage disequilibrium (LD) between lead fine-mapped variants for all 60,510 primary iPSCORE QTLs and eQTLs and chromatin QTLs from adult datasets^1,33,34^. We identified 7,614 (12.6%) EDev-unique QTLs that were not in LD with any adult QTL including 4,897 caQTLs (16.0% of total 30,605 primary caQTLs), 1,437 haQTLs (13.6% of total 10,600 primary haQTLs), and 1,280 eQTLs (6.6% of total 19,305 eQTLs) (Figure 6a, Tables S7-S9).

**Figure 6.**
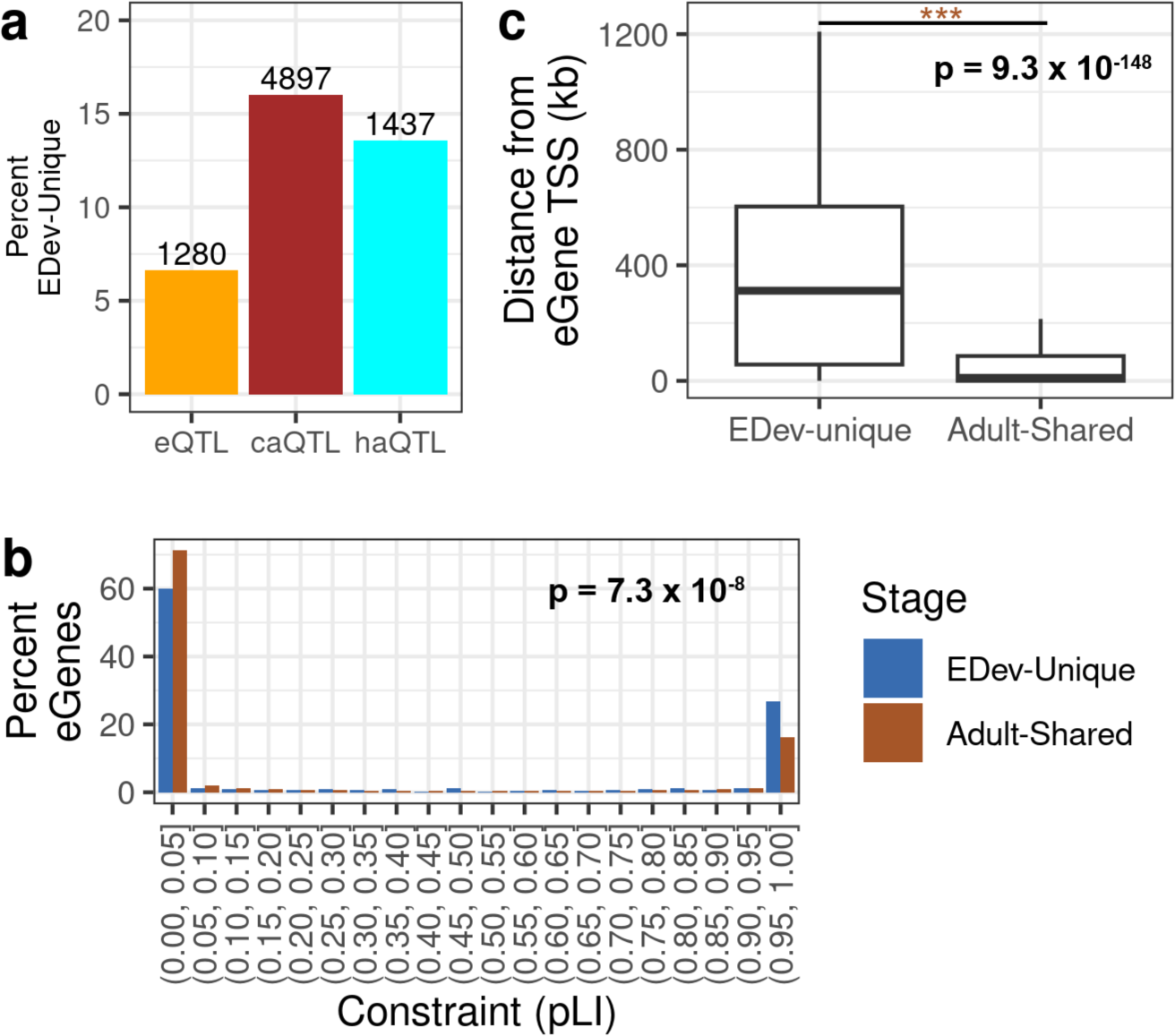
Characterization of EDev-unique QTLs. **a)** Bar plot showing the number of primary EDev-unique QTLs across the three molecular datasets. The x-axis is the molecular QTL type, the y-axis is the percent of QTLs that are EDev-unique, and the top of each bar is labeled with the number of EDev-unique QTLs. **b)** Bar plot showing the distribution of pLI scores for EDev-unique (dark blue) and shared (brown) eGenes. The x-axis reports the pLI range and the y-axis reports the percent of eGenes. The reported P-value is from a one-sided Wilcoxon test. **c)** Box plot showing the distance of primary EDev-unique and primary adult-shared eQTLs from the eGene TSS. The y-axis represents the distance between the eGene TSS window (±1 kb) and the nearest fine-mapped variant with causal PP ≥ 1% (in kilobases). For plot legibility, the y-axis upper limit was set to 1.2 Mb and outliers were omitted. The reported P-value is from a one-sided Wilcoxon test.

Given that disruption of early developmental processes has an impact on lifespan health and disease^35–37^, we investigated if eGenes associated with the EDev-unique eQTLs were under higher evolutionary constraint compared with eGenes associated with adult-shared eQTLs. We annotated 9,132 eGenes in all three tissues with the probability of loss of function tolerance (pLI) from the gnomAD database^38^, then removed 686 eGenes that had both an EDev-unique and adult-shared eQTL, resulting in 532 eGenes with only EDev-unique eQTLs and 8,257 eGenes with only adult-shared eQTLs. We tested the enrichment of EDev-unique eGenes for having strong negative selection (i.e. high evolutionary constraint; pLI ≥ 0.9). We compared the number of EDev-unique versus adult-shared eGenes with evidence for strong purifying selection (pLI ≥ 0.9) and found that EDev-unique eGenes were enriched for high pLI scores (OR = 1.8, P-value = 1.5 x 10^−8^). We then examined the distribution of pLI scores in the EDev-unique and adult-shared eGenes, and confirmed that EDev-unique eGenes have higher pLI scores (one-sided Wilcoxon P-value = 7.3 x 10^−8^; Figure 6b).

Because EDev-unique eGenes were under stronger constraint and therefore less tolerant of deleterious variants, we hypothesized that their eQTLs had weaker effects compared to the adult-shared QTLs. We compared the absolute effect sizes of the EDev-unique and adult-shared eQTLs and observed that EDev-unique eQTLs have significantly lower effect sizes (one-sided Wilcoxon P-value = 3.6 x 10^−10^). We next examined the minimum distance between any fine-mapped variant with causal PP ≥ 1% and the corresponding eGene and observed that compared with adult-shared eQTLs, EDev-unique eQTL variants were significantly further from their eGene TSS (one-sided Wilcoxon P-value = 9.3×10^−148^; Figure 6c).

Noting that EDev-unique QTLs have lower effect sizes compared to adult-shared QTLs, we next examined whether QTL modules were depleted for EDev-unique QTLs. Of the 7,614 primary EDev-unique QTLs, only 545 were in QTL modules, which is a strong depletion compared to adult-shared QTLs (OR = 0.31, P-value = 1.0×10^−183^). Significant depletions of EDev-unique QTLs in modules were observed across the three molecular phenotypes (eQTL OR = 0.26, P-value = 4.4×10^−48^; caQTL OR = 0.31, P-value = 3.1×10^−104^; haQTL OR = 0.49, P-value = 1.4×10^−19^). Because QTL modules are depleted for EDev-unique QTLs, these temporal-specific QTLs are largely singletons impacting the expression single qElements.

Altogether, these results show that in the iPSCORE EDev-like tissues regulatory variation uniquely active during embryonic and fetal development has different characteristics compared with regulatory variation shared with adult tissues. EDev-unique eGenes are under higher evolutionary constraint, and EDev-unique eQTLs have weaker effects, and are more distal from their eGenes compared to adult-shared eQTLs. Our findings also revealed that EDev-unique QTLs are singleton QTLs that affect a single qElement.

### Early developmental-unique QTLs are depleted for colocalizing with GWAS loci

To examine the impact of EDev-unique QTLs on adult disease and traits, we initially determined the fraction that colocalized with GWAS loci compared with adult-shared QTLs. Of the 7,614 EDev-unique QTLs, 82 were in a QTL module that also contained an adult-shared QTL, leaving 7,532 EDev-unique QTLs for this analysis. Similarly, we excluded 109 adult-shared QTLs that were in the same modules as EDev-unique QTLs, leaving 52,787 adult-shared QTLs. We identified 12 GWAS loci that uniquely colocalized with 13 EDev-unique QTLs (0.17% of 7,532), while 1,150 of 52,787 (2.2%) adult-shared QTLs colocalized with the 572 GWAS loci (Figure 7a, Table S11); thus, the EDev-unique QTLs were strongly depleted for GWAS colocalization (two-tailed FET OR = 0.08, P-value = 2.1×10^−49^).

**Figure 7.**
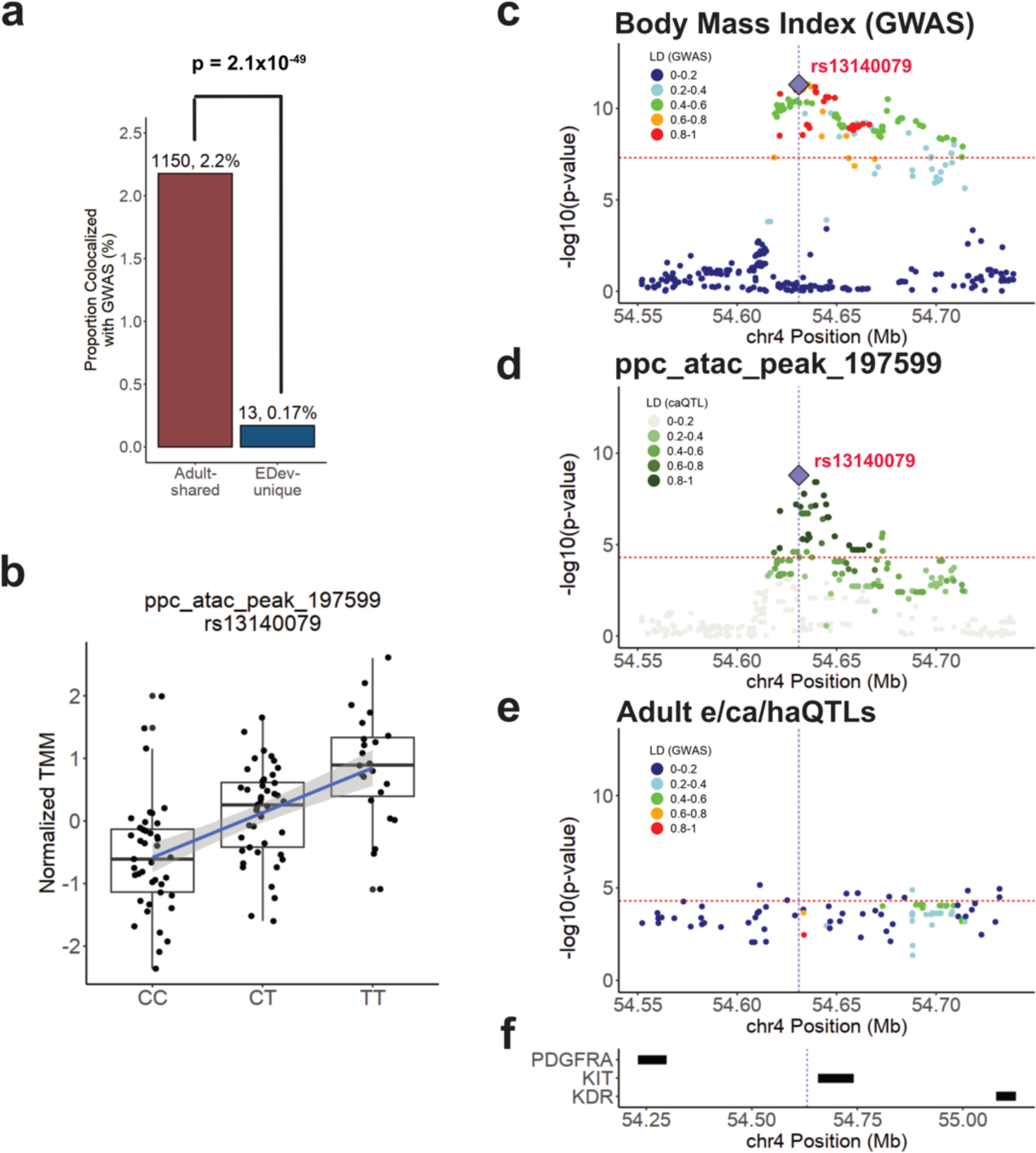
Body Mass Index KIT locus colocalizes with EDev-unique PPC caQTL. **a)** Proportion of adult-shared QTLs (gray) that colocalized with GWAS compared to EDev-unique QTLs (red). P-value was calculated used a two-sided Fisher’s Exact test. **b)** Box plot showing the association between the lead fine-mapped variant (rs13140079, chr4:546310000:C>T, causal PP = 22.4%) and normalized chromatin accessibility of ppc_atac_peak_197599. **c-f)** A body mass index GWAS locus that colocalized with a PPC EDev-unique singleton caQTL. Panel **c** shows the −log10(P-values) for the associations between the genotype of the tested variants and GWAS body mass index. Panel **d** shows the −log10(P-values) for the associations between the genotype of the tested variants and chromatin accessibility. Panel **e** shows the −log10(P-values) for the associations between the genotype of the tested variants and adult gene expression, chromatin accessibility, and histone acetylation. Panel **f** shows the hg38 coordinates for genes within the GWAS locus. Red horizontal lines indicate genome-wide significance thresholds for GWAS (P-value = 5×10^−8^) and QTL (P-value = 5×10^−5^) for plotting purposes. Each variant was colored according to their LD with the lead fine-mapped variant (purple diamond; rs13140079, chr4:546310000:C>T, causal PP = 22.4%) using the 1000 Genomes Phase 3 Panel (Europeans only) as reference. Adult e/ca/haQTL statistics were not available for rs13140079; therefore, rs13140079 is not depicted in Panel **e**.

Because EDev-unique QTLs were enriched for being singletons, we tested their depletion for GWAS colocalization using only the 49,615 singleton QTLs. Of the 7,068 EDev-unique singleton QTLs, only 11 (0.16%) colocalized with a GWAS locus, while 490 of the 42,547 (1.2%) adult-shared singleton QTLs colocalized (Table S11), which again indicates a strong depletion (two-tailed FET OR = 0.13, P-value = 1.5×10^−20^).

These results suggest that the specificity of their activity in early development could account for the depletion of EDev-unique QTLs in GWAS.

### Body Mass Index KIT locus colocalizes only with EDev-unique PPC caQTL

Of the 12 GWAS loci that colocalized only with EDev-unique QTLs, a body mass index GWAS locus on chromosome 4 that colocalized with an EDev-unique PPC caQTL singleton (ppc_atac_peak_197599) (Figure 7b-e), was particularly interesting. Fine mapping identified rs13140079 as the lead variant (chr4:546310000:C>T, causal PP = 22.4%) for this locus, which is strongly associated with ppc_atac_peak_197599 accessibility (Figure 7b). rs13140079 is located 27 kb upstream of *KIT*, which encodes a tyrosine kinase receptor involved in c-KIT signaling (Figure 7f). c-KIT signaling is important for many cellular processes, including cell proliferation, survival, migration, and metabolism, and plays a role in pancreas beta cell development and survival^39^. Mice with a mutation in *Kit* have been shown to develop early onset of diabetes due to impaired glucose tolerance, decreased insulin secretion, and a marked reduction in beta cell mass^39–41^. Our findings suggest that an EDev-unique regulatory element (ppc_atac_peak_197599) located 27 kb upstream of *KIT* may disrupt c-KIT signaling during early pancreas development and influence metabolism and body mass index later in life.

In summary, most of the QTLs discovered in the EDev tissues that colocalized with adult GWAS trait loci were also active in adult tissues, and hence, the inclusion of the EDev-unique QTLs resulted in a relatively small increase in annotated GWAS loci. Nonetheless, we identified 12 adult disease-associated GWAS loci that only colocalized with EDev-unique QTLs, underscoring the importance of considering the temporal dynamics of regulatory variation in disease etiology and trait development.

## Discussion

After the discovery that 90% of the GWAS variants were in intergenic, non-coding regions, eQTL analyses were pursued to understand the mechanisms by which these variants affect gene expression and disease pathologies^42^. Surprisingly, current eQTL analyses only explain approximately 43% of GWAS loci, indicating that they alone are not sufficient to understand genetic associations with disease. There are two leading hypotheses for the poor overlap between GWAS and eQTLs; 1) regulatory variation acts in a context-specific manner (e.g. during fetal development) and with effects that are not present in post-mortem adult donors, and 2) eQTL and GWAS loci have fundamentally different characteristics. Specifically, eQTLs have strong effect sizes and are biased towards promoter-proximal regulatory regions of genes under low evolutionary constraint, while GWAS variants have weak effect sizes and are biased towards distal enhancer regions of genes that are under high evolutionary constraint. We set out to determine if the GWAS colocalization gaps could be overcome by focusing on tissues representing earlier developmental time points, and by mapping chromatin QTLs that specifically capture variation affecting active regulatory elements, such as promoters and enhancers.

In our study, we leveraged multi-omic data from the iPSCORE collection to study regulatory variation that is active during the earliest stages of embryonic, cardiac, and pancreatic development. We defined QTL modules that affected multiple molecular elements and showed that these phenotypically complex loci colocalize with GWAS loci at greater rates than singleton QTLs. Our study also enabled us to examine if chromatin QTLs, which capture regulatory variation affecting promoters, complex enhancers, and other distal regulatory elements, explain GWAS loci that do not colocalize with eQTLs. The integration of chromatin QTLs nearly doubled (1.9-fold increase) the number of GWAS loci that were annotated with iPSCORE EDev eQTLs. This suggests that the fraction of adult disease GWAS loci annotated as harboring identifiable regulatory variation would nearly double to ∼79% if studies comparable in sample size to GTEx conducted integrative eQTL, caQTL, and haQTL analyses.

We also evaluated the fetal origins of adult disease hypothesis, which suggests that missing heritability from integrative GWAS-eQTL studies is explained by temporal regulatory variation that is exclusively active during fetal development, and therefore is not captured in adult tissues^2,4,43^. The EDev-like phenotypic properties of iPSCs and derived tissues provide a powerful model to address this understudied interval of human development. Of the primary QTLs discovered in our study, 12.6% (7,614) are not active in adult tissues and thus are unique to the early development stage. We showed that EDev-unique QTLs have weaker effect sizes than adult-shared QTLs and that genes with EDev-unique eQTLs are under higher evolutionary constraint compared to genes with QTLs that are adult-shared. These observations suggest that EDev-unique regulatory variation is strongly selected against. Despite GWAS and EDev-unique eQTLs both affecting genes under high evolutionary constraint, we observed a strong depletion for EDev-unique QTLs colocalizing with GWAS loci. Further studies with large sample sizes are needed to determine whether this strong depletion results from the biological characteristics of regulatory variation that is uniquely active in early development, or from colocalization limitations due to weak effect sizes. Evidence for the former would support that early developmental-specific regulatory variation plays a minimal role in adult disease and traits. Nevertheless, these findings suggest that QTL studies composed of early developmental tissues similar in sample size to GTEx will find limited numbers of temporal-specific QTLs colocalizing with adult disease GWAS loci.

In summary, we identified 79,081 QTLs that affect gene expression, chromatin accessibility or H3K27 acetylation using 1,261 molecular assays (including WGS) from iPSCs, CVPCs, and PPCs in the iPSCORE collection, making this one of the largest QTL studies conducted using paired multi-omic data. By characterizing the properties of the QTLs, we showed that integrating chromatin QTLs can explain a large fraction of GWAS loci that are not explained by eQTLs. We also demonstrated that EDev-unique QTLs are associated with genes under evolutionary constraint and showed strong depletion in colocalization with GWAS loci. Our study provides biological insights into the characteristics of regulatory variation underlying GWAS loci and QTLs and provides a valuable resource for guiding experimental investigation of disease-associated regulatory variation.

## Code Availability

Scripts for processing FASTQ files and performing downstream analyses are publicly available at https://github.com/frazer-lab/iPSCORE_Multi-QTL_Resource.

## Data Availability

FASTQ sequencing data for 140 CVPC ATAC-seq, 109 PPC ATAC-seq, 43 iPSC H3K27ac ChIP-seq, and 101 CVPC H3K27ac ChIP-seq samples have been deposited into GSE261376. FASTQ sequencing data for iPSC ATAC-seq and PPC RNA-seq are available at GSE203377 and GSE182758, respectively. WGS data for iPSCORE subjects were downloaded as a VCF file (hg19) from phs001325. GWAS summary statistics were obtained from the Pan UK BioBank resource (https://pan.ukbb.broadinstitute.org/), the MAGIC (Meta-Analyses of Glucose and Insulin-related traits) Consortium (https://magicinvestigators.org/downloads/; https://doi.org/10.1038/s41588-021-00852-9), the DIAMANTE Consortium (https://diagram-consortium.org/downloads.html; http://doi.org/10.1038/s41588-018-0241-6), and a previously published studies^44–46^. Full QTL summary statistics, phenotype matrices, element coordinates, TFBS predictions, and colocalization results have been deposited in Figshare: XXX.

## Author information

**iPSCORE Consortium, University of California, San Diego, La Jolla, CA, 92093, US**

Angelo D. Arias, Timothy D. Arthur, Paola Benaglio, Victor Borja, Megan Cook, Matteo D’Antonio, Agnieszka D’Antonio-Chronowska, Christopher DeBoever, Margaret K.R. Donovan, KathyJean Farnam, Kelly A. Frazer, Kyohei Fujita, Melvin Garcia, Olivier Harismendy, David Jakubosky, Kristen Jepsen, Isaac Joshua, He Li, Hiroko Matsui, Naoki Nariai, Jennifer P. Nguyen, Daniel T. O’Connor, Jonathan Okubo, Fengwen Rao, Joaquin Reyna, Lana Ribeiro Aguiar, Bianca Salgado, Nayara Silva, Erin N. Smith, Josh Sohmer, Shawn Yost, William W. Young Greenwald

## Contributions

TDA, JPN, and KAF conceived the study. JPN, TDA, BH, and JJ performed the computational analyses. KAF, MD, GM, JCIB, and iPSCORE consortium members oversaw the study. ADC, NS, and members of the iPSCORE Consortium performed the differentiations and generated molecular data. TDA, JPN, and KAF prepared the manuscript.

## Acknowledgments

This work was supported by the National Library Training Grant T15LM011271, the National Institute of Diabetes and Digestive and Kidney Disease (NIDDK) F31DK131867, U01DK105541, DP3DK112155 and P30DK063491, the National Heart, Lung and Blood Institute (NHLBI) F31HL158198 and U01HL107442, and the National Human Genome Research Institute (NHGRI) RM1HG011558 and R41HG008118. Additional support was also received from a California Institute for Regenerative Medicine grant GC1R-06673-B, NSF-CMMI division award 1728497. This publication includes data generated at the UC San Diego IGM Genomics Center utilizing an Illumina NovaSeq 6000 that was purchased with funding from a National Institutes of Health SIG grant S10OD026929.

## METHODS

### 1. Subject Information

The iPSCORE collection consists of whole genome sequences (WGS) for 273 iPSCORE subjects, 238 iPSCs derived from 221 of these individuals (Figure S1, Table S1) as well as iPSC-derived cell types (cardiovascular progenitor cells [CVPC] and pancreatic progenitor cells [PPC]) with RNA-seq, ATAC-seq and H3K27 acetylation ChIP-seq. Of the 221 individuals with iPSCs, 141 belong to 40 families composed of two or more subjects (range: 2–14 subjects) and 80 genetically unrelated individuals (some individuals were in the same family but only related by marriage)^7^. Each subject was assigned a Universal Unique Identifier (UUID) and an iPSCORE_ID (i.e, iPSCORE_4_1) which designates family (4) and individual number (1). Sex, age, and self-reported race/ethnicity were recorded at the time of enrollment. We previously estimated the ancestry of each subject by comparing their genomes to those of individuals in the 1000 Genomes Project (KGP)^7^. Recruitment of individuals was approved by the Institutional Review Boards of the University of California, San Diego, and The Salk Institute (project no. 110776ZF). The individual iPSC lines in the iPSCORE resource are available to non-profit organizations through WiCell Research Institute (www.wicell.org). Non-profit organizations interested in obtaining the entire iPSCORE collection and for-profit organizations can contact the corresponding author directly to discuss the availability of iPSC lines as well as differentiated cell types.

### 2. Molecular Data Sources

We used the following datasets from the iPSCORE resource.

- 50X WGS (Illumina; 150 bp paired-end) generated from the blood or skin fibroblasts of the 273 iPSCORE subjects^6^.
- RNA-seq data (Table S2) from:

- 220 iPSC lines from 220^干^ individuals^6^
- 178 CVPCs derived from 147 iPSC lines from 137 individuals^4,5,12^
- 107 PPCs derived from 106 iPSC lines from 106 individuals^2^
- ATAC-seq data (Table S3) from:

- 142 iPSC lines from 129 individuals^17^
- 140 CVPCs derived from 132 iPSC lines from 124 individuals *
- 109 PPCs derived from 108 iPSC lines from 108 individuals *
- H3K27ac ChIP-seq data (Table S4) from:

- 43 iPSC lines from 38 individuals *
- 101 CVPCs derived from 97 iPSC lines from 96 individuals *

*The datasets previously published are referenced while the four datasets released here are indicated by an asterisk. In addition to the samples in these four new datasets, 7 of the 220 iPSC RNA-seq samples were not previously published.

^干^There are 221 subjects in the study and 220 iPSC RNA-seq samples from 220 subjects. One subject had CVPC and PPC RNA-seq samples but not an iPSC RNA-seq sample.

### 3. iPSC generation and differentiation

#### 3.1 iPSC generation

Generation of the 238 iPSC lines has previously been described in detail^7^. Briefly, cultures of primary dermal fibroblast cells were generated from a punch biopsy tissue, expanded for approximately 3 passages, and cryopreserved. In batch, the fibroblasts were thawed and plated at a density of 2.5×10^5^ cells/well of 6-well plate and infected with the Cytotune Sendai virus (Life Technologies) per manufacturer’s protocol to initiate reprogramming. The Sendai infected cells were maintained with 10% FBS/DMEM (Invitrogen) for Days 4-7 until the cells recovered and repopulated the well. These cells were then enzymatically dissociated using TrypLE (Life Technologies) and seeded onto a 10-cm dish pre-coated with mitotically inactive-mouse embryonic fibroblasts (MEFs) at a density of 5×10^5^ cells/dish and maintained with hESC medium, as previously described. Emerging iPSC colonies were manually picked after Day 21 and maintained on Matrigel (BD Corning) with mTeSR1 medium (Stem Cell Technologies). From each individual, multiple independently established iPSC clones (i.e. referred to as lines) were derived, cultured typically to passage 12 (P12), and then cryopreserved. Sendai virus clearance typically occurred at or before P9 and was not detected in the iPSC lines at the P12 stage of cryopreservation. A subset of the iPSC lines was evaluated by flow cytometry for expression of two pluripotent markers: Tra-1-81 (Alexa Fluor 488 anti-human, Biolegend) and SSEA-4 (PE anti-human, Biolegend). Pluripotency was also examined using PluriTest-RNAseq^47^.

Harvesting of material for molecular assays:

a. At P12, 220 iPSC lines from 220 individuals, pellets were collected and frozen in RTL plus buffer (Qiagen) for the RNA-seq assay.
b. iPSC nuclear pellets for the ATAC-seq and H3K27ac ChIP-seq assays were collected during the CVPC differentiation protocol (see below: 3.2 CVPC differentiation).

#### 3.2 CVPC differentiation

As previously described in detail^48^, to generate CVPCs we used a small molecule cardiac differentiation protocol^49^. The 25-day differentiation protocol consisted of four phases:

1. *Expansion of iPSC*: One vial of each iPSC line was thawed into mTeSR1 medium containing 10 μM ROCK Inhibitor (Sigma) and plated on one well of a 6-well plate coated overnight with matrigel. During the expansion phase cells were cultured in mTeSR. The iPSCs were passaged using Versene (Lonza) from one well into three wells of a 6-well plate. Next, the iPSCs were passaged using Versene onto three 10 cm dishes at 2.5×10^4^ per cm^2^ density. The iPSCs monolayer was plated onto three T150 flasks at the density of 3.7 x 10^4^ per cm^2^ using Accutase (Innovative Cell Technologies Inc.). iPSCs were at passage 22.7 ± 4.8 (range 17 to 44) at the monolayer stage (i.e., initiation of differentiation). When iPSC lines were visually estimated to be at 80% confluency they were passaged in mTeSR1 medium containing 5 μM ROCK inhibitor.
2. *Differentiation:* After reaching 80% confluency differentiation (D0) was initiated with the addition of the medium containing RPMI 1960 (Gibco-life technologies) with Penicillin – Streptomycin (Gibco/Life Technologies) and B-27 Minus Insulin (Gibco/Life Technologies) (hereafter referred to as RPMI Minus supplemented with 12μM CHIR-99021. After 24h of exposure to CHIR-99021 (D1), medium was changed to RPMI Minus. On D3 medium was changed to 1:1 mix of spent and fresh RPMI Minus, supplemented with 7.5μM IWP-2 (Tocris). On D5, after 48h of exposure to IWP-2, the medium was changed to RPMI Minus. On D7, medium was changed to RPMI 1960 with Penicillin – Streptomycin (Gibco/Life 22 Technologies) and B-27 Supplement 50X (hereafter referred to as RPMI Plus) (Gibco/Life Technologies). Between D7 and D13, RPMI Plus medium was changed every 48h. The first beating cells were usually observed between D7 and D9 and as early as D7 (immediately after the media change) and robust beating was usually observed between D8 and D11.
3. *Purification:* Since fetal cardiomyocytes have a higher capacity to use lactate as a primary energy source than other cell types^50,51^, we incorporated lactate metabolic selection for five days to increase CVPC purity^52^. On D15, the cells were collected from the flask using Accutase and plated onto fresh T150 flasks at confluency 1-1.3 x 10^6^ per cm^2^. On D16, cells were washed with PBS without Ca^2+^ and Mg^2+^ (Gibco/Life Technologies) and medium was changed for RPMI 1960 no glucose (Gibco/Life Technologies) supplemented with Non-Essential Amino Acids (Gibco/Life Technologies), L-Glutamine (Gibco/Life Technologies), Penicillin-Streptomycin 10,000U (Gibco/Life Technologies) and 4mM Sodium L-Lactate (Sigma) in 1M HEPES (Gibco/Life Technologies). Medium supplemented with lactate was changed on D17 and D19. During the lactate selection, CVPCs were beating robustly less than 16 hours after reseeding.
4. *Recovery:* After metabolic selection, CVPCs were maintained in cell culture for five days. On D21, cells were washed with PBS, and the medium was changed to RPMI Plus. We changed the media on D23 using RPMI Plus. To evaluate the efficiency of CVPC differentiation, we performed flow cytometry on D25 with the cardiac marker Troponin T (cTnT, TNNT2). Specifically, 5×10^5^ CVPCs were permeabilized and blocked in 0.5% BSA, 0.2% TX-100 and 5% goat serum in PBS for 30 minutes at room temperature. Cells were stained with Troponin T, Cardiac Isoform Ab-1, Mouse Monoclonal Antibody (Thermo Scientific, MS-295-P0) at 4°C for 45 minutes, followed by Alexa Fluor 488 secondary antibody (Life Technologies, A11001). Stained cells were acquired using BD FACSCanto II system (BD Biosciences) and the fraction of cTnT-positive cells were calculated using FlowJo software version 10.2^12^.

Harvesting of material for molecular assays:

a. At D0, 142 iPSC lines from 129 individuals were collected and frozen as nuclear pellets for the ATAC-seq assay.
b. At D0, 43 iPSC lines from 41 individuals were collected and frozen as nuclear pellets for the H3K27ac ChIP-seq assay.
c. At D25, 178 CVPCs derived from 147 iPSC clones from 137 individuals, pellets were collected and frozen in RTL plus buffer (Qiagen) for the RNA-seq assay.
d. At D25, 140 CVPCs derived from 132 iPSC clones from 124 individuals were collected and frozen as nuclear pellets for the ATAC-seq assay.
e. At D25, 101 CVPCs derived from 97 iPSC clones from 96 individuals were collected and cross-linked for the H3K27ac ChIP-seq assay.

#### 3.3 PPC Differentiation

As previously described, the iPSC lines were differentiated into PPCs using the STEMdiff^TM^ Pancreatic Progenitor Kit (StemCell Technologies, Catalog #05120) protocol with minor modifications^2^. Briefly, iPSC lines were thawed into mTeSR1 medium containing 10 µM Y-27632 ROCK Inhibitor (Selleckchem) and plated onto one well of a 6-well plate coated with Matrigel. iPSCs were grown until they reached 80% confluency and then passaged using 2mg/ml solution of Dispase II (ThermoFisher Scientific) onto three wells of a 6-well plate (ratio 1:3). To expand the iPSC cells for differentiation, iPSCs were passaged a second time onto six wells of a 6-well plate (ratio 1:2). When the iPSCs reached 80% confluency, cells were dissociated into single cells using Accutase (Innovative Cell Technologies Inc.) and resuspended at a concentration of 1.85 x 10^6^ cells/ml in mTeSR medium containing 10 µM Y-27632 ROCK inhibitor. Cells were then plated onto six wells of a 6-well plate and grown for approximately 16 to 20 hours to achieve a uniform monolayer of 90-95% confluence (3.7 x 10^6^ cells/well; about 3.9 x 10^5^ cells/cm^2^). Differentiation of the iPSC monolayers was initiated by replacing the mTeSR medium with STEMdiff^TM^ Stage Endoderm Basal medium supplemented with Supplement MR and Supplement CJ (2 ml/well) (Day 1, D1). The following media changes were performed every 24 hours following initiation of differentiation (2 ml/well). On D2 and D3, the medium was changed to fresh STEMdiff^TM^ Stage Endoderm Basal medium supplemented with Supplement CJ. On D4, the medium was changed to STEMdiff^TM^ Pancreatic Stage 2-4 Basal medium supplemented with Supplement 2A and Supplement 2B. On D5 and D6, the medium was changed to STEMdiff^TM^ Pancreatic Stage 2-4 Basal medium supplemented with Supplement 2B. From D7 to D9, the medium was changed to STEMdiff^TM^ Pancreatic Stage 2-4 Basal medium supplemented with Supplement 3. From D10 to D14, the medium was changed to STEMdiff^TM^ Pancreatic Stage 2-4 Basal medium supplemented with Supplement 4. On D15, to evaluate the efficiency of PPC differentiation, we performed flow cytometry on two pancreatic precursor markers, PDX1 and NKX6-1. Specifically, at least 2 x 10^6^ cells were fixed and permeabilized using the Fixation/Permeabilized Solution Kit with BD GolgiStop TM (BD Biosciences) following the manufacturer’s recommendations. Cells were resuspended in 1x BD Perm/Wash TM Buffer at a concentration of 1 x 10^7^ cells/ml. For each flow cytometry staining, 2.5 x 10^5^ cells were stained for 75 minutes at room temperature with PE Mouse anti-PDX1 Clone-658A5 (BD Biosciences; 1:10) and Alexa Fluor® 647 Mouse anti-NKX6.1 Clone R11-560 (BD Bioscience; 1:10), or with the appropriate class control antibodies: PE Mouse anti-IgG1 κ R-PE Clone MOPC-21 (BD Biosciences) and Alexa Fluor® 647 Mouse anti IgG1 κ Isotype Clone MOPC-21 (BD Biosciences). Stained cells were washed three times, resuspended in PBS containing 1% BSA and 1% formaldehyde, and immediately analyzed using FACS Canto II flow cytometer (BD Biosciences). The fraction of PDX1- and NKX6-1-positive cells were calculated using FlowJo software version 10.4^2^.

Harvesting of material for molecular assays:

a. At D15, 107 PPCs derived from 106 iPSC clones from 106 individuals, pellets were collected and frozen in RTL plus buffer (Qiagen) for the RNA-seq assay.
b. At D15, 109 PPCs derived from 108 iPSC clones from 108 individuals were collected and frozen as nuclear pellets for the ATAC-seq assay.

### 4. Molecular data generation and sequencing

#### 4.1 Whole Genome Sequencing

As previously described^6,7^, we generated whole genome sequences (WGS) for 273 iPSCORE subjects, though only 221 had their fibroblasts reprogrammed into 238 iPSC lines. Genomic DNA was isolated from whole blood (or in 19 cases directly from the fibroblasts) using DNEasy Blood & Tissue Kit (Qiagen), quantified, normalized, and sheared with a Covaris LE220 instrument. The samples were normalized to 1 μg and submitted to Human Longevity (HLI) for whole genome sequencing. DNA libraries were prepared (TruSeq Nano DNA HT kit, Illumina), characterized with regards to size (LabChip DX Touch, Perkin Elmer) and concentration (Quant-iT, Life Technologies), normalized to 2-3.5nM, combined into 6-sample pools, clustered and sequenced to ∼ 30X depth on the Illumina HiSeqX (150 bp paired-end).

#### 4.2 RNA-seq

As previously described in detail^6^, for the iPSCs, total RNA was isolated from cell lysates using AllPrep DNA/RNA Mini Kit (QIAGEN). RNA quality was assessed based on RNA integrity number (RIN) using an Agilent Bioanalyzer, and libraries were prepared using the Illumina TruSeq stranded mRNA kit and sequenced with 100 bp paired-end reads on an Illumina HiSeq2500 (an average of 22 million read pairs/per sample).

As previously described in detail^4,5,12^, for the CVPCs, total RNA was isolated from cell lysates using the Quick-RNA^TM^ MiniPrep Kit (Zymo Research). RNA quality was assessed based on RIN, and libraries were prepared using the Illumina TruSeq stranded mRNA kit and sequenced with either 100 bp paired-end or 150 bp paired-end reads on an Illumina HiSeq4000 (an average of 28 million read pairs/per sample).

As previously described in detail^2^, for the PPCs, total RNA was isolated from cell lysates using the Quick-RNA^TM^ MiniPrep Kit (Zymo Research), RNA quality was assessed based on RIN, and libraries were prepared using the Illumina TruSeq stranded mRNA kit and sequenced with 100 bp paired-end reads on an Illumina NovaSeq 6000 (an average of 71 million read pairs/sample).

#### 4.3 ATAC-seq

All ATAC-seq samples were processed in the same manner using a modified version of the Buenrostro et al. protocol^18^ as previously described^17^. Briefly, frozen nuclear pellets of 2.5×10^4^ iPSC or 1×10^5^ CVPC or PPC cells were thawed on ice and tagmented in total volume of 25μl in permeabilization buffer containing digitonin (10mM Tris-HCl pH 7.5, 10mM NaCl, 3mM MgCl^2^, 0.01% digitonin) and 2.5μl of Tn5 from Nextera DNA Library Preparation Kit (Illumina) for 45-75min at 37°C in a thermomixer (500 RPM shaking). We included a double size selection step during purification using AMPure XP DNA beads (Beckman Coulter). To eliminate confounding effects due to index hopping, all libraries within a pool were indexed with unique pairs of i7 and i5 barcodes. Libraries were amplified for 12 cycles using NEBNext^®^ High-Fidelity 2X PCR Master Mix (NEB) in total volume of 25µl in the presence of 800nM of barcoded primers (400nM each) custom synthesized by Integrated DNA Technologies (IDT) and sequenced with either 100 bp paired-end and 150 bp paired-end reads on an Illumina HiSeq4000 for iPSCs and CVPCs and 150 bp paired-end reads on an Illumina NovaSeq 6000 for PPCs.

#### 4.4 H3K27 acetylation ChIP-seq data

All H3K27ac ChIP-seq samples were processed in the same manner. For H3K27ac, 5-15 x 10^6^ formaldehyde crosslinked cells were lysed and sonicated in 110µl of SDS Lysis Buffer (0.5% SDS, 50mM Tris-HCl pH 8.0, 20mM EDTA, 1x cOmplete™ Protease Inhibitor Cocktail (Sigma)) using Covaris E220 Focused-ultrasonicators (Covaris) for 14 cycles, 1 min per cycle, duty cycle 5. For each sample, H3K27ac antibody (Abcam ab4729, lot GR00324078) was coupled for 4 hours to 40µl of 1:1 mix Protein G and Protein A Dynabeads (Thermo Scientific) and used for overnight chromatin immunoprecipitation in IP buffer (1% Triton X-100, 0.1% DOC, 1x TE buffer, 1x cOmplete™ Protease Inhibitor Cocktail). Beads with immunoprecipitated chromatin were washed with 150µl of following buffer: four times with RIPA Low Salt Buffer (0.1% SDS, 1% Triton X-100, 2mM EDTA, 20mM Tris-HCl pH 8.0, 300mM NaCl, 0.1% DOC), two times in RIPA High Salt Buffer (0.1% SDS, 1% Triton X-100, 1mM EDTA, 20mM Tris-HCl pH 8.0, 500mM NaCl, 0.1% DOC), twice in LiCl Buffer (250mM LiCl, 0.5% NP-40, 0.5% DOC, 1mM EDTA, 10mM Tris-HCl pH 8.0) and twice in 1X TE buffer (10mM Tris-HCl pH 8.0, 1mM EDTA). Next samples were eluted in 150 µl of Direct Elution Buffer (0.1% SDS, 10mM Tris-HCl pH 8.0, 5mM EDTA) and reverse crosslinked by incubation for 15 min at 65°C with rotation and subsequent incubation with 5 µl RNAse (Sigma) for 1h at 37°C and Proteinase K Solution (20 mg/mL, Thermo Fisher Scientific) for 1h at 55°C. After reverse crosslinking samples were purified with 2X Agencourt AMPure XP DNA beads (Beckman Coulter), eluted in 30 µl of H_2_O and Qubit (Thermo Scientific) quantified. Libraries were generated using KAPA Hyper Prep Kit (KAPA Biosystems) and KAPA Real Time Library Amplification Kit (KAPA Biosystems) following manufacturers manual. Libraries were barcoded using TruSeq RNA Indexes (Illumina), size selected for 300 bp to 500 bp, and sequenced with either 100 bp paired-end or 150 bp paired-end reads on an Illumina HiSeq 4000 (an average of 44 million read pairs/per sample).

### 5. Data Processing

#### 5.1 WGS

We downloaded the VCF in hg19 for 273 iPSCORE individuals^6,7^ (see **4.1 Whole Genome Sequencing**) from dbGaP (phs001325.v4.p1), phased the 273 WGS with the Michigan Imputation Server using the 1000 Genomes 30x GRCh38 as the reference panel^53–55^, and performed liftOver to hg38 using CrossMap^56^ and the hg38 reference genome from UCSC (https://hgdownload.soe.ucsc.edu/goldenPath/hg38/bigZips/).

#### 5.2 RNA-seq

All RNA-seq samples were processed in a uniform manner (Table S2). Libraries that were sequenced more than once were merged by concatenating the FASTQ files. The reads were aligned onto the hg38 human reference genome downloaded from Gencode version 44^57,58^ using STAR 2.7.10b (https://github.com/alexdobin/STAR) with the following parameters: --outSAMattributes All --outSAMunmapped Within --outFilterMultimapNmax 20 --outFilterMismatchNmax 999 -- alignIntronMin 20 --alignIntronMax 1000000 --alignMatesGapMax 1000000. PCR duplicates were marked with Picard (https://github.com/broadinstitute/picard) (v3.1.0) and counted using samtools flagstat^59^ (v1.17). Number and percentage of mapped reads were calculated using samtools flagstat^59^ (v1.17). Percentage of intergenic and mRNA bases were determined using Picard (v3.1.0) CollectRnaSeqMetrics. Gene TPM expression and read counts were calculated using RSEM^60^ (v1.3.3) with gene annotations from Gencode version 44^57,58^ (hg38) and the following parameters: --seed 3272015 --estimate-rspd --forward-prob 0 --paired-end. RNA-seq samples were examined for quality using GTEx standards^1^. Specifically, we required that samples met the following metrics: 1) the number of mapped reads > 10 million; 2) percent of intergenic bases < 30; 3) percent of mRNA bases > 70; 4) percent of duplicate reads < 30; 5) percent mapped reads > 85%; 6) number of reads passing filters > 25M, and 7) matched via a sample identity check to the correct subject with PI_HAT > 0.90 (Figure S7).

For eQTL mapping, gene expression values were normalized and filtered using the same procedure as GTEx^1^. Specifically, 1) read counts were TMM normalized across all genes using edgeR^61^ (v3.38.4) with functions *DGEList*, *calcNormFactors*, and *cpm*; 2) autosomal genes were selected and filtered based on expression thresholds of ≥ 0.1 TPM in ≥ 20% of samples and ≥ 6 reads (unnormalized) in ≥ 20% of samples; 3) TMM expression values for each gene were inverse normal transformed across samples using *rank* and *qnorm* in R v4.2.1 and used as input for eQTL analyses. This resulted in 18,720 iPSC, 18,314 CVPC, and 20,738 PPC genes used for eQTL mapping.

#### 5.3 ATAC-seq

All ATAC-seq samples were processed in a uniform manner using the same procedure as the ENCODE (https://github.com/ENCODE-DCC/atac-seq-pipeline) (Table S3). Illumina adapters were removed from the reads using cutadapt^62^. Reads were aligned using BWA MEM^63,64^ (https://bio-bwa.sourceforge.net/bwa.shtml) onto the hg38 human reference genome from UCSC (https://hgdownload.soe.ucsc.edu/goldenPath/hg38/bigZips/). Multi-mapped reads were randomly assigned using ENCODE’s custom script (assign_multimappers.py) (https://github.com/ENCODE-DCC/atac-seq-pipeline). Using samtools^59^, reads that were either unmapped, not in primary alignment, failed Illumina QC metrics, or had an unmapped mate were removed (samtools view -F 1804). Properly paired reads with mapping quality ≥ 30 were retained (samtools view -f 2 -q 30). Duplicates were marked by Picard and then removed with samtools. Mitochondrial reads were also excluded from downstream analyses. Filtered BAM files were converted to bed files (bedtools bamtobed) and shifted for Tn5 bias, and then used to call narrow peaks using MACS2^65^ with ENCODE default parameters (https://github.com/ENCODE-DCC/atac-seq-pipeline): –shift 75 –extsize 150 -q 0.01 –nomodel -B –SBMR –keep-dup all. Peaks overlapping blacklisted regions were removed. ATAC-seq samples were examined for quality and excluded if they did not pass one of the following metrics: 1) non-redundant fraction (NRF) > 0.9; 2) PCR-bottlenecking coefficient 1 (PBC1) > 0.9; 3) PCR-bottlenecking coefficient 2 (PBC2) > 3; 4) percent of mapped reads > 0.95; 5) fraction of reads in peaks (FRIP) > 10; 6) TSS enrichment (TSSE) > 4, and 7) matched via a sample identity check to the correct subject with PI_HAT > 0.90 (Figure S8). Across all ATAC-seq samples, the number of read pairs passing filters ranged from 6.4 million to 46.2 million with an average of 31.4 million.

To identify consensus peaks for each tissue, we selected high quality reference samples using the following filters: 1) 25 < FRiP < 45; 2) 5 < TSSE < 25; and 3) 75,000 < number of peaks < 200,000) from unrelated individuals in different families with two or more individuals in the iPSCORE collection (Figure S8). If multiple samples from the same family passed these filters, we selected the sample with the highest TSSE, which resulted in 24 iPSC, 24 PPC, and 23 CVPC reference samples. For each reference sample, we removed short peaks (<150 bp), then concatenated and merged peaks across all the reference samples for each tissue, resulting in 208,581 iPSC, 289,980 PPC, and 278,471 CVPC reference peaks used for downstream analyses. For each ATAC-seq sample, we used featureCounts^66^ (v2.0.6) to count the number of reads in each reference peak.

For caQTL mapping, for each dataset, we first TMM-normalized the reference ATAC-seq peak counts across samples using the *calcNormFactors* and *cpm* functions in in the edgeR package v3.38.4. We then removed ATAC-seq peaks on sex chromosomes or with low accessibility (TMM < 1 in at least 20% of the samples), resulting in 172,343 iPSC, 204,129 CVPC, and 193,428 PPC ATAC-seq peaks used for caQTL mapping (Table S7).

#### 5.4 H3K27ac ChIP-seq

All H3K27ac ChIP-seq data was processed in a uniform manner using the same procedure as the ENCODE (https://github.com/ENCODE-DCC/chip-seq-pipeline2) (Table S4). Illumina adapters were removed from the reads using cutadapt^62^. Reads were aligned using BWA MEM onto the hg38 human reference genome downloaded from UCSC (https://hgdownload.soe.ucsc.edu/goldenPath/hg38/bigZips/). Properly paired reads were retained (samtools view -f 2). Using samtools unmapped and non-uniquely mapped reads were removed (samtools view -F 1804), and PCR duplicates were marked with Picard and removed (samtools - F 1804). BAM files for each sequencing run were merged by library and then used to call narrow peaks using MACS2 with ENCODE default parameters: --shift −75 --extsize 150 -p 0.01 –nomodel -B --SPMR --keep-dup all. H3K27ac ChIP-seq samples were examined for quality and excluded if they did not pass one of the following metrics: 1) > 20 million usable fragments; 2) NRF > 0.9; 3) PBC1 > 0.9; 4) PBC2 > 10; and 5) FRIP > 1; and 6) matched via a sample identity check to the correct subject with PI_HAT > 0.90 (Figure S9). Across all H3K27ac ChIP-seq samples, the number of read pairs passing filters ranged from 10.0M to 64.6M with an average of 27.2M.

To identify CVPC H3K27ac ChIP-seq consensus peaks, we selected 29 high-quality reference samples using the following filters: 1) 5 < FRiP < 10; 2) 40M < number of reads passing filters < 100M; and 3) 25,000 < number of peaks < 75,000) from unrelated individuals in 29 families (Figure S9). If multiple samples from the same family passed these filters, we selected the sample with the highest FRiP. Since there were only 43 iPSC H3K27ac ChIP-seq samples, we did not consider sample quality to establish consensus peaks and selected 28 iPSC reference samples from 28 families. For families that had multiple individuals with iPSC H3K27ac ChIP-seq samples, we selected the sample with the highest FRiP. For each reference sample, we filtered peaks (500 bp > peak length > 5000 bp), then concatenated and merged peaks across all the reference samples in the corresponding tissue, resulting in 68,284 iPSC, and 64,135 CVPC reference peaks used for downstream analyses. For each H3K27ac ChIP-seq sample, we used featureCounts^66^ v2.0.6 to count the number of reads in each reference peak.

For haQTL mapping, for each dataset we first TMM-normalized the reference ChIP-seq peak counts across samples using the *calcNormFactors* and *cpm* functions in the edgeR^61^ package v3.38.4. We then removed ChIP-seq peaks on sex chromosomes or with low accessibility (TMM < 1 in at least 20% of the samples), resulting in 66,251 iPSC, and 60,543 CVPC ChIP-seq peaks used for haQTL mapping.

#### 5.5 Sample Identity

Sample identity was performed as previously described^2,4,6,7,12,17^. Briefly, genotypes were called from BAM files of each molecular dataset for common variants with minor allele frequency (MAF) > 45% and < 55% using bcftools^59^ mpileup and call, and then compared to WGS genotypes using plink --genome^67^, which calculates IBD between each pair of samples. Samples that matched the correct subject with PI_HAT > 0.90 passed sample identity check.

### 6. Analyses

#### 6.1 Characterizing ATAC-seq peak chromatin accessibility and H3K27ac ChIP-seq histone acetylation

##### 6.1.1 Comparing chromatin accessibility and histone acetylation across tissues

Independent consensus ATAC-seq peaks and H3K27ac ChIP-seq peaks were defined for each tissue and molecular dataset, therefore we could not use them to compare chromatin accessibility or histone acetylation across tissues. To create global consensus ATAC-seq peaks to compare all 391 ATAC-seq samples, we concatenated the 172,343 iPSC, 193,428 PPC, and 204,129 CVPC consensus ATAC-seq peaks, then sorted and merged them, resulting in 348,429 global consensus ATAC-seq peaks. Likewise, we created 101,393 global consensus ChIP-seq peaks by concatenating, sorting, and merging the 68,284 iPSC, and 64,135 CVPC consensus ChIP-seq peaks. We applied featureCounts^66^ (v2.0.6; as described in **5.3 ATAC-seq**) to count the number of reads in 391 ATAC-seq samples and 143 ChIP-seq samples, using the corresponding global consensus peaks. The counts were TMM-normalized, using edgeR^61^ (v3.38.4; as described in **5.3 ATAC-seq** and **5.4 H3K27ac ChIP-seq**). A PC analysis was performed on the top 2000 most variable global consensus peaks for ATAC-seq and ChIP samples, independently. A UMAP dimensionality reduction was performed on the top 9 PCs for ATAC and the top 10 PCs for ChIP samples (Figure S2a-b).

##### 6.1.2 Identification of tissue-specific and shared ATAC-seq peaks

We performed pairwise bedtools intersect using the consensus ATAC-seq peaks from the three tissues to identify peaks that were only present in one tissue. An ATAC-seq peak from a given tissue that overlapped (≥ 1 bp) at least one ATAC-seq peak from a different tissue was considered “Shared” (Figure S2c).

##### 6.1.3 ATAC-seq Peak Transcription Factor Predictions

The TOBIAS^22^ algorithm leverages the distribution of reads across the genome for a given sample, therefore to profile TF occupancy, we ran TOBIAS to predict binding at 1,147 motifs across ATAC-seq peaks for each tissue, independently. We first merged BAM files for the reference samples used to establish reference peaks for each tissue. We followed the standard workflow in the TOBIAS tutorial (https://github.com/loosolab/TOBIAS). Briefly, for each merged reference BAM file, we applied *ATACorrect* to correct for cut site biases introduced by the Tn5 transposase within the ATAC-seq peaks, using the following parameters: --genome hg38.fa (https://hgdownload.soe.ucsc.edu/goldenPath/hg38/bigZips/) and –blacklist hg38-blacklist.v2.be (https://github.com/Boyle-Lab/Blacklist/blob/master/lists/hg38-blacklist.v2.bed.gz). Next, we calculated footprints scores with *ScoreBigwig*, using the corresponding narrowPeak file for each tissue as input. To identify the predicted transcription factor binding sites, we ran *BINDetect* with 746 motifs from JASPAR^23^ (2020 version) and 401 HOCOMOCO^24^ v11 TF motifs independently, using hg38 fasta file and respective narrowPeak file as the genome and regions. For downstream analyses, we used 747 predicted TFBSs (the 746 JASPAR motifs and the NANOG motif from HOCOMOCO v11 database because it was not present in JASPAR; Figure 2a). The tables for predicted TFBSs at JASPAR and HOCOMOCO motifs are deposited on Figshare.

##### 6.1.4 Enrichment of TFBSs in tissue-specific and shared ATAC-seq peaks

We performed Fisher’s Exact tests to calculate the enrichment of TFBSs in tissue-specific and shared ATAC-seq peaks. For the three sets of tissue-specific ATAC-seq peaks, we calculated the differential TFBS enrichment relative to the corresponding shared ATAC-seq peaks, using peaks with at least one TFBS. For example, of the 66,370 tissue-specific and 105,705 shared ATAC-seq peaks in the iPSCs (Figure S2c), we used the 5,110 tissue-specific and 40,632 shared peaks that were bound by at least one TFBS to calculate enrichment, using the shared peaks as background. We calculated TFBS enrichment in “Shared” ATAC-seq peaks for each tissue independently. For each tissue we performed Fisher’s Exact tests on the set of shared ATAC-seq peaks in that tissue with at least one TFBS, using the tissue-specific peaks as background. For plot legibility, we plotted the log2 of the average Odds Ratios from the three Fisher’s Exact tests in Figure 2a.

#### 6.2 Quantitative Trait Loci (QTL) Mapping

##### 6.2.1 WGS Variant Selection

For all QTL analyses, we used single nucleotide polymorphisms (SNPs) that met the following criteria across the 273 individuals (see **5.1 WGS**): 1) passed Illumina QC; 2) in Hardy-Weinberg equilibrium (p > 0.000001); 3) genotyped in at least 99% of the individuals; and 4) had MAF > 0.05. After filtering, 5,536,303 SNPs remained.

##### 6.2.2 Kinship Matrix

To account for genetic relatedness between samples, we performed LD pruning on the 5,536,303 variants using plink^67^ 1.90b6.21 (-indep-pairwise 50 5 0.2). We then used the 323,697 LD-pruned variants to construct a kinship matrix for the 273 iPSCORE individuals using plink^67^ 1.90b6.21 (– make-rel square).

##### 6.2.3 Global Ancestry: Genotype Principal Component Analysis

We performed genotype principal component analysis (PCA) across all 273 individuals in the iPSCORE Collection. First, we intersected the 323,697 LD-pruned variants above with 1000 Genomes^53–55^ single nucleotide polymorphisms (SNPs). Then, using plink^67^ 1.90b6.21 (--pca-cluster-names AFR EUR AMR EAS SAS –pca), we performed PCA excluding 1000 Genome subjects without super-population information. We determined that the first five genotype PCs for QTL analysis were sufficient and captured the majority of the variability that was due to global ancestry (Figure S1c). The ancestries reported for the 221 subjects in this study (Figure S1a-b), were assigned in a previous study describing the iPSCORE Collection^7^.

##### 6.2.4 PEER Factor Calculation

To account for hidden technical and biological confounders that influence gene expression variability, we used Probabilistic Estimation of Expression Residuals^68^ (PEER) to estimate a set of latent factors for each tissue (iPSC, CVPC, PPC) and molecular data type (RNA-seq, ATAC-seq, H3K27ac ChIP-seq). We used the top 2,000 most variable genes/peaks to calculate a maximum number of PEER factors that is equivalent to ∼25% of the samples (for instance, for the CVPC RNA-seq dataset of 178 samples, we set the maximum number of PEER factors to 50), as recommended by the original developers^68^. As previously described^2,4^, to determine the number of PEER factors to use for QTL discovery, we piloted QTL mapping on a random set of 1,000 genes or 4,000 peaks using varying numbers of PEER factors as covariates (listed in Table S2; Table S3; Table S4) and selected the least number of PEERs that resulted in maximum eGene, caPeak, and haPeak discovery (Figure S10). For eQTLs, we used 51, 35, and 22 PEER factors for iPSC, CVPC, and PPC, respectively as covariates. For caQTLs, we used 28, 28, and 20 PEER factors for iPSC, CVPC, and PPC, respectively, as covariates. For haQTLs, we used 8 and 19 PEER factors for iPSC and CVPC, respectively, as covariates. We found that the variance captured by PEER factors was correlated with known biological and technical factors recorded for each sample (Figure S11). In particular, we observed that the top PEER factors across all the molecular data types were highly correlated with sequencing quality, differentiation efficiency, and sex.

##### 6.2.5 QTL Covariates

For all QTL analyses, we included the following as general covariates: sex, iPSC passage number, the first five genotype PCs to control for global ancestry, and PEER factors to account for hidden confounders of molecular phenotype variability (see **6.2.4 PEER Factor Calculation**).

##### 6.2.6 QTL Mapping

QTL analysis was performed on each of the 8 iPSCORE molecular datasets independently using a linear mixed model (LMM) with the kinship matrix as a random effect to account for the genetic relatedness between samples. First, using rank and qnorm in R (v4.2.1), we inverse normal transformed the TMM gene expression/peak accessibility or acetylation values across the samples. Genes within 1 Mb and peaks within 100 kb of the MHC region^69^ (chr6:28,510,120-33,480,577) were removed due to the complex LD structure in the interval. For the elements (i.e. genes and peaks) outside the MHC region, we used bcftools^59^ query to obtain the genotypes for all the variants within 1 Mb for genes or 100 kb for ATAC-seq peaks and H3K27ac ChIP-seq peaks. Then, we applied the scan function in limix (v3.0.4) (https://github.com/limix/limix) to run the following linear mixed model:

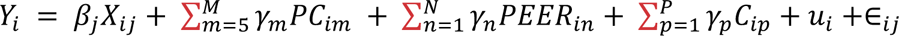

Where *Y_i_* is the normalized expression value for sample *i*, *β_j_* is the effect size (fixed effect) of SNP *j*, *X_ij_* is the genotype of sample *i* at SNP *j*, *M* is the number of genotype principal components used (*M* = 5 for all QTL analyses), *γ_m_* is the effect size of the *m*th genotype principal component, *PC_im_* is the value of the *m*th genotype principal component for the individual associated with sample *i*, *N* is the number of PEER factors (See **6.2.4 PEER Factor Calculation**), *γ_m_* is the effect size of the *n*th PEER factor, *PEER_in_* is the value for the *n*th PEER factor for sample *i*, *P* is the number of covariates used (*P* = 1 for all QTL analyses corresponding to the iPSC passage number), *γ_p_* is the effect size of the *p*th covariate, *C_ip_* is the value for the *p*th covariate for sample *i*, *u_i_* is a vector of random effects for the individual associated with sample *i* defined from the kinship matrix, and ∈_*ij*_ is the error term for individual *i* at SNP *j*.

##### 6.2.7 FDR correction

We used a two-step procedure described in Huang et al.^70^, which first corrects at the gene or peak level and then at the genome-wide level. First, we performed FDR correction on the p-values of all independent variants tested for each gene or isoform using eigenMT^71^, which considers the LD structure of the variants. Then, we extracted the lead variant for the QTL for each gene or peak based on the most significant FDR-corrected p-value. If more than one variant had the same FDR-corrected p-value, we selected the one with the largest absolute effect size as the lead variant for the QTL. For the second correction, we performed FDR-correction on all lead variants using Benjamini-Hochberg (q-value). We considered only QTLs with q-value < 0.05 as significant (Table S6; Table S7; Table S8).

##### 6.2.8 Conditional QTL mapping

To identify additional independent QTL associations for a gene or peak (i.e., conditional QTLs), we performed stepwise regression analysis in which we re-performed QTL analysis with the genotype of the lead variant for the QTL as a covariate. We repeated the procedure to discover up to three conditional associations. For each iteration, we performed the two-step procedure described above and considered conditional eQTLs with q-values < 0.05 as significant.

#### 6.3 QTL characterization

##### 6.3.1 QTL fine-mapping

For each QTL, we used the SNP effect sizes and allele frequencies as input to the *finemap.abf* function from the coloc^72^ R package (v5.2.2) to generate posterior probabilities (PP) for all tested SNPs. We selected the SNP with the highest PP for each QTL as the lead fine-mapped variant. Of the 79,081 QTLs, 68,745 had a fine-mapped variant with a PP ≥ 1%.

##### 6.3.2 Primary versus conditional QTLs proximity to qElements

For each QTL, we calculated the minimum distance between the 68,745 QTLs that had a fine-mapped variant with a PP ≥ 1% and the corresponding Peak (Figure S5). For eGenes, we considered the TSS to be the region 1 kb ± start codon.

##### 6.3.3 eGene discovery rate

To examine the relative power of identifying eQTLs, we compared the three iPSCORE EDev-like tissues to the 49 tissues in the GTEx Consortium^1^ and showed that they had similar eGene discovery rates (Figure S4).

##### 6.3.4 caPeaks enriched for being near eGenes

Because we identified caQTLs within 100 kb of an ATAC-seq peak, we tested the enrichment of caPeaks within a 100 kb window around any eGene compared to within a 100 kb window around any expressed gene without an eQTL.

##### 6.3.5 caQTL Discovery Rate by Peak Width

We examined the caQTL discovery rate by ATAC-seq peak width for iPSCs, CVPCs, and PPCs and showed that caQTL discovery rates were associated with the ATAC-seq peak width (Figure S12).

##### 6.3.6 Chromatin State Enrichment

We obtained ChromHMM chromatin states in the hg38 build for iPSC line 18a (BSS00737) and embryonic stem cell-derived cardiac muscle (BSS00171) from the EpiMap Repository (https://compbio.mit.edu/epimap). We collapsed the 18 chromatin states into 5 categories Promoter (which included TssA, TssBiv, TssFlnkU, TssFlnkD, TssFlnk), Enhancer (EnhA1, EnhA2, EnhG1, EnG2, EnhWk, EnhBiv), Repressed (ReprPC, ReprPCWk, Het, ZNF/Rpts), Transcribed (Tx, TxWk), and Quiescent (Quies). We annotated the lead variant, regardless of significance, from each QTL test (252,529 iPSC and 311,814 CVPC) across the three molecular phenotypes with the 5 collapsed chromatin states that they overlapped in their corresponding EpiMap tissue using *bedtools intersect*. We concatenated the annotations from both tissues, then, for each condition (i.e. primary, conditional 1, conditional 2, and conditional 3) and molecular phenotype, we performed two-sided Fisher’s Exact tests to test the enrichment of lead variants from significant QTLs in each of the 5 collapsed chromatin states, using the lead variants (q-value > 0.05) from non-qElements (i.e., genes and peaks without a QTL) as background.

##### 6.3.7 TFBS Enrichment in CVPC caPeaks and haPeaks

To characterize TF binding in regulatory elements affected by variation, we analyzed the CVPC ATAC-seq and H3K27ac ChIP-seq datasets. We first classified CVPC ATAC-seq peaks based on whether they were a caPeak and whether that caPeak overlapped (≥ 1 bp) a CVPC haPeak. Using only ATAC-seq peaks with at least one predicted TFBS, we performed a Fisher’s Exact test to calculate the enrichment of the 748 TFBS motifs (see **6.1.3 ATAC-seq Peak Transcription Factor Predictions)** in caPeaks, caPeaks-haPeaks and CVPC ATAC-seq peaks without a caQTL (no QTL).

#### 6.4 Identification of QTL Modules

##### 6.4.1 Intra-Tissue QTL Colocalization

To identify primary QTLs within the same tissue across the three molecular data types that shared causal variants, we performed pairwise Bayesian colocalization using the *coloc.abf* function in *coloc*^72^ v5.2.2 between the six pairwise combinations of QTLs (eQTL-eQTL, caQTL-caQTL, haQTL-haQTL, caQTL-eQTL, haQTL-eQTL, and caQTL-haQTL; PPCs do not have H3K27ac ChIP-seq, therefore combinations with haQTLs were not analyzed). First, for each tissue, we created a bed file containing the coordinates of windows tested for each QTL (i.e. for each eGene, we included 1 Mb upstream and downstream; for each caPeak, we included 100 kb upstream and downstream; and for each haPeak coordinates, we included 100 kb upstream and downstream, then we identified overlapping regions between QTLs. We then performed colocalization (*coloc.abf*) on each QTL pair, considering only pairs that shared at least 50 biallelic SNPs. We used the following filters to determine if two QTL signals were colocalized; 1) posterior probability of the signal being shared (PP.H4) ≥ 80%, 2) top SNP nominal p-value < 5 x 10^−5^ in both QTLs, and 3) causal posterior probability for the top SNP ≥ 1%.

##### 6.4.2 QTL Module Assignments

For each tissue, we loaded each pair of colocalized primary QTLs as edges into an *igraph*^73,74^ (v1.3.2) (https://igraph.org) network. To identify modules, we clustered the colocalized QTL networks using the *cluster_louvain* function, then divided each module into a subgraph, using the induced subgraph function, and calculated each subgraph’s modularity, using the *modularity* function. If a subgraph exhibited high modularity (> 0.3), the subgraph was recursively clustered using the *cluster_louvain* function and divided into multiple modules. In total, there were 4,402 QTL modules composed of two or more QTLs, and 49,616 singleton QTLs that did not colocalize with another QTL (Table S9; Table S10).

#### 6.5 GWAS associations with QTLs

##### 6.5.1 GWAS Traits

To estimate the enrichment of heritability for developmental and adult GWAS traits in ATAC-seq and ChIP-seq peaks in the three early developmental-like tissues, we considered the following 17 traits: fasting glucose, chronic ischemic heart disease, birth weight, type 2 diabetes, LDL direct levels, HDL cholesterol levels, angina pectoris, type 1 diabetes, body mass index, ventricular rate, pulse rate, atrial fibrillation and flutter, QRS duration, and acute myocardial infarction, and childhood obesity. GWAS summary statistics for the 17 traits were downloaded from the 1) UK Biobank (https://pan.ukbb.broadinstitute.org/downloads/index.html) for angina pectoris, atrial fibrillation, body mass index, HDL cholesterol, ischemic heart disease, LDL direct, myocardial infarction, pulse rate, QRS duration, and ventricular rate, 2) the Early Growth Genetic Consortium (http://egg-consortium.org/) for childhood obesity^75^ and birth weight^76^, 3) the Meta-Analyses of Glucose and Insulin-related Traits Consortium (http://magicinvestigators.org/downloads/) for fasting glucose^77^, 4) a previous study^45^ for type 1 diabetes, 5) the DIAGRAM Consortium (https://diagram-consortium.org) for type 2 diabetes^78^, and 5) the Edinburgh Data Share (https://datashare.ed.ac.uk/handle/10283/3209; https://datashare.ed.ac.uk/handle/10283/3599) for longevity GWAS for parental lifespan^79^ and aging^80^. All traits are listed in Table S11, along with their study sources. All of the data, except for type 1 diabetes, were provided in hg19 coordinates. To convert coordinates from hg19 to hg38, we used the liftOver software downloaded from UCSC (https://genome-store.ucsc.edu/). Then, we sorted and indexed each GWAS summary statistics file using tabix^59^.

##### 6.5.2 GWAS-QTL Colocalization

For each primary QTL identified in the 8 iPSCORE QTL datasets, we performed pairwise colocalization with GWAS variants (see **6.1.5 LD Score Regression** for list of GWAS summary statistics) using effect size and variance as input into the *coloc.abf* function in *coloc*^81^ (v5.2.2). For each GWAS-QTL colocalization, *coloc*^81^ outputs a causal PP for each variant that was tested during colocalization. We assigned a lead candidate causal variant for each GWAS-QTL pair by taking the variant with the highest causal PP. To determine whether a QTL colocalized with GWAS variants, we required that all of the following criteria were satisfied: 1) had at least 50 overlapping variants; 2) PP.H4 ≥ 80%; 3) the lead candidate causal variant is genome-wide significant for GWAS association (p-value ≤ 5 x 10^−8^); 4) the lead candidate causal variant is genome-wide significant for QTL association (p-value ≤ 5 x 10^−5^); and 5) the lead candidate causal variant had a PP ≥ 1%.

For QTL modules, we required that at least one of the QTLs in the module to colocalize with GWAS with all of the above criteria satisfied (Table S11). To assign a single lead candidate causal variant to each GWAS-colocalized QTL module, we assigned the variant with the highest causal PP among the QTLs that colocalized with the GWAS signal (PP.H4 ≥ 80%). As an example, for a module with four QTLs, two QTLs can colocalize with the GWAS signal, each having their own lead candidate causal variant (PP.H4 ≥ 80%). To assign a single candidate causal variant for the module, we assigned the one from the QTL that had the maximum causal PP.

##### 6.5.3 Fraction of GWAS Loci colocalized with QTLs

To determine the fraction of GWAS loci explained by QTLs, we calculated the number of independent genome-wide significant loci for each of the 17 GWAS studies. Specifically, we first filtered for variants that were above the genome-wide significant threshold of p < 5×10^−8^. Then, we applied LD pruning using plink^67^ with the following parameters: *--indep-pairwise 500 50 0.1*, where 500 is the variant count window, 50 is the step count, and 0.1 is the LD threshold. This command outputs a list of 6,371 independent variants (i.e., not in LD) that each represent an independent genome-wide significant GWAS loci. After identifying the 6,371 independent GWAS loci, we then sought to identify the subset that had colocalized to a QTL (either a module or singleton) (PP.H4 ≥ 80%). Using 1000 Genomes Phase 3 (Europeans only) as reference, we calculated LD between the lead candidate causal variants from the GWAS-QTL colocalization (see **6.5.2 GWAS-QTL Colocalization)** and the LD-pruned GWAS variants using *plink –tag-kb 500 –tag-kb 0.7*^67^. If the lead candidate causal variant was in high LD (r^2^ ≥ 0.7 within 500 kb) with an LD pruned GWAS variant, then we assigned the QTL module or singleton to that GWAS locus. For lead candidate causal variants absent from the reference panel, and therefore LD could not be calculated, we assigned the QTL module or singleton to the nearest GWAS locus. We observed that 65 of the 6,371 (1.0%) GWAS loci were in LD with multiple QTL modules or singletons from the same tissue (range 2-4 QTL modules/singletons per GWAS signal) (Table S11), which could reflect independent signals within the same GWAS locus or a limitation of coloc in assuming a single causal variant^72^.

##### 6.5.4 Enrichment of QTL Modules with GWAS Variants

We annotated each QTL based on their associated molecular phenotype and whether they were a module or a singleton (Table S10). For example, if a QTL module was composed of only caQTL(s) and eQTL(s), we annotated the module as “caQTL-eQTL module”. There were a total of ten categories: 1) caQTL-haQTL-eQTL module, 2) caQTL-haQTL module, 3) haQTL-eQTL module, 4) caQTL-eQTL module, 5) caQTL module, 6) eQTL module, 7) haQTL module, 8) caQTL singleton, 9) eQTL singleton, 10) haQTL singleton (Figure 4b). Enrichment of each of these categories for GWAS colocalization was calculated using a two-sided Fisher’s Exact test, where the contingency table consisted of two classifications: 1) if the QTL module or singleton corresponded to the category, and 2) if the QTL module or singleton colocalized with at least one GWAS trait. A category was considered enriched for GWAS colocalization if the Bonferroni-corrected p-value < 0.05.

#### 6.6 Identification of EDev-unique QTLs

To examine whether the primary QTLs discovered in the 8 iPSCORE QTL datasets were EDev-unique, we examined their overlap with the following publicly available adult QTLs: 1) 29,762 caVariants (p < 1×10^−5^) from QTLbase2^34^ (http://www.mulinlab.org/qtlbase) generated from adult tissues; 2) 244,563 haVariants (p < 1×10^−5^) from QTLbase2^34^ (http://www.mulinlab.org/qtlbase) generated from adult tissues; 3) 4,172 primary and conditional lead eQTLs from adult pancreatic islets^82^ using https://zenodo.org/records/3408356/files/PacreaticIslets_independent_exon_eQTLs.txt); 4) 2,948 lead caQTLs from adult pancreatic islets^33^; and 5) 276,116 primary and 408,175 conditional eQTLs from the GTEx database (version 8) for 49 adult tissues (https://www.gtexportal.org/home/downloads/adult-gtex#qtl). Because adult pancreatic islet e/caQTLs were available only in hg19 coordinates, we performed liftOver to hg38 using the liftOver R package^83^.

We considered an iPSCORE QTL as EDev-unique if their fine-mapped lead variant was not in LD with any adult QTL (r^2^ < 0.2 within 500 kb) regardless of molecular phenotype. If the QTL was in a module, we required that all QTLs in the module were not in LD with any adult QTL (r^2^ < 0.2 within 500 kb) to be considered EDev-unique. LD between variants was calculated with plink^67^ - -tag-r2 0.2 --tag-kb 500 using the 1000 Genomes Panel 3^53–55^ (Europeans) as reference. For fine-mapped lead variants that were not in the reference panel, and therefore LD could not be calculated, if the iPSCORE QTL was within 500 kb of an adult QTL, it was not considered as EDev-unique (Table S6; Table S7; Table S8). We identified 7,614 primary EDev-unique QTLs that we used for analyses.

## SUPPLEMENTAL FIGURES

**Figure S1.**
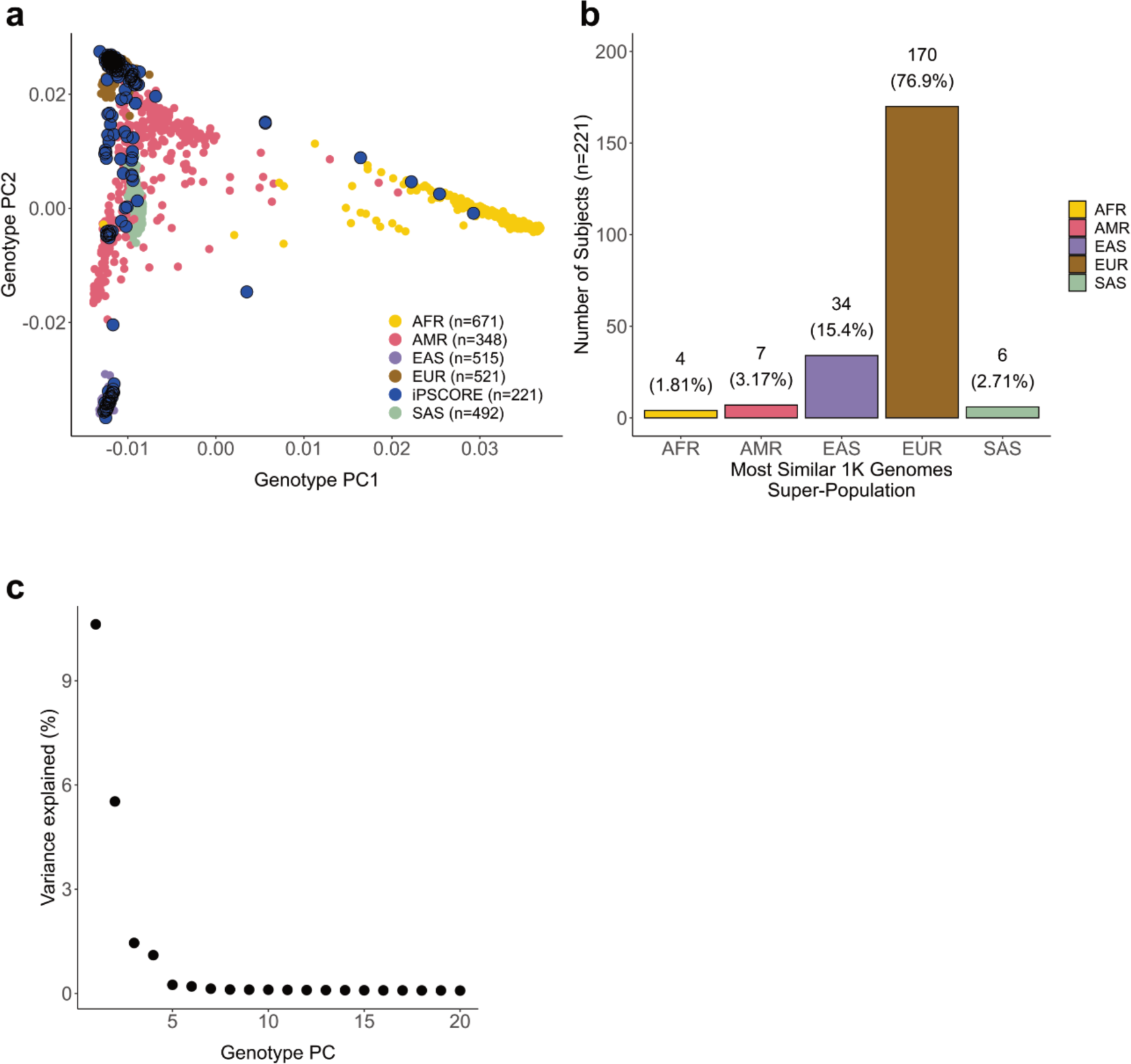
Genotype Principal Component Analysis (PCA) **a)** Scatter plot showing the distribution of the 221 iPSCORE subjects (blue) included in this study, across the five super-populations defined in the 1000 Genomes. The x-axis represents genotype principal component 1, and the y-axis represents genotype principal component 2. **b)** Distribution of the 221 iPSCORE subjects based on their closest matching super population as previously described^7^. **c)** Elbow plot showing the genotypic variance explained (y-axis) by each genotype principal component (x-axis). We found that genotype principal components 1-5 were sufficient to capture most of the genotypic variance explained by global ancestry and hence, will be used as covariates for QTL analyses.

**Figure S2.**
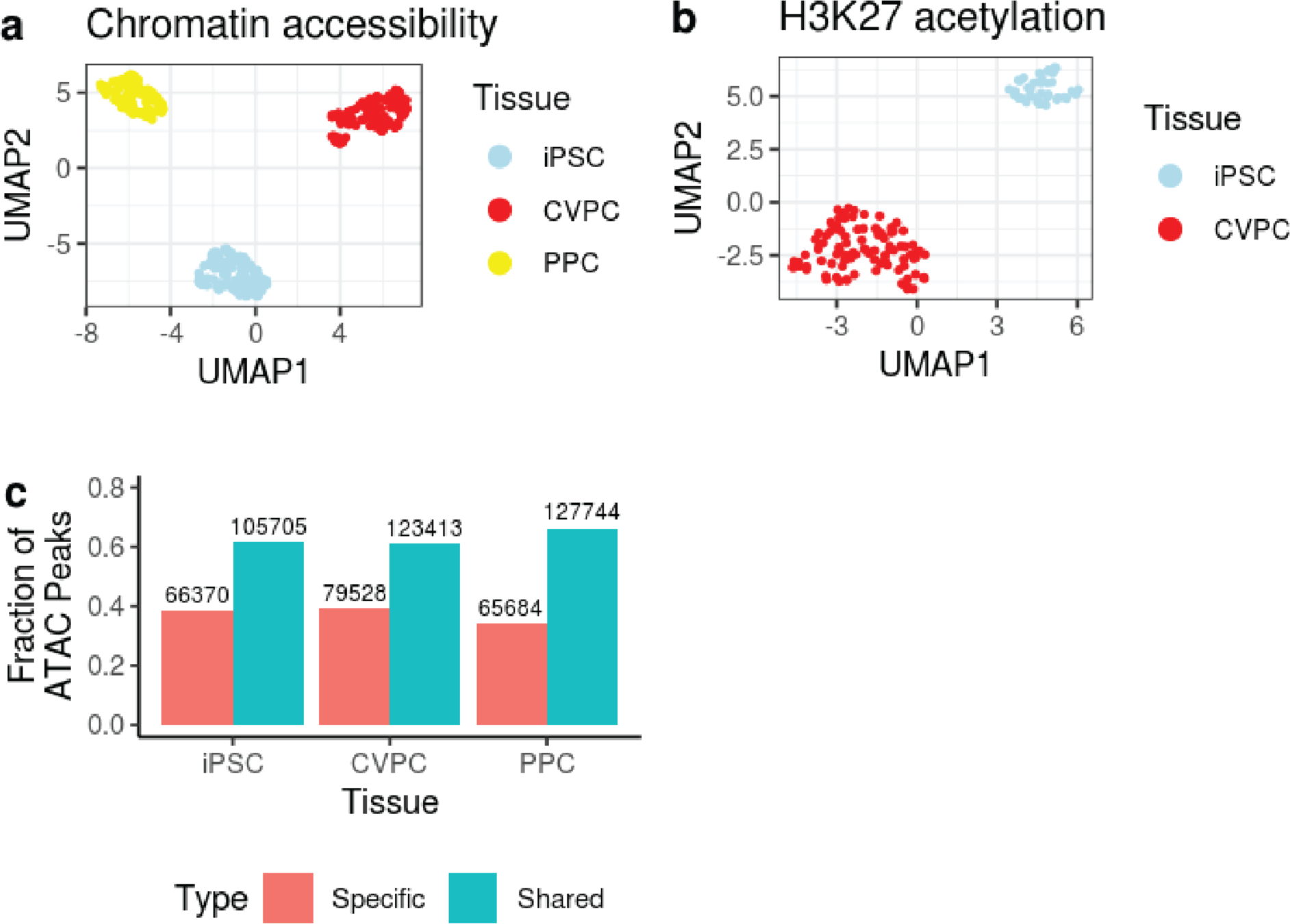
Characterization of iPSCORE Epigenomic Datasets. **a-b)** UMAP of chromatin accessibility of 391 ATAC-seq **(a)** and 144 histone acetylation of 144 H3K27ac ChIP-seq **(b)** samples from the iPSCORE Collection. Consensus peaks were curated for each dataset independently, and then merged to compare chromatin accessibility and histone acetylation across samples (See Methods). A UMAP analysis was performed using the top 2,000 most variable peaks. The samples cluster by tissue, indicating that the iPSCs and the derived fetal-like CVPCs and PPCs each have distinct regulatory landscapes. Each point represents an ATAC-seq sample colored by their corresponding tissue. **c)** Bar plot showing the fraction of ATAC-seq peaks that are tissue-specific or shared. We intersected the three independent consensus ATAC-seq peak sets to annotate peaks that were specific to one tissue or shared between at least two tissues. Across all three EDev-like tissues, approximately one-third of ATAC-seq peaks were tissue-specific (range = 33.9-39.2%) and approximately two-thirds were shared (range = 60.8-66.1%). The bar colors correspond to tissue-specific (red) and shared (blue) peaks. The labels at the top of each bar correspond to the number of ATAC-seq peaks. The x-axis contains the tissue labels, and the y-axis is the fraction of ATAC-seq peaks.

**Figure S3.**
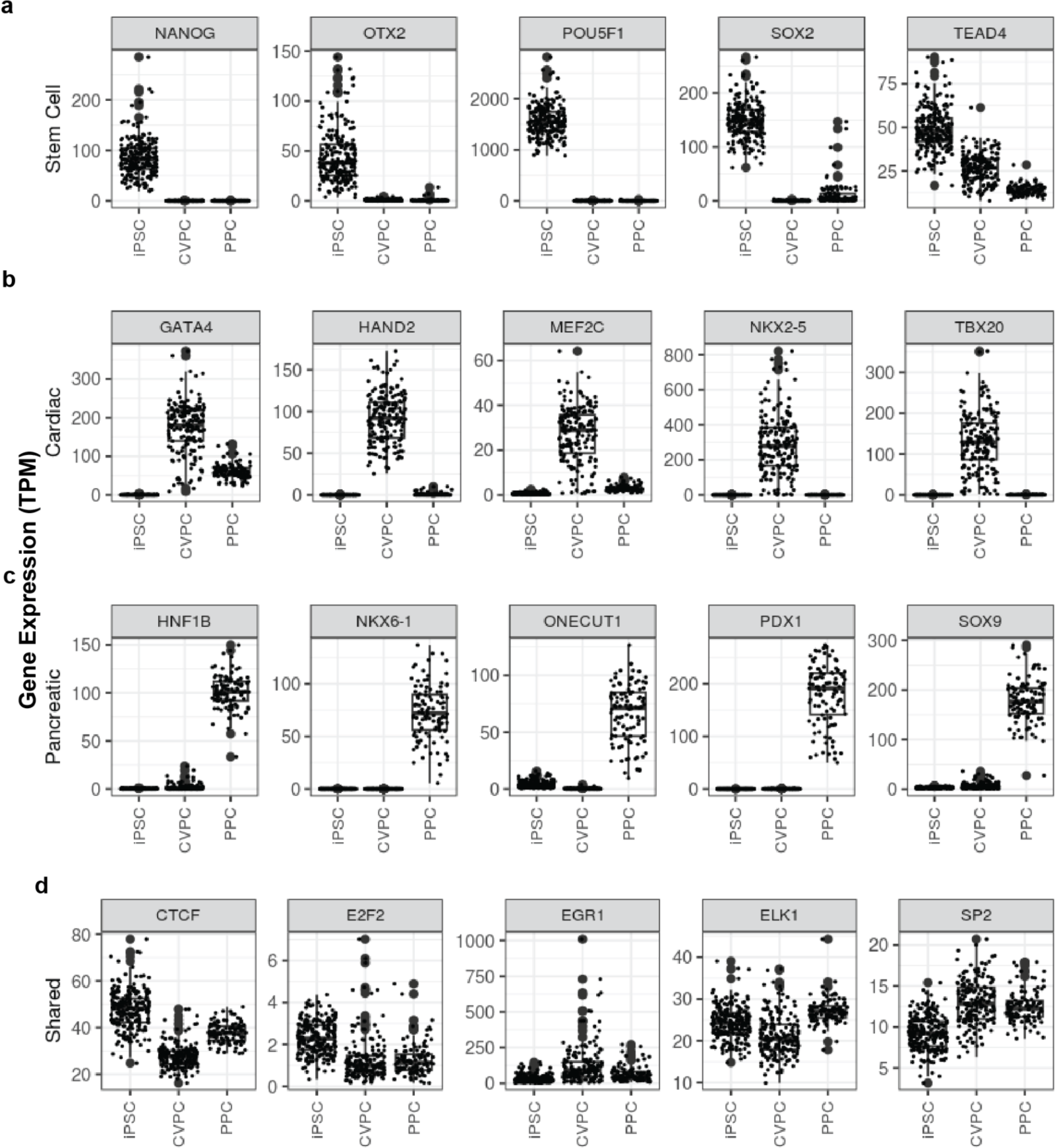
Expression of Tissue-specific TF Markers. **a-d)** Boxplots showing the expression of iPSC **(a)**, CVPC **(b)**, PPC **(c),** and shared (**d**) TF markers. Each point represents an iPSCORE RNA-seq sample from the three tissues (x-axis) and the expression of the corresponding TFs (y-axis). The tissue-specific TF markers exhibit differential expression relative to the other tissues.

**Figure S4.**
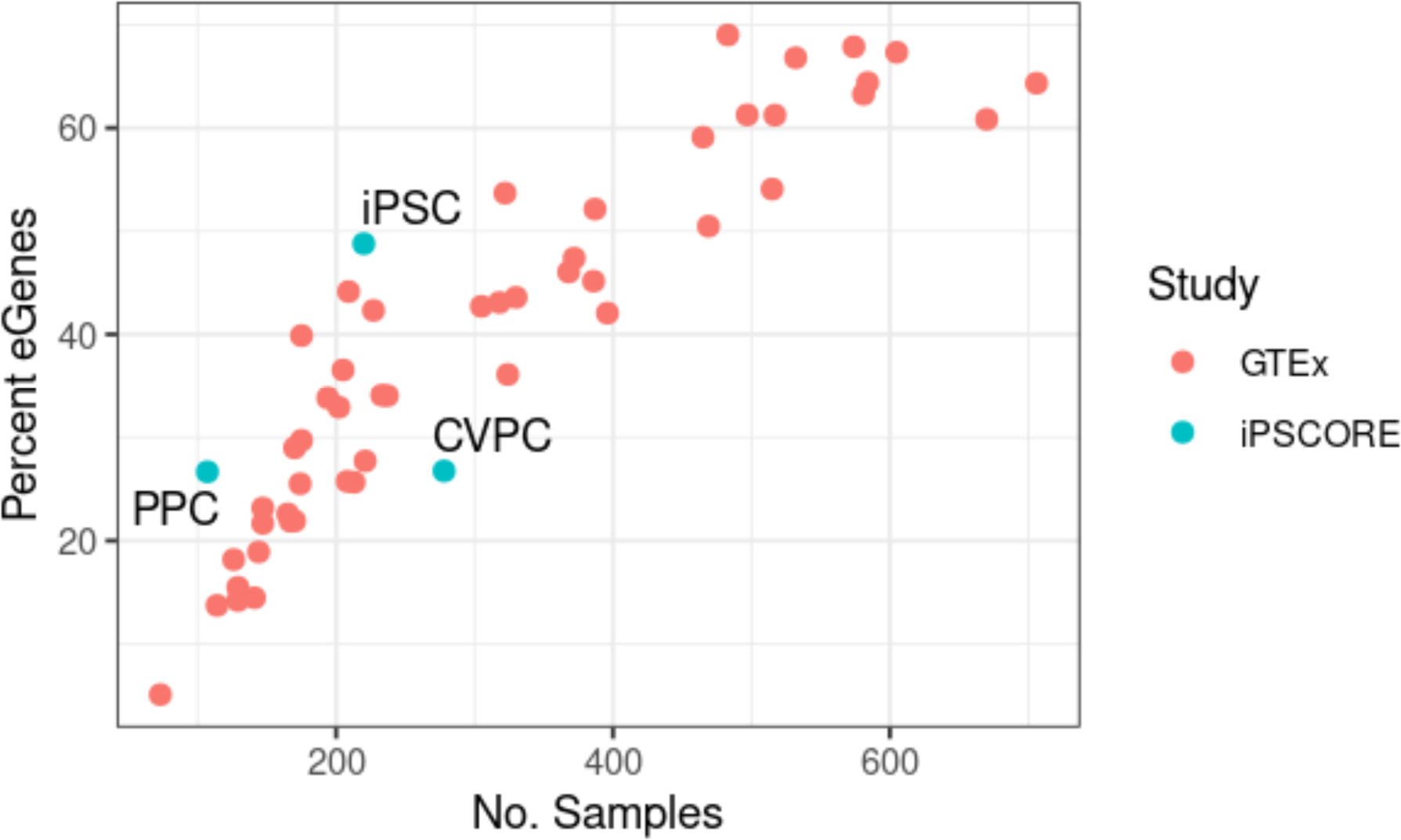
eGene Discovery Rate. Scatter plot showing the percentage of expressed genes that are eGenes in each tissue relative to the 49 GTEx adult tissues (version 8), as a function of sample size. These findings support that the eGene discovery rate in the three iPSCORE EDev-like tissues is similar to the 49 tissues in the GTEx Consortium^1^. Red points represent GTEx adult tissues while blue points represent the three iPSCORE EDev-like tissues in this study.

**Figure S5.**
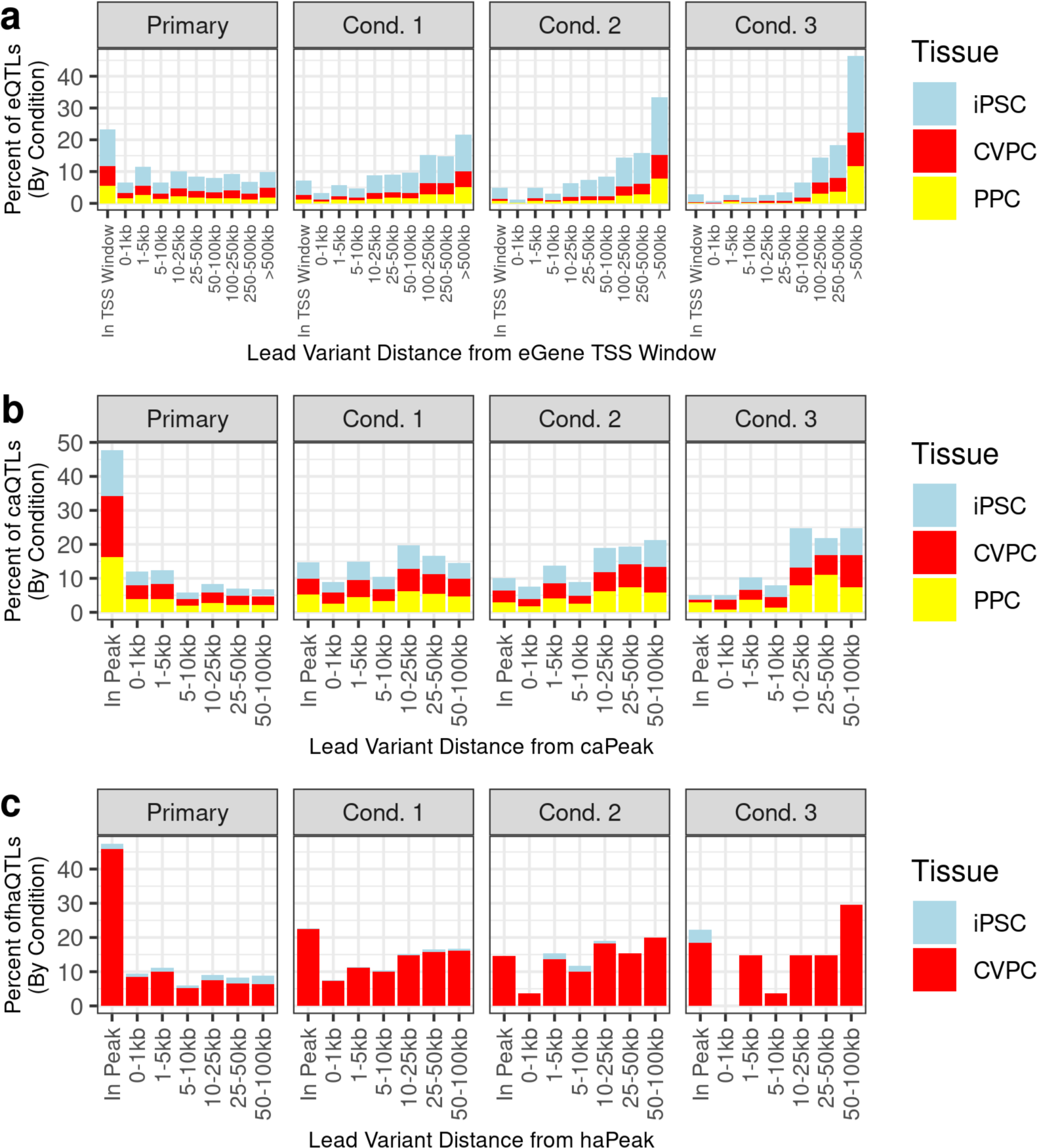
Conditional QTLs are more distal to their qElement. **a-c)** Bar plots demonstrating the minimum distance between fine-mapped variants with PP ≥ 1% in eQTLs (**a**), caQTLs (**b**), haQTLs (**c**) and the associated element for primary and conditional QTLs. The x-axis contains binned distance categories, and the y-axis is the percent of QTLs. The different panels correspond to primary and conditional QTLs. Each bar is colored by tissue (iPSC = “light blue”, CVPC = “red”, PPC = “yellow”).

**Figure S6.**
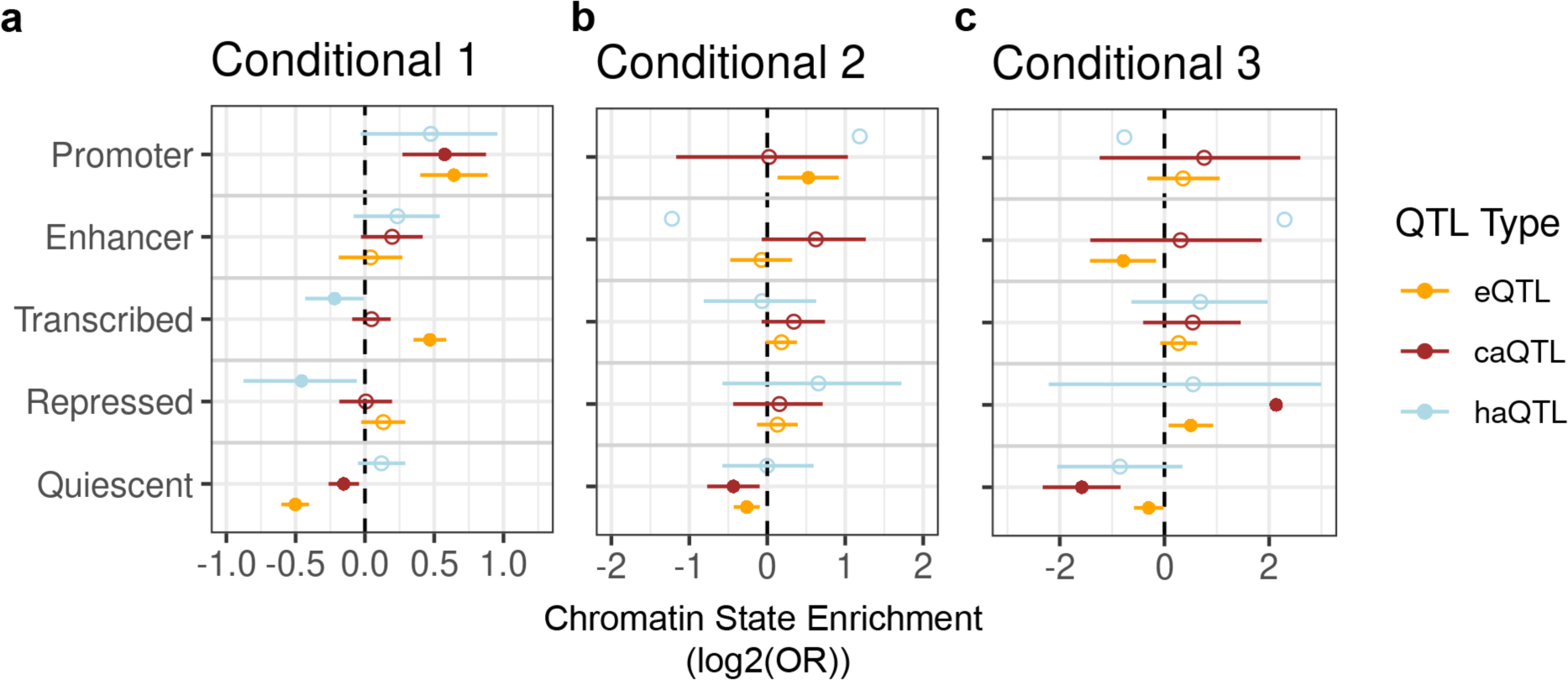
Conditional QTLs Chromatin State Enrichment. Plot showing the enrichment of conditional iPSC and CVPC eQTLs, caQTLs, and haQTLs in chromatin states. The x-axis is the enrichment log2(Odds Ratio) and the y-axis contains the five collapsed chromatin states. The points are colored by the QTL type (eQTL = “orange”, caQTL = “brown”, and haQTL = “light blue”). The whiskers represent the log2 upper and lower 95% confidence intervals. Significant enrichments are represented by filled circles and non-significant enrichments are represented by circles without a fill.

**Figure S7.**
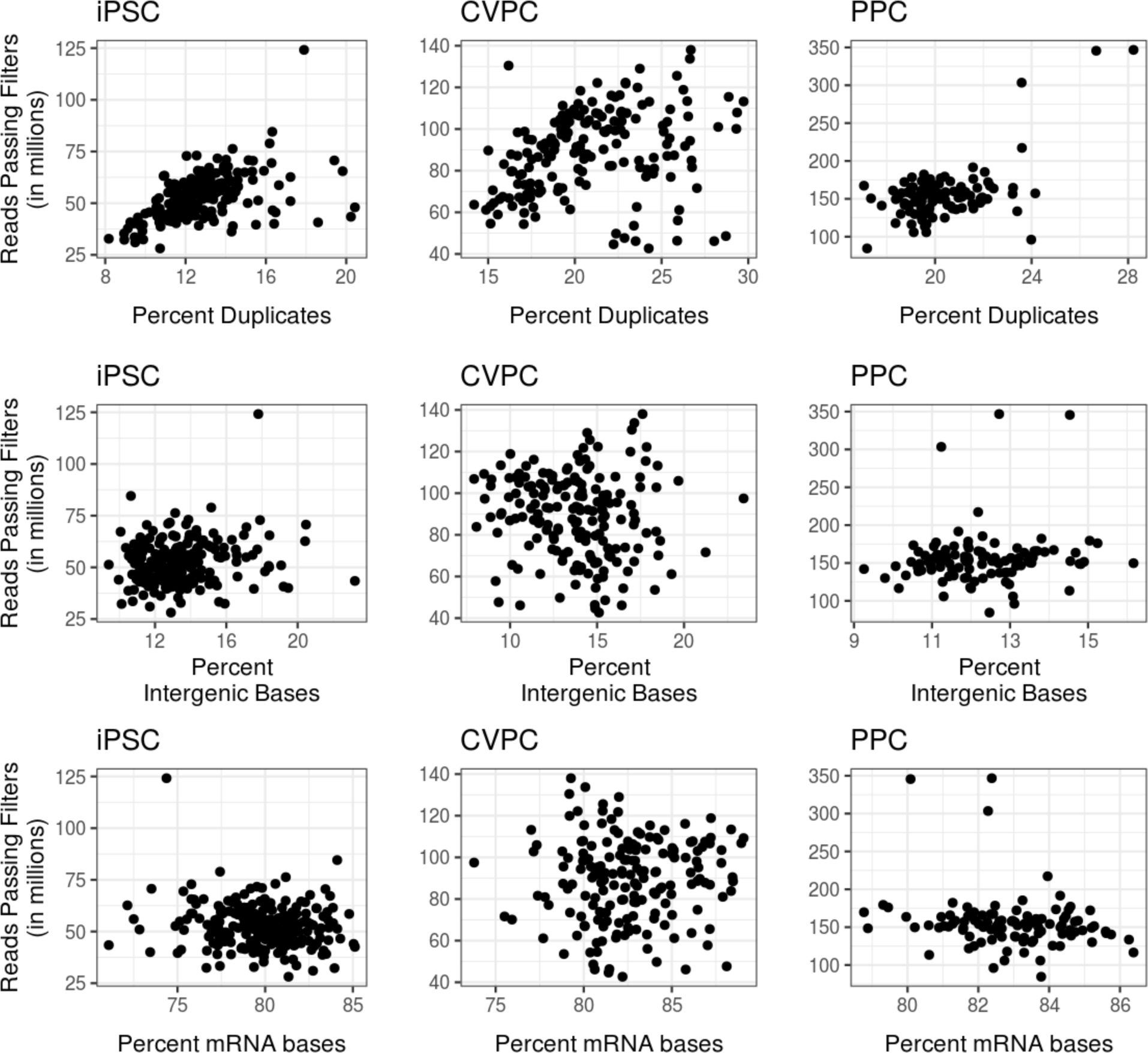
RNA-seq Quality Control. Scatter plots displaying the quality of the RNA-seq samples from iPSCs (n = 220), CVPCs (n = 178), and PPCs (n = 107). The plots show the number of reads passing filters (y-axis) against the percent of duplicate reads (row 1), the percent of intergenic bases (row 2), and the percent mRNA bases (row 3) for all 505 RNA-seq samples. All samples have been published in previous iPSCORE eQTL publications^2,4,6^, with the exception of 7 iPSC samples that were newly incorporated into this study.

**Figure S8.**
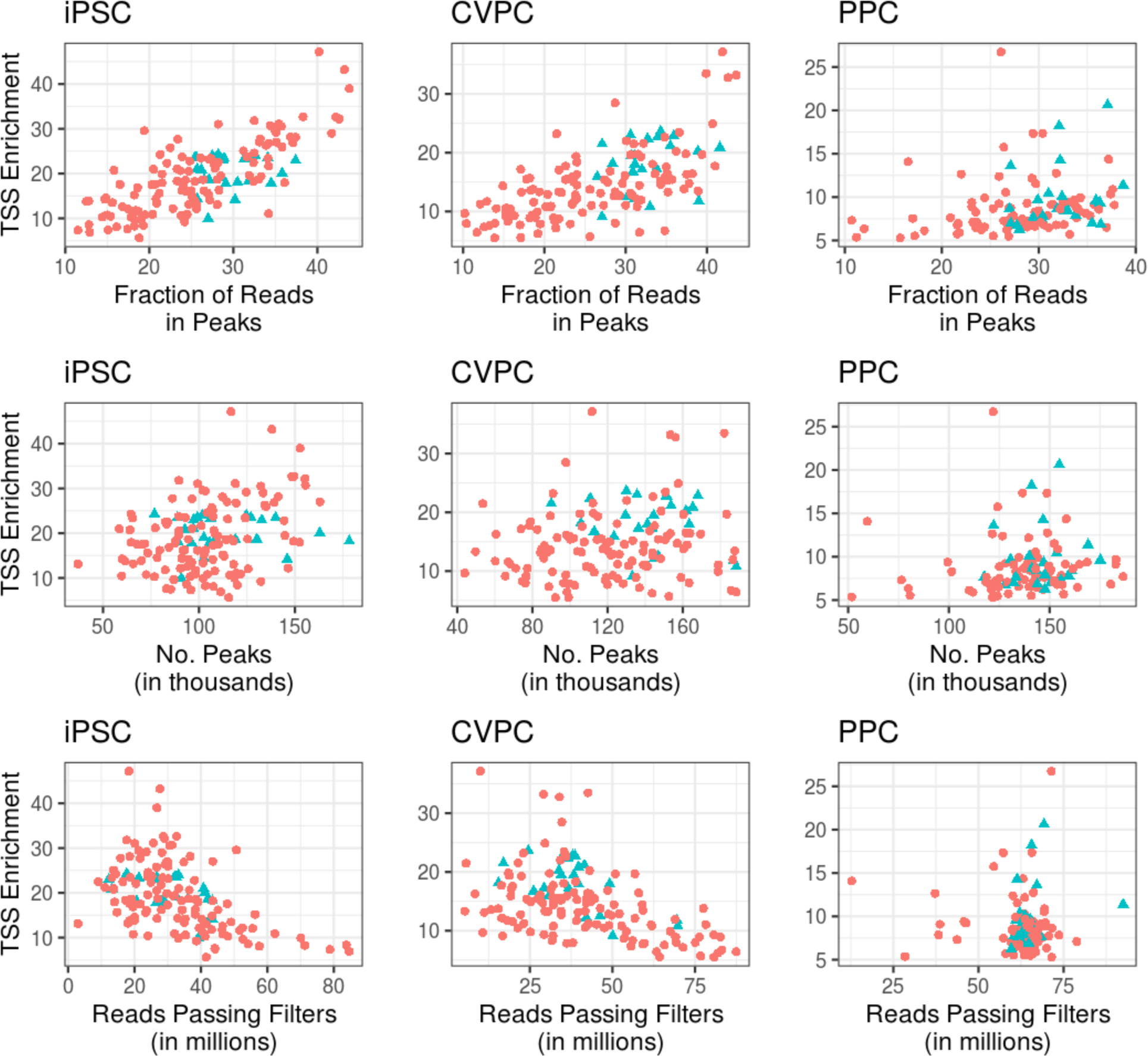
ATAC-seq Quality Control and Reference Sample Selection. Scatter plots displaying the quality of the ATAC-seq samples from all three tissues. The top row shows the transcription start site enrichment (TSSE; calculated by the ATACseqQC R package) of the samples plotted against the fraction of reads in peaks (FRiP), the middle row shows the TSSE and the number of ATAC-seq peaks per sample, and the bottom row shows the TSSE and the number of reads passing filters. We selected reference samples (blue triangles) based on QC metrics (See Methods) from unrelated individuals for each tissue (iPSC n=24; PPC n=24; and CVPC n=23) to establish a set of consensus peaks for the quantification of chromatin accessibility across the respective tissues.

**Figure S9.**
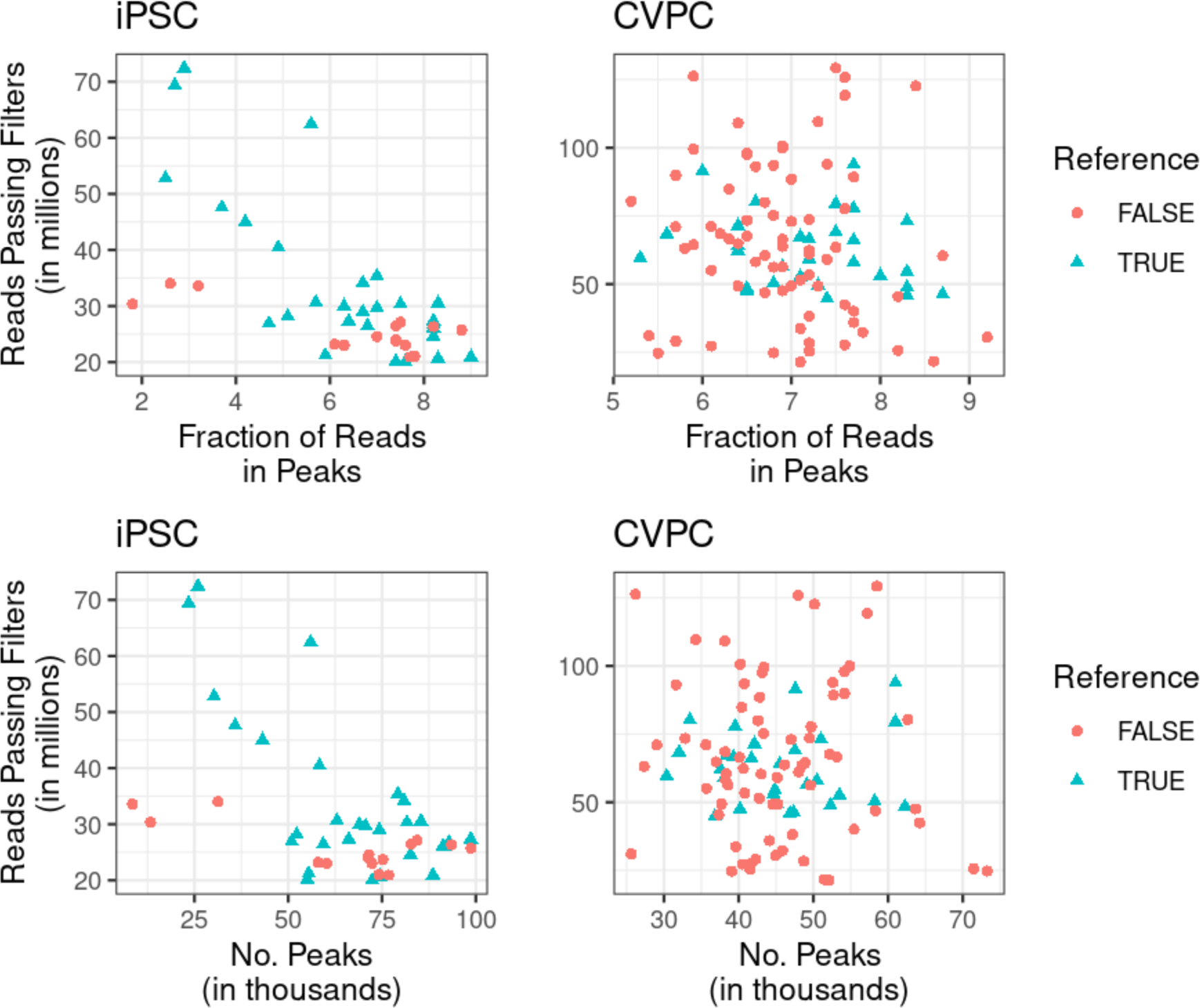
H3K27ac ChIP-seq Quality Control and Reference Sample Selection. Scatter plots displaying the quality of the H3K27ac ChIP-seq samples from iPSCs (n = 43) and CVPCs (n =101). The top row shows the number of reads passing filters of the samples plotted against the fraction of reads in narrow peaks (FRiP), and the bottom row shows the number of reads passing filters against the number of peaks. We selected reference samples (blue triangles) based on QC metrics (see Methods) from unrelated individuals for each tissue (iPSC n = 28; and CVPC n = 29) to establish a set of consensus peaks for the quantification of chromatin accessibility across the respective tissues.

**Figure S10.**
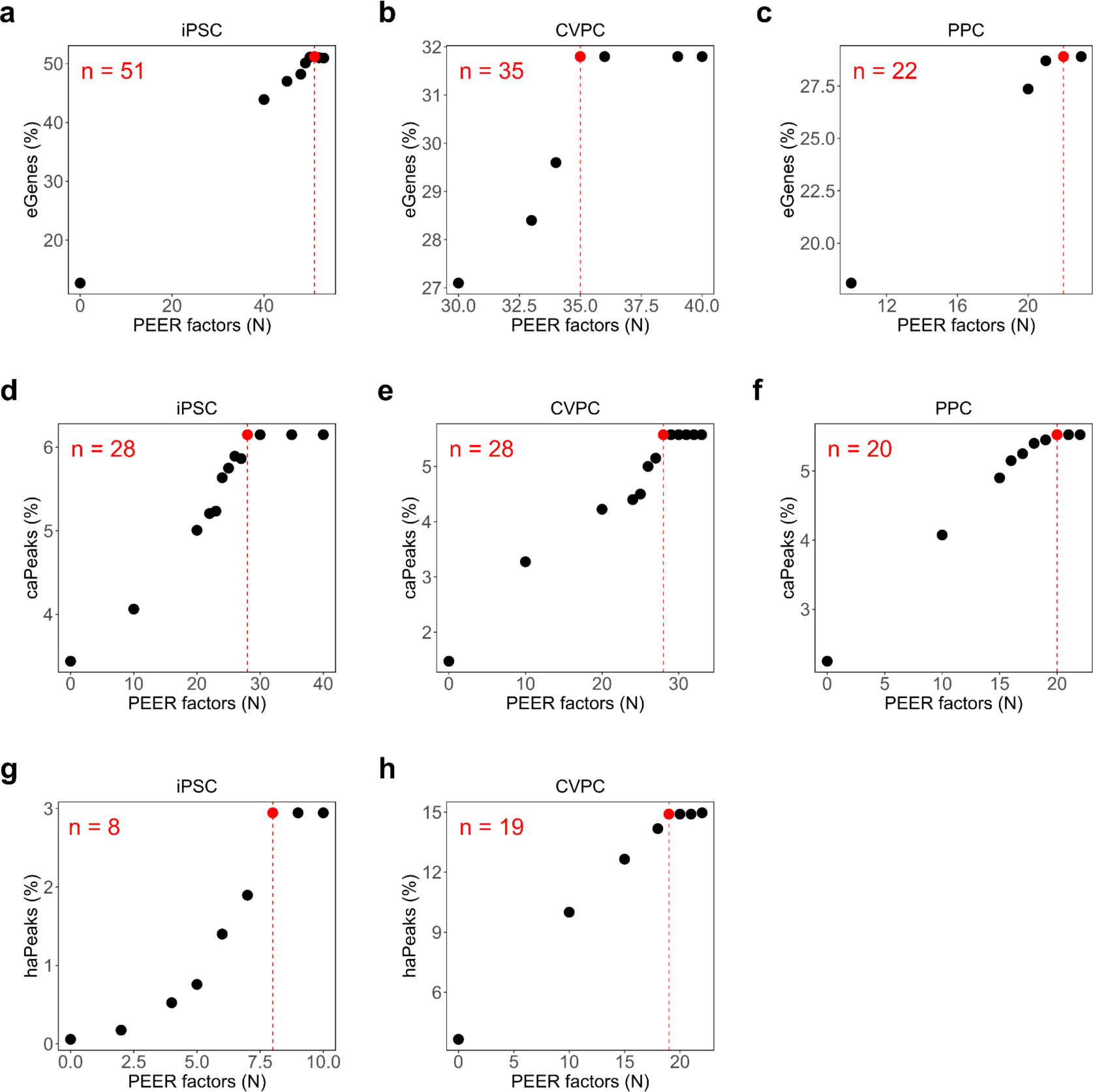
PEER Factor Optimization. Point plots showing the percentage of eGenes/caPeaks/haPeaks that were discovered with varying numbers of PEER factors as covariates. The top row shows PEER optimization results for eQTLs, the middle row shows results for caQTLs, and the bottom row shows results for haQTLs. Red indicates the least number of PEER factors that resulted in maximum eGene/caPeak/haPeak discovery.

**Figure S11.**
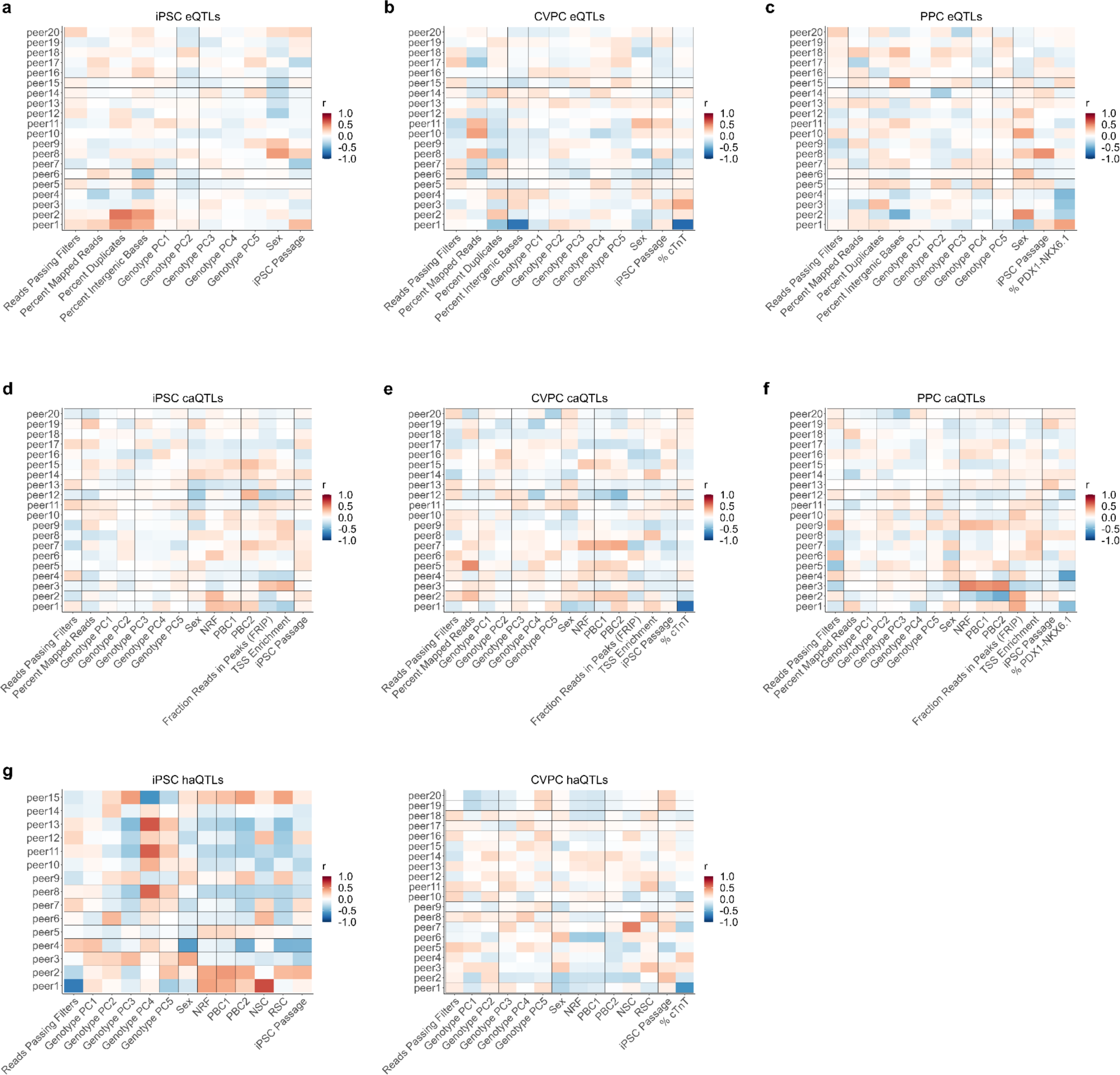
PEER Factor Correlation with Known Covariates. Heatmaps showing the Pearson correlation (r) between PEER factors and the known covariates for each tissue type. First row shows the correlation with PEER factors calculated for eQTLs. Second row shows the correlation with PEER factors calculated for caQTLs. Third row shows the correlation with PEER factors calculated for haQTLs. “Reads passing filters” is the number of reads passing filters. “Genotype PC1-5” are the genotype principal components capturing global ancestry. “Percent Duplicates” is the percentage of duplicate reads. “Percent Intergenic Bases” is the percentage of bases that mapped to intergenic regions. “iPSC Passage’’ is the passage of the iPSCs before CVPC or PPC differentiation. For iPSCs, this indicates the passage of iPSCs upon cell harvest. For CVPCs, “% cTnT” is the percentage of cardiac troponin detected by flow cytometry. For PPCS, “% PDX1-NKX6.1” is the percentage of double-positive PDX1^+^NKX6-1^+^ cells detected by flow cytometry. “NRF” is the non-redundant fraction of reads (i.e., fraction of distinct uniquely mapping reads). “PBC1” is the PCR bottleneck coefficient 1. “PBC2” is the PCR bottleneck coefficient 2. “NSC” is the normalized strand cross-correlation coefficient. “RSC” is the relative strand cross-correlation coefficient. These results show that the top 1-3 PEER factors were correlated with sequencing quality, differentiation efficiency and sex.

**Figure S12.**
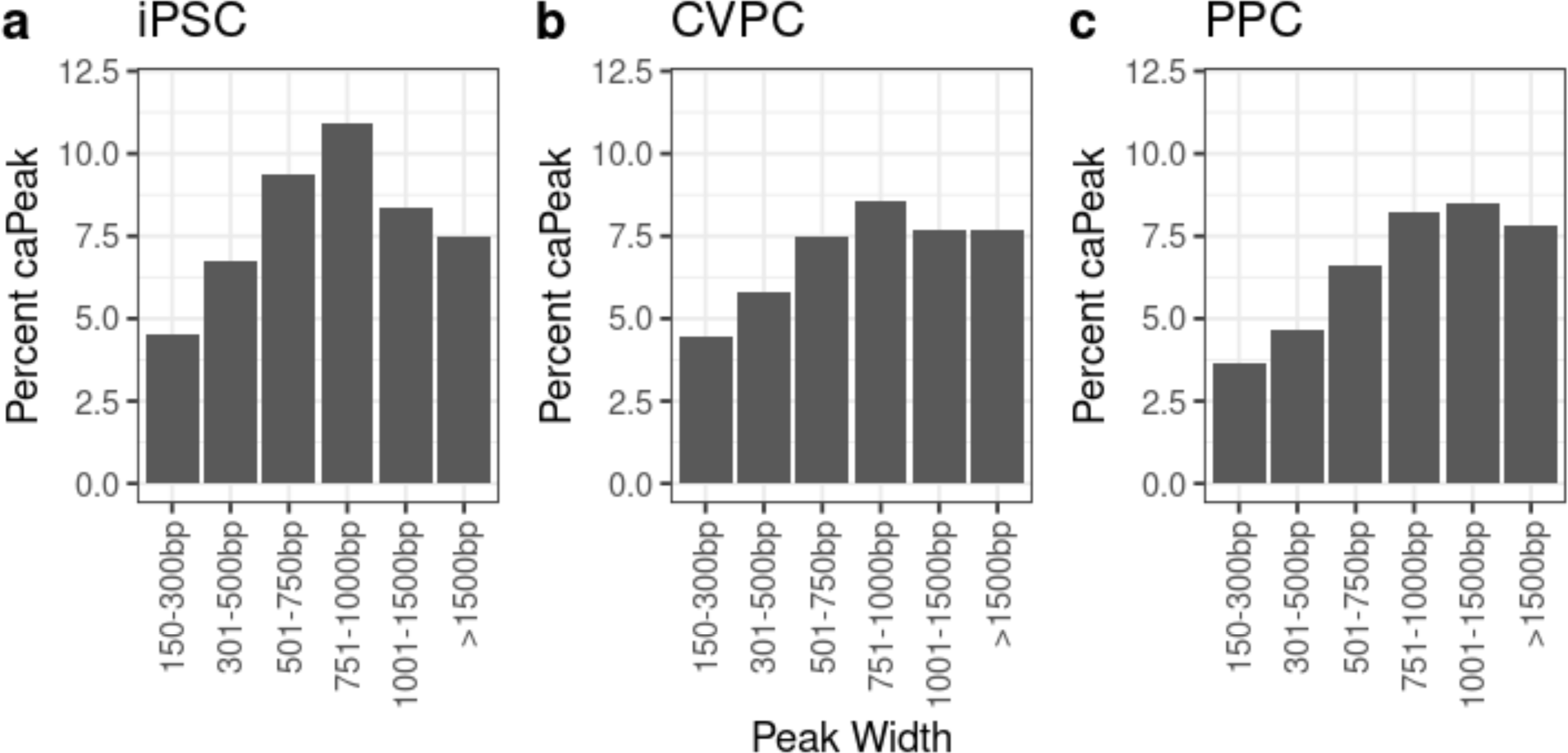
caQTL Discovery Rate by Peak Width. Bar plot showing the caQTL discovery rate by ATAC-seq peak width for iPSCs, CVPCs, and PPCs. caQTL discovery rates were associated with the ATAC-seq peak width. About 8-10% of peaks with widths between 750-1000 bp had caQTLs, while only 3-4% of shorter peaks (150-300 bp) had caQTLs.

## Supplemental Table Legends

**Table S1: Subject Metadata**

This table reports the metadata for the 221 iPSCORE individuals with data used in this study. We provide: **iPSCORE ID:** subject identifier containing the family ID and the individual ID**, Family ID:** iPSCORE family ID**, Sex:** sex**, Age at enrollment:** age of the subject at the time of enrollment**, Most Similar Super Population:** the 1000 Genomes super population that best describes the subject, **Subject UUID:** the universally unique identifier (UUID) for the subject**, WGS UUID:** the UUID for the whole genome sequence corresponding to the subject, and **GPC1-5:** the global genotype principle components (PCs) calculated using the WGS.

**Table S2: RNA Metadata**

This table reports the metadata for the 505 iPSCORE RNA-seq samples used in this study. We provide: **RNA UUID:** the UUID corresponding to the RNA-seq sample, **WGS UUID:** the UUID for the whole genome sequence corresponding to the subject, **Subject UUID:** the UUID corresponding to the subject, **Tissue:** the corresponding tissue, **Clone:** the iPSC clone of the corresponding sample**, Passage:** the passage of the corresponding sample, **iPSC Line ID:** the identifier for the iPSC line (subject iPSCORE ID, and clone) from which the CVPCs and PPCs were derived, **Unique Differentiation ID (UDID):** the label for each differentiation that matches paired molecular data (data from RNA-seq, ATAC-seq, and H3K27ac), **Number of Reads Passing Filters**, **Number of Mapped Reads**, **Number of Properly Paired Reads**, **Duplicates**, **Percent Intergenic Bases**, **Percent mRNA bases**, **Pi hat:** the identity by descent estimate for the matching subject, and **peer1-60:** the PEER factors calculated for each sample. See Figure S8 for the number of PEER factors used for each tissue.

**Table S3: ATAC Metadata**

This table reports the metadata for the 391 iPSCORE ATAC-seq samples used in this study. We provide: **ATAC UUID:** the UUID corresponding to the ATAC-seq sample, **WGS UUID:** the UUID for the whole genome sequence corresponding to the subject, **Subject UUID:** the UUID corresponding to the subject, **Tissue**, **Clone**, **Passage**, **iPSC Line ID:** the identifier for the iPSC line (subject iPSCORE ID, and clone) from which the CVPCs and PPCs were derived, **Unique Differentiation ID (UDID):** the label for each differentiation that matches paired molecular data (data from RNA-seq, ATAC-seq and H3K27ac), **Reference:** a TRUE/FALSE annotation indicating whether the sample was used to establish the consensus ATAC-seq peaks for the corresponding tissue, QC metrics, including **Total Number of Reads**, **Number of Properly Paired Reads**, **Number of Mapped Reads**, **Number of Mitochondrial Reads**, **Non-Redundant Fraction (NRF)**, **PCR Bottlenecking Coefficient 1 (PBC1)**, **PCR Bottlenecking**, **Coefficient 2 (PBC2)**, **Fraction of Reads in Peaks (FRiP)**, and **peer1-30:** the PEER factors calculated for each sample. See Figure S8 for the number of PEER factors used for each tissue.

**Table S4: ChIP Metadata**

This table reports the metadata for the 144 iPSCORE H3K27ac ChIP-seq samples used in this study. We provide: **ChIP UUID:** the UUID corresponding to the ATAC-seq sample, **WGS UUID:** the UUID for the whole genome sequence corresponding to the subject, **Subject UUID**, **Tissue**, **Clone**, **Passage**, **iPSC Line ID:** the identifier for the iPSC line (subject iPSCORE ID, and clone) from which the CVPCs and PPCs were derived, **Unique Differentiation ID (UDID):** the label for each differentiation that matches paired molecular data (data from RNA-seq, ATAC-seq and H3K27ac), **Reference:** a TRUE/FALSE annotation indicating whether the sample was used to establish the consensus ChIP-seq peaks for the corresponding tissue, QC metrics, including, **Number of Reads Passing Filters**, **Number of Mapped Reads**, **Number of Properly Paired Reads**, **Non-Redundant Fraction (NRF)**, **PCR Bottlenecking Coefficient 1 (PBC1)**, **PCR Bottlenecking Coefficient 2 (PBC2)**, **Normalized Strand Cross-correlation coefficient (NSC)**, **Relative strand cross-correlation coefficient (RSC)**, **Fraction of Reads in Peaks (FRiP)**, **Pi Hat:** the identity by descent estimate for the matching subject, and **peer1-20:** the PEER factors calculated for each sample. See Figure S8 for the number of PEER factors used for each tissue.

**Table S5: Enrichment of TFBS**

This table reports the results for the transcription factor binding site enrichment in tissue-specific or shared ATAC-seq peaks. We report: **Motif ID:** the 746 JASPAR and 401 HOCOMOCO v11 motif IDs, **TF Name:** the abbreviated Motif ID name, **AltID:** the Gencode gene name for the TF, **Database:** JASPAR or HOCOMOCO, **Tissue:** the tissue of the specific ATAC-seq peaks enriched in the analysis (iPSC, CVPC, PPC), **Type:** indicates if the enrichment corresponds to “specific” or “shared” peaks, and the results from the Fisher’s Exact tests, including **Odds Ratio**, **the Lower and Upper 95% Confidence Intervals**, nominal and adjusted **P-values.**

**Table S6: Lead variants for all eQTLs**

This table reports the lead variant for each eQTL discovered in the three tissues. We provide: **Tissue**: tissue, **Element ID**: element identifier, **Condition**: discovery order of the QTL where condition = 0 is primary and 1-3 are conditional QTLs, **Element Cond**: element identifier and conditional, **Element Name**: gene name, **Element Chom**: chromosome of element, **Element Start**: start position of element, **Element End**: end position of element, **SNP ID:** the identifier for the lead variant, designated as “VAR_[chromosome]_[position]_[reference allele]_[alternate allele]”, **SNP Chrom:** chromosome of the lead variant, **SNP Pos**: base pair position of the lead variant, **Ref**: reference allele of the lead variant, **Alt**: alternate allele of the lead variant, **RSID**: RSID of the lead variant (if available), **AF**: allele frequency of the lead variant, **Beta**: beta value for the association of the lead variant on gene expression, **SE**: standard error for the association of the lead variant on gene expression, **P-value:** nominal p-value for the lead variant association with gene expression, **FDR**: FDR-corrected p-value calculated by eigenMT, **Tests**: number of independent variants used for eigenMT p-value correction, **Q-value**: q-value from Benjamini-Hochberg correction of FDR, **Significant**: True/False indicating whether the eQTL was significant or not with q-value threshold < 0.05, **Cluster ID**: cluster identifier given to each QTL module and singleton (only applies for primary QTLs that were significant, q-value < 0.05), **Membership**: an annotation for whether the QTL is in a “Module” or is a “Singleton” (only applies for primary QTLs that were significant, q-value < 0.05), **EDev-unique:** True/False indicating whether the eQTL was EDev-unique or not based on LD with adult QTLs (only applies for primary QTLs that were significant, q-value < 0.05). Full summary statistics are available on Figshare as a tar-zipped directory containing text files for each expressed gene tested.

**Table S7: Lead variants for all caQTLs**

This table reports the lead variant for each caQTL discovered in the three tissues. We provide: **Tissue**: tissue, **Element ID**: element identifier, **Condition**: discovery order of the QTL where condition = 0 is primary and 1-3 are conditional QTLs, **Element Cond**: element identifier and conditional, **Element Name**: element name (same as Element_ID), **Element Chrom**: chromosome of element, **Element Start**: start position of element, **Element End**: end position of element, **SNP ID:** the identifier for the lead variant, designated as “VAR_[chromosome]_[position]_[reference allele]_[alternate allele]”, **SNP Chrom:** chromosome of the lead variant, **SNP Pos**: base pair position of the lead variant, **Ref**: reference allele of the lead variant, **Alt**: alternate allele of the lead variant, **RSID**: RSID of the lead variant (if available), **AF**: allele frequency of the lead variant, **Beta**: beta value for the association of the lead variant on gene expression, **SE**: standard error for the association of the lead variant on gene expression, **P-value:** nominal p-value for the lead variant association with gene expression, **FDR**: FDR-corrected p-value calculated by eigenMT, **Tests**: number of independent variants used for eigenMT p-value correction, **Q-value**: q-value from Benjamini-Hochberg correction of FDR, **Significant**: True/False indicating whether the caQTL was significant or not with q-value threshold < 0.05, **Cluster ID**: cluster identifier given to each QTL module and singleton (only applies for primary QTLs that were significant, q-value < 0.05), **Membership**: an annotation for whether the QTL is in a “Module” or is a “Singleton” (only applies for primary QTLs that were significant, q-value < 0.05), **EDev-unique:** True/False indicating whether the caQTL was EDev-unique or not based on LD with adult QTLs (only applies for primary QTLs that were significant, q-value < 0.05). Full summary statistics are available on Figshare as a tar-zipped directory containing text files for each expressed gene tested.

**Table S8: Lead variants for all haQTLs**

This table reports the lead variant for each haQTL discovered in iPSC and CVPC. We provide: **Tissue**: tissue, **Element ID**: element identifier, **Condition**: discovery order of the QTL where condition = 0 is primary and 1-3 are conditional QTLs, **Element Cond**: element identifier and conditional, **Element Name**: element name (same as Element_ID), **Element Chom**: chromosome of element, **Element Start**: start position of element, **Element End**: end position of element, **SNP ID:** the identifier for the lead variant, designated as “VAR_[chromosome]_[position]_[reference allele]_[alternate allele]”, **SNP Chrom:** chromosome of the lead variant, **SNP Pos**: base pair position of the lead variant, **Ref**: reference allele of the lead variant, **Alt**: alternate allele of the lead variant, **RSID**: RSID of the lead variant (if available), **AF**: allele frequency of the lead variant, **Beta**: beta value for the association of the lead variant on gene expression, **SE**: standard error for the association of the lead variant on gene expression, **P-value:** nominal p-value for the lead variant association with gene expression, **FDR**: FDR-corrected p-value calculated by eigenMT, **Tests**: number of independent variants used for eigenMT p-value correction, **Q-value**: q-value from Benjamini-Hochberg correction of FDR, **Significant**: True/False indicating whether the haQTL was significant or not with q-value threshold < 0.05, **Cluster ID**: cluster identifier given to each QTL module and singleton (only applies for primary QTLs that were significant, q-value < 0.05), **Membership**: an annotation for whether the QTL is in a “Module” or is a “Singleton” (only applies for primary QTLs that were significant, q-value < 0.05), **EDev-unique:** True/False indicating whether the haQTL was EDev-unique or not based on LD with adult QTLs (only applies for primary QTLs that were significant, q-value < 0.05). Full summary statistics are available on Figshare as a tar-zipped directory containing text files for each expressed gene tested.

**Table S9: Colocalization between QTLs**

This table reports the subset of results of the intra-tissue pairwise primary QTL colocalization results for all three tissues. The full table with all 1,198,476 intra-tissue pairwise QTL colocalizations is uploaded to Figshare. We provide: **Tissue:** tissue, **Element1 ID, Element2 ID:** the names of the tested qElements, **PP.H4**: posterior probability for H4 being true (both traits have a genetic association and share the same causal variant; output from coloc), **Top SNP ID:** the variant with the highest posterior probability of being causal under the H4 model, **Top SNP Posterior Probability:** causal posterior probability of the top SNP, and **QTL1 Top SNP P-value, QTL2 Top SNP P-value:** the nominal p-values of the top SNP obtained from the QTL analyses. We considered two elements to have a shared QTL signal if the PP.H4 ≥ 0.8 and the top SNP must have a nominal p-value < 5×10^−5^ in both QTLs and a posterior probability ≥ 1%.

**Table S10: QTL Modules and Singletons**

The table reports QTL module membership and singleton QTLs. We provide: **Tissue:** tissue, **Cluster ID:** the identifier for the QTL module or singleton, **Element ID:** the identifier of qElement, **Molecular QTL Type:** an annotation for eQTL, caQTL, or haQTL, **Membership:** an annotation for whether the QTL is in a “Module” or is a “Singleton”, **No. Colocalizations:** the number of times the QTL colocalizes with another QTL, and **No. QTLs in Module:** the number of QTLs in the corresponding QTL module.

**Table S11: QTL Colocalization with GWAS**

Sheet 1 (Manifest): This table provides information about the GWAS summary statistics used in this study. We provide: **Trait Description**: description of the GWAS trait, **Trait ID**: trait ID, **Download Link**: link where the summary statistics were downloaded, **Publication**: the study where the summary statistics originated.

Sheet 2 (Results): This table reports colocalization results between primary QTLs and GWAS variants. We provide: **Tissue**: tissue, **Trait Description**: description of trait, **Trait ID**: trait ID, **Cluster ID**: the identifier for the QTL module or singleton, **QTL Combination**: all QTL types that are associated with the QTL module or singleton (e.g., “haQTL-eQTL” indicates that the QTL module has at least one haQTL and at least one eQTL), **Membership:** label indicating whether the QTL is in a module or a singleton, **Element ID**: element identifier, **Element Name**: element name, **Element Chrom**: chromosome of element, **Element Start**: start position of element, **Element End**: end position of element, **Nsnps**: number of variants used in the colocalization (output from coloc), **PP.H0**: posterior probability for H0 being true (neither trait has a genetic association in the region; output from coloc), **PP.H1**: posterior probability for H1 being true (only trait 1 has a genetic association in the region; in this study, trait 1 = QTL; output from coloc), **PP.H2**: posterior probability for H2 being true (only trait 2 has a genetic association in the region; in this study, trait 2 = GWAS; output from coloc), **PP.H3**: posterior probability for H3 being true (both traits have a genetic association but have distinct causal variants; output from coloc), **PP.H4**: posterior probability for H4 being true (both traits have a genetic association and share the same causal variant; output from coloc), **Max Hypothesis PP**: the maximum posterior probability among the four colocalization hypotheses, **Likely Colocalization Hypothesis**: the colocalization hypothesis with the highest posterior probability, **Top SNP ID**: the variant with the highest posterior probability of being causal under the H4 model, **Top SNP PP:** causal posterior probability for the top SNP, **Proportion Module Colocalized**: proportion of QTLs in the module that colocalized (for singleton QTLs, this value would either be 0 or 1), **Beta.QTL**: effect size for the top SNP’s association with the QTL, **SE.QTL**: standard error for top SNP’s association with the QTL, **Pvalue.QTL**: p-value for top SNP’s association with the QTL, **Beta.GWAS**: effect size for the top SNP’s association with the GWAS trait, **SE.GWAS**: standard error for top SNP’s association with the GWAS trait, **Pvalue.GWAS:** p-value for top SNP’s association with the GWAS trait, **99Credible Set Size**: number of variants in 99% credible set, **Colocalized**: True/False whether the QTL colocalized with GWAS (see Methods), **Has TFBS**: True/False indicating whether the QTL overlapped a TFBS based on whether the caPeak was predicted to have a TFBS or whether the haPeak overlapped an ATAC-seq that was predicted to have a TFBS, **GWAS Index**: index variant for the GWAS signal that colocalized with the QTL (only applies for entries with Colocalized = True), **GWAS QTL Combination Collapse**: all QTL types that are associated with the GWAS signal in the corresponding tissue, because multiple QTLs can colocalize to the same GWAS locus (only applies for entries with Colocalized = True, see Methods), and **EDev-unique GWAS Locus**: True/False whether the GWAS locus was EDev-unique (i.e. colocalized with only EDev-unique QTLs).

## REFERENCES

1. GTEx Consortium (2020). The GTEx Consortium atlas of genetic regulatory effects across human tissues. Science 369, 1318–1330. 10.1126/science.aaz1776.

2. Nguyen, J.P., Arthur, T.D., Fujita, K., Salgado, B.M., Donovan, M.K.R., iPSCORE Consortium, Matsui, H., Kim, J.H., D’Antonio-Chronowska, A., D’Antonio, M., et al. (2023). eQTL mapping in fetal-like pancreatic progenitor cells reveals early developmental insights into diabetes risk. Nat Commun 14, 6928. 10.1038/s41467-023-42560-4.

3. Mostafavi, H., Spence, J.P., Naqvi, S., and Pritchard, J.K. (2023). Systematic differences in discovery of genetic effects on gene expression and complex traits. Nat Genet 55, 1866– 1875. 10.1038/s41588-023-01529-1.

4. D’Antonio, M., Nguyen, J.P., Arthur, T.D., iPSCORE Consortium, Matsui, H., D’Antonio-Chronowska, A., and Frazer, K.A. (2023). Fine mapping spatiotemporal mechanisms of genetic variants underlying cardiac traits and disease. Nat Commun 14, 1132. 10.1038/s41467-023-36638-2.

5. D’Antonio, M., Nguyen, J.P., Arthur, T.D., Matsui, H., Donovan, M.K.R., D’Antonio-Chronowska, A., and Frazer, K.A. (2022). In heart failure reactivation of RNA-binding proteins is associated with the expression of 1,523 fetal-specific isoforms. PLoS Comput Biol 18, e1009918. 10.1371/journal.pcbi.1009918.

6. DeBoever, C., Li, H., Jakubosky, D., Benaglio, P., Reyna, J., Olson, K.M., Huang, H., Biggs, W., Sandoval, E., D’Antonio, M., et al. (2017). Large-Scale Profiling Reveals the Influence of Genetic Variation on Gene Expression in Human Induced Pluripotent Stem Cells. Cell Stem Cell 20, 533–546.e7. 10.1016/j.stem.2017.03.009.

7. Panopoulos, A.D., D’Antonio, M., Benaglio, P., Williams, R., Hashem, S.I., Schuldt, B.M., DeBoever, C., Arias, A.D., Garcia, M., Nelson, B.C., et al. (2017). iPSCORE: A Resource of 222 iPSC Lines Enabling Functional Characterization of Genetic Variation across a Variety of Cell Types. Stem Cell Reports 8, 1086–1100. 10.1016/j.stemcr.2017.03.012.

8. Panopoulos, A.D., Smith, E.N., Arias, A.D., Shepard, P.J., Hishida, Y., Modesto, V., Diffenderfer, K.E., Conner, C., Biggs, W., Sandoval, E., et al. (2017). Aberrant DNA Methylation in Human iPSCs Associates with MYC-Binding Motifs in a Clone-Specific Manner Independent of Genetics. Cell Stem Cell 20, 505–517.e6. 10.1016/j.stem.2017.03.010.

9. D’Antonio, M., Benaglio, P., Jakubosky, D., Greenwald, W.W., Matsui, H., Donovan, M.K.R., Li, H., Smith, E.N., D’Antonio-Chronowska, A., and Frazer, K.A. (2018). Insights into the Mutational Burden of Human Induced Pluripotent Stem Cells from an Integrative Multi-Omics Approach. Cell Rep 24, 883–894. 10.1016/j.celrep.2018.06.091.

10. Greenwald, W.W., Li, H., Benaglio, P., Jakubosky, D., Matsui, H., Schmitt, A., Selvaraj, S., D’Antonio, M., D’Antonio-Chronowska, A., Smith, E.N., et al. (2019). Subtle changes in chromatin loop contact propensity are associated with differential gene regulation and expression. Nat Commun 10, 1054. 10.1038/s41467-019-08940-5.

11. Benaglio, P., D’Antonio-Chronowska, A., Ma, W., Yang, F., Young Greenwald, W.W., Donovan, M.K.R., DeBoever, C., Li, H., Drees, F., Singhal, S., et al. (2019). Allele-specific NKX2-5 binding underlies multiple genetic associations with human electrocardiographic traits. Nat Genet 51, 1506–1517. 10.1038/s41588-019-0499-3.

12. D’Antonio-Chronowska, A., Donovan, M.K.R., Young Greenwald, W.W., Nguyen, J.P., Fujita, K., Hashem, S., Matsui, H., Soncin, F., Parast, M., Ward, M.C., et al. (2019). Association of Human iPSC Gene Signatures and X Chromosome Dosage with Two Distinct Cardiac Differentiation Trajectories. Stem Cell Reports 13, 924–938. 10.1016/j.stemcr.2019.09.011.

13. D’Antonio, M., Reyna, J., Jakubosky, D., Donovan, M.K., Bonder, M.-J., Matsui, H., Stegle, O., Nariai, N., D’Antonio-Chronowska, A., and Frazer, K.A. (2019). Systematic genetic analysis of the MHC region reveals mechanistic underpinnings of HLA type associations with disease. Elife 8. 10.7554/eLife.48476.

14. Jakubosky, D., D’Antonio, M., Bonder, M.J., Smail, C., Donovan, M.K.R., Young Greenwald, W.W., Matsui, H., i2QTL Consortium, D’Antonio-Chronowska, A., Stegle, O., et al. (2020). Properties of structural variants and short tandem repeats associated with gene expression and complex traits. Nat Commun 11, 2927. 10.1038/s41467-020-16482-4.

15. Jakubosky, D., Smith, E.N., D’Antonio, M., Jan Bonder, M., Young Greenwald, W.W., D’Antonio-Chronowska, A., Matsui, H., i2QTL Consortium, Stegle, O., Montgomery, S.B., et al. (2020). Discovery and quality analysis of a comprehensive set of structural variants and short tandem repeats. Nat Commun 11, 2928. 10.1038/s41467-020-16481-5.

16. Bonder, M.J., Smail, C., Gloudemans, M.J., Frésard, L., Jakubosky, D., D’Antonio, M., Li, X., Ferraro, N.M., Carcamo-Orive, I., Mirauta, B., et al. (2021). Identification of rare and common regulatory variants in pluripotent cells using population-scale transcriptomics. Nat Genet 53, 313–321. 10.1038/s41588-021-00800-7.

17. Arthur, T.D., Nguyen, J.P., D’Antonio-Chronowska, A., Matsui, H., Silva, N.S., Joshua, I.N., iPSCORE Consortium, Luchessi, A.D., Greenwald, W.W.Y., D’Antonio, M., et al. (2024). Complex regulatory networks influence pluripotent cell state transitions in human iPSCs. Nat Commun 15, 1664. 10.1038/s41467-024-45506-6.

18. Buenrostro, J.D., Wu, B., Chang, H.Y., and Greenleaf, W.J. (2015). ATAC-seq: A Method for Assaying Chromatin Accessibility Genome-Wide. Curr Protoc Mol Biol 109, 21.29.1–21.29.9. 10.1002/0471142727.mb2129s109.

19. ENCODE Project Consortium (2012). An integrated encyclopedia of DNA elements in the human genome. Nature 489, 57–74. 10.1038/nature11247.

20. Creyghton, M.P., Cheng, A.W., Welstead, G.G., Kooistra, T., Carey, B.W., Steine, E.J., Hanna, J., Lodato, M.A., Frampton, G.M., Sharp, P.A., et al. (2010). Histone H3K27ac separates active from poised enhancers and predicts developmental state. Proc Natl Acad Sci U S A 107, 21931–21936. 10.1073/pnas.1016071107.

21. Ernst, J., and Kellis, M. (2017). Chromatin-state discovery and genome annotation with ChromHMM. Nat Protoc 12, 2478–2492. 10.1038/nprot.2017.124.

22. Bentsen, M., Goymann, P., Schultheis, H., Klee, K., Petrova, A., Wiegandt, R., Fust, A., Preussner, J., Kuenne, C., Braun, T., et al. (2020). ATAC-seq footprinting unravels kinetics of transcription factor binding during zygotic genome activation. Nat Commun 11, 4267. 10.1038/s41467-020-18035-1.

23. Fornes, O., Castro-Mondragon, J.A., Khan, A., van der Lee, R., Zhang, X., Richmond, P.A., Modi, B.P., Correard, S., Gheorghe, M., Baranašić, D., et al. (2019). JASPAR 2020: update of the open-access database of transcription factor binding profiles. Nucleic Acids Research, gkz1001. 10.1093/nar/gkz1001.

24. Kulakovskiy, I.V., Vorontsov, I.E., Yevshin, I.S., Sharipov, R.N., Fedorova, A.D., Rumynskiy, E.I., Medvedeva, Y.A., Magana-Mora, A., Bajic, V.B., Papatsenko, D.A., et al. (2018). HOCOMOCO: towards a complete collection of transcription factor binding models for human and mouse via large-scale ChIP-Seq analysis. Nucleic Acids Res 46, D252–D259. 10.1093/nar/gkx1106.

25. Rozowsky, J., Gao, J., Borsari, B., Yang, Y.T., Galeev, T., Gürsoy, G., Epstein, C.B., Xiong, K., Xu, J., Li, T., et al. (2023). The EN-TEx resource of multi-tissue personal epigenomes & variant-impact models. Cell 186, 1493–1511.e40. 10.1016/j.cell.2023.02.018.

26. Currin, K.W., Erdos, M.R., Narisu, N., Rai, V., Vadlamudi, S., Perrin, H.J., Idol, J.R., Yan, T., Albanus, R.D., Broadaway, K.A., et al. (2021). Genetic effects on liver chromatin accessibility identify disease regulatory variants. Am J Hum Genet 108, 1169–1189. 10.1016/j.ajhg.2021.05.001.

27. Kumasaka, N., Knights, A.J., and Gaffney, D.J. (2016). Fine-mapping cellular QTLs with RASQUAL and ATAC-seq. Nat Genet 48, 206–213. 10.1038/ng.3467.

28. Alasoo, K., Rodrigues, J., Mukhopadhyay, S., Knights, A.J., Mann, A.L., Kundu, K., HIPSCI Consortium, Hale, C., Dougan, G., and Gaffney, D.J. (2018). Shared genetic effects on chromatin and gene expression indicate a role for enhancer priming in immune response. Nat Genet 50, 424–431. 10.1038/s41588-018-0046-7.

29. King, E.A., Dunbar, F., Davis, J.W., and Degner, J.F. (2021). Estimating colocalization probability from limited summary statistics. BMC Bioinformatics 22, 254. 10.1186/s12859-021-04170-z.

30. Chen, Y., Xiao, D., Zhang, L., Cai, C.-L., Li, B.-Y., and Liu, Y. (2021). The Role of Tbx20 in Cardiovascular Development and Function. Front Cell Dev Biol 9, 638542. 10.3389/fcell.2021.638542.

31. Leeson, C.P., Kattenhorn, M., Morley, R., Lucas, A., and Deanfield, J.E. (2001). Impact of low birth weight and cardiovascular risk factors on endothelial function in early adult life. Circulation 103, 1264–1268. 10.1161/01.cir.103.9.1264.

32. Yu, L.-W., Wang, F., Yang, X.-Y., Sun, S.-N., Zheng, Y.-F., Li, B.-B., Gui, Y.-H., and Wang, H.-Y. (2016). Mild decrease in TBX20 promoter activity is a potentially protective factor against congenital heart defects in the Han Chinese population. Sci Rep 6, 23662. 10.1038/srep23662.

33. Khetan, S., Kursawe, R., Youn, A., Lawlor, N., Jillette, A., Marquez, E.J., Ucar, D., and Stitzel, M.L. (2018). Type 2 Diabetes-Associated Genetic Variants Regulate Chromatin Accessibility in Human Islets. Diabetes 67, 2466–2477. 10.2337/db18-0393.

34. Huang, D., Feng, X., Yang, H., Wang, J., Zhang, W., Fan, X., Dong, X., Chen, K., Yu, Y., Ma, X., et al. (2023). QTLbase2: an enhanced catalog of human quantitative trait loci on extensive molecular phenotypes. Nucleic Acids Res 51, D1122–D1128. 10.1093/nar/gkac1020.

35. Barker, D.J. (1990). The fetal and infant origins of adult disease. BMJ 301, 1111. 10.1136/bmj.301.6761.1111.

36. Vo, T., and Hardy, D.B. (2012). Molecular mechanisms underlying the fetal programming of adult disease. J Cell Commun Signal 6, 139–153. 10.1007/s12079-012-0165-3.

37. Gluckman, P.D., Cutfield, W., Hofman, P., and Hanson, M.A. (2005). The fetal, neonatal, and infant environments-the long-term consequences for disease risk. Early Hum Dev 81, 51–59. 10.1016/j.earlhumdev.2004.10.003.

38. Chen, S., Francioli, L.C., Goodrich, J.K., Collins, R.L., Kanai, M., Wang, Q., Alföldi, J., Watts, N.A., Vittal, C., Gauthier, L.D., et al. (2024). A genomic mutational constraint map using variation in 76,156 human genomes. Nature 625, 92–100. 10.1038/s41586-023-06045-0.

39. Huang, Z., Ruan, H.-B., Xian, L., Chen, W., Jiang, S., Song, A., Wang, Q., Shi, P., Gu, X., and Gao, X. (2014). The stem cell factor/Kit signalling pathway regulates mitochondrial function and energy expenditure. Nat Commun 5, 4282. 10.1038/ncomms5282.

40. Krishnamurthy, M., Ayazi, F., Li, J., Lyttle, A.W., Woods, M., Wu, Y., Yee, S.-P., and Wang, R. (2007). c-Kit in early onset of diabetes: a morphological and functional analysis of pancreatic beta-cells in c-KitW-v mutant mice. Endocrinology 148, 5520–5530. 10.1210/en.2007-0387.

41. Feng, Z.-C., Riopel, M., Popell, A., and Wang, R. (2015). A survival Kit for pancreatic beta cells: stem cell factor and c-Kit receptor tyrosine kinase. Diabetologia 58, 654–665. 10.1007/s00125-015-3504-0.

42. Edwards, S.L., Beesley, J., French, J.D., and Dunning, A.M. (2013). Beyond GWASs: illuminating the dark road from association to function. Am J Hum Genet 93, 779–797. 10.1016/j.ajhg.2013.10.012.

43. Strober, B.J., Elorbany, R., Rhodes, K., Krishnan, N., Tayeb, K., Battle, A., and Gilad, Y. (2019). Dynamic genetic regulation of gene expression during cellular differentiation. Science 364, 1287–1290. 10.1126/science.aaw0040.

44. Timmers, P.R., Mounier, N., Lall, K., Fischer, K., Ning, Z., Feng, X., Bretherick, A.D., Clark, D.W., eQTLGen Consortium, Shen, X., et al. (2019). Genomics of 1 million parent lifespans implicates novel pathways and common diseases and distinguishes survival chances. Elife 8, e39856. 10.7554/eLife.39856.

45. Chiou, J., Geusz, R.J., Okino, M.-L., Han, J.Y., Miller, M., Melton, R., Beebe, E., Benaglio, P., Huang, S., Korgaonkar, K., et al. (2021). Interpreting type 1 diabetes risk with genetics and single-cell epigenomics. Nature 594, 398–402. 10.1038/s41586-021-03552-w.

46. Timmers, P.R.H.J., Wilson, J.F., Joshi, P.K., and Deelen, J. (2020). Multivariate genomic scan implicates novel loci and haem metabolism in human ageing. Nat Commun 11, 3570. 10.1038/s41467-020-17312-3.

47. Müller, F.-J., Schuldt, B.M., Williams, R., Mason, D., Altun, G., Papapetrou, E.P., Danner, S., Goldmann, J.E., Herbst, A., Schmidt, N.O., et al. (2011). A bioinformatic assay for pluripotency in human cells. Nat Methods 8, 315–317. 10.1038/nmeth.1580.

48. D’Antonio-Chronowska, A., D’Antonio, M., and Frazer, K.A. (2020). In vitro Differentiation of Human iPSC-derived Cardiovascular Progenitor Cells (iPSC-CVPCs). Bio Protoc 10, e3755. 10.21769/BioProtoc.3755.

49. Lian, X., Zhang, J., Azarin, S.M., Zhu, K., Hazeltine, L.B., Bao, X., Hsiao, C., Kamp, T.J., and Palecek, S.P. (2013). Directed cardiomyocyte differentiation from human pluripotent stem cells by modulating Wnt/β-catenin signaling under fully defined conditions. Nat Protoc 8, 162–175. 10.1038/nprot.2012.150.

50. Fisher, D.J., Heymann, M.A., and Rudolph, A.M. (1981). Myocardial consumption of oxygen and carbohydrates in newborn sheep. Pediatr Res 15, 843–846. 10.1203/00006450-198105000-00003.

51. Werner, J.C., and Sicard, R.E. (1987). Lactate metabolism of isolated, perfused fetal, and newborn pig hearts. Pediatr Res 22, 552–556. 10.1203/00006450-198711000-00016.

52. Tohyama, S., Hattori, F., Sano, M., Hishiki, T., Nagahata, Y., Matsuura, T., Hashimoto, H., Suzuki, T., Yamashita, H., Satoh, Y., et al. (2013). Distinct metabolic flow enables large-scale purification of mouse and human pluripotent stem cell-derived cardiomyocytes. Cell Stem Cell 12, 127–137. 10.1016/j.stem.2012.09.013.

53. Byrska-Bishop, M., Evani, U.S., Zhao, X., Basile, A.O., Abel, H.J., Regier, A.A., Corvelo, A., Clarke, W.E., Musunuri, R., Nagulapalli, K., et al. (2022). High-coverage whole-genome sequencing of the expanded 1000 Genomes Project cohort including 602 trios. Cell 185, 3426–3440.e19. 10.1016/j.cell.2022.08.004.

54. Zheng-Bradley, X., Streeter, I., Fairley, S., Richardson, D., Clarke, L., Flicek, P., and 1000 Genomes Project Consortium (2017). Alignment of 1000 Genomes Project reads to reference assembly GRCh38. Gigascience 6, 1–8. 10.1093/gigascience/gix038.

55. Lowy-Gallego, E., Fairley, S., Zheng-Bradley, X., Ruffier, M., Clarke, L., Flicek, P., and 1000 Genomes Project Consortium (2019). Variant calling on the GRCh38 assembly with the data from phase three of the 1000 Genomes Project. Wellcome Open Res 4, 50. 10.12688/wellcomeopenres.15126.2.

56. Zhao, H., Sun, Z., Wang, J., Huang, H., Kocher, J.-P., and Wang, L. (2014). CrossMap: a versatile tool for coordinate conversion between genome assemblies. Bioinformatics 30, 1006–1007. 10.1093/bioinformatics/btt730.

57. Frankish, A., Diekhans, M., Jungreis, I., Lagarde, J., Loveland, J.E., Mudge, J.M., Sisu, C., Wright, J.C., Armstrong, J., Barnes, I., et al. (2021). GENCODE 2021. Nucleic Acids Res 49, D916–D923. 10.1093/nar/gkaa1087.

58. Frankish, A., Diekhans, M., Ferreira, A.-M., Johnson, R., Jungreis, I., Loveland, J., Mudge, J.M., Sisu, C., Wright, J., Armstrong, J., et al. (2019). GENCODE reference annotation for the human and mouse genomes. Nucleic Acids Res 47, D766–D773. 10.1093/nar/gky955.

59. Danecek, P., Bonfield, J.K., Liddle, J., Marshall, J., Ohan, V., Pollard, M.O., Whitwham, A., Keane, T., McCarthy, S.A., Davies, R.M., et al. (2021). Twelve years of SAMtools and BCFtools. Gigascience 10, giab008. 10.1093/gigascience/giab008.

60. Li, B., and Dewey, C.N. (2011). RSEM: accurate transcript quantification from RNA-Seq data with or without a reference genome. BMC Bioinformatics 12, 323. 10.1186/1471-2105-12-323.

61. Robinson, M.D., McCarthy, D.J., and Smyth, G.K. (2010). edgeR: a Bioconductor package for differential expression analysis of digital gene expression data. Bioinformatics 26, 139–140. 10.1093/bioinformatics/btp616.

62. Martin, M. (2011). Cutadapt removes adapter sequences from high-throughput sequencing reads. EMBnet j. 17, 10. 10.14806/ej.17.1.200.

63. Li, H., and Durbin, R. (2009). Fast and accurate short read alignment with Burrows-Wheeler transform. Bioinformatics 25, 1754–1760. 10.1093/bioinformatics/btp324.

64. Li, H. (2013). Aligning sequence reads, clone sequences and assembly contigs with BWA-MEM. Preprint at arXiv.

65. Zhang, Y., Liu, T., Meyer, C.A., Eeckhoute, J., Johnson, D.S., Bernstein, B.E., Nusbaum, C., Myers, R.M., Brown, M., Li, W., et al. (2008). Model-based analysis of ChIP-Seq (MACS). Genome Biol 9, R137. 10.1186/gb-2008-9-9-r137.

66. Liao, Y., Smyth, G.K., and Shi, W. (2014). featureCounts: an efficient general purpose program for assigning sequence reads to genomic features. Bioinformatics 30, 923–930. 10.1093/bioinformatics/btt656.

67. Purcell, S., Neale, B., Todd-Brown, K., Thomas, L., Ferreira, M.A.R., Bender, D., Maller, J., Sklar, P., de Bakker, P.I.W., Daly, M.J., et al. (2007). PLINK: a tool set for whole-genome association and population-based linkage analyses. Am J Hum Genet 81, 559–575. 10.1086/519795.

68. Stegle, O., Parts, L., Piipari, M., Winn, J., and Durbin, R. (2012). Using probabilistic estimation of expression residuals (PEER) to obtain increased power and interpretability of gene expression analyses. Nat Protoc 7, 500–507. 10.1038/nprot.2011.457.

69. Dilthey, A.T. (2021). State-of-the-art genome inference in the human MHC. Int J Biochem Cell Biol 131, 105882. 10.1016/j.biocel.2020.105882.

70. Huang, Q.Q., Ritchie, S.C., Brozynska, M., and Inouye, M. (2018). Power, false discovery rate and Winner’s Curse in eQTL studies. Nucleic Acids Res 46, e133. 10.1093/nar/gky780.

71. Davis, J.R., Fresard, L., Knowles, D.A., Pala, M., Bustamante, C.D., Battle, A., and Montgomery, S.B. (2016). An Efficient Multiple-Testing Adjustment for eQTL Studies that Accounts for Linkage Disequilibrium between Variants. Am J Hum Genet 98, 216–224. 10.1016/j.ajhg.2015.11.021.

72. Giambartolomei, C., Vukcevic, D., Schadt, E.E., Franke, L., Hingorani, A.D., Wallace, C., and Plagnol, V. (2014). Bayesian test for colocalisation between pairs of genetic association studies using summary statistics. PLoS Genet 10, e1004383. 10.1371/journal.pgen.1004383.

73. Csardi, Gabor, N., Tamas (2006). The igraph software package for complex network research. InterJounal Complex Systems, 1695.

74. Csárdi, G., Nepusz, T., Müller, K., Horvát, S., Traag, V., Zanini, F., and Noom, D. (2024). igraph for R: R interface of the igraph library for graph theory and network analysis. Version v2.0.1.1 (Zenodo). 10.5281/ZENODO.7682609 10.5281/ZENODO.7682609.

75. Bradfield, J.P., Vogelezang, S., Felix, J.F., Chesi, A., Helgeland, Ø., Horikoshi, M., Karhunen, V., Lowry, E., Cousminer, D.L., Ahluwalia, T.S., et al. (2019). A trans-ancestral meta-analysis of genome-wide association studies reveals loci associated with childhood obesity. Hum Mol Genet 28, 3327–3338. 10.1093/hmg/ddz161.

76. Horikoshi, M., Beaumont, R.N., Day, F.R., Warrington, N.M., Kooijman, M.N., Fernandez-Tajes, J., Feenstra, B., van Zuydam, N.R., Gaulton, K.J., Grarup, N., et al. (2016). Genome-wide associations for birth weight and correlations with adult disease. Nature 538, 248–252. 10.1038/nature19806.

77. Chen, J., Spracklen, C.N., Marenne, G., Varshney, A., Corbin, L.J., Luan, J., Willems, S.M., Wu, Y., Zhang, X., Horikoshi, M., et al. (2021). The trans-ancestral genomic architecture of glycemic traits. Nat Genet 53, 840–860. 10.1038/s41588-021-00852-9.

78. Mahajan, A., Taliun, D., Thurner, M., Robertson, N.R., Torres, J.M., Rayner, N.W., Payne, A.J., Steinthorsdottir, V., Scott, R.A., Grarup, N., et al. (2018). Fine-mapping type 2 diabetes loci to single-variant resolution using high-density imputation and islet-specific epigenome maps. Nat Genet 50, 1505–1513. 10.1038/s41588-018-0241-6.

79. Timmers, P.R., Mounier, N., Lall, K., Fischer, K., Ning, Z., Feng, X., Bretherick, A.D., Clark, D.W., eQTLGen Consortium, Shen, X., et al. (2019). Genomics of 1 million parent lifespans implicates novel pathways and common diseases and distinguishes survival chances. Elife 8, e39856. 10.7554/eLife.39856.

80. Timmers, P.R.H.J., Wilson, J.F., Joshi, P.K., and Deelen, J. (2020). Multivariate genomic scan implicates novel loci and haem metabolism in human ageing. Nat Commun 11, 3570. 10.1038/s41467-020-17312-3.

81. Giambartolomei, C., Vukcevic, D., Schadt, E.E., Franke, L., Hingorani, A.D., Wallace, C., and Plagnol, V. (2014). Bayesian Test for Colocalisation between Pairs of Genetic Association Studies Using Summary Statistics. PLoS Genet 10, e1004383. 10.1371/journal.pgen.1004383.

82. Viñuela, A., Varshney, A., van de Bunt, M., Prasad, R.B., Asplund, O., Bennett, A., Boehnke, M., Brown, A.A., Erdos, M.R., Fadista, J., et al. (2020). Genetic variant effects on gene expression in human pancreatic islets and their implications for T2D. Nat Commun 11, 4912. 10.1038/s41467-020-18581-8.

83. Bioconductor Package liftOver. ([object Object]). 10.18129/B9.BIOC.LIFTOVER 10.18129/B9.BIOC.LIFTOVER.

